# Molecular and Pathological Characterization of Cervicovaginal Specimens Evince Significant Protection Against HPV Infections, Bacterial Vaginosis, and Epithelial Cell Abnormalities by *Lactobacillus* species

**DOI:** 10.1101/2024.12.30.630800

**Authors:** John Osei Sekyere, Jason Trama, Martin Adelson, Charulata Trikannad, Desiree DiBlasi, Rachel Schuster, Jing Jing Yang, Eli Mordechai

## Abstract

**Background:** Bacterial vaginosis (BV) results in dysbiosis in the vaginal microbiome and facilitates secondary human papilloma virus (HPV) infections, which can cause cervical abnormalities. The effects of *Lactobacilli* on BV-HPV infections and cervicovaginal pathology were investigated.

**Method:** Electronic molecular and cytology laboratory results, and demographic records of 19,105 females screened for BV, HPV, and cervical cancer between August 2021 and April 2023 were collected. After data cleaning and reorganization, 15, 607 sample results were used. The data was subjected to epidemiological, statistical, and machine learning analyses to identify patterns, correlations, associations, and interactive effects of age, State, *Lactobacilli,* BV-associated pathogens, high-risk HPV (hrHPV) genotypes, BV, and cervical cytology.

**Results:** The final dataset comprised of data from anonymized females (n = 15541, 99.57%), males (n = 13, 0.08%) and unknown gender (n = 53, 0.34%) between 14 and 95 years from 32 States and Washington D.C. Most samples were BV positive (n = 8282, 53.07%) and/or NILM (n = 11749) followed by 14.29% ASCUS (n = 2231), 7.14% LSIL (n = 1115), 1.04% RCC (n = 163), 0.53% HSIL (n = 82), 0.39% ASC-H (n = 61), and 0.16% AGC (n = 25). HPV-positive samples had higher BV pathogens and BV-positive diagnosis (between 51.90% and 68.60%) than BV-negative diagnosis. *Lactobacillus gasseri* (79.75%)*, Lactobacillus jensenii* (82.68%)*, Lactobacillus crispatus* (90.91%) were mostly found in BV-negative and NILM specimens. BV and ECA outcomes were significantly associated with age and State. Statistical, correlation, and machine learning models predictively confirmed these findings.

**Conclusion:** *L. crispatus,* followed by *L. gasseri* and *L. jensenii*, significantly maintains a healthy cervicovaginal microbiome by inhibiting harmful bacterial pathogens and high-risk HPV subtypes; *L. iners* have an opposing or neutral effect.

**Lay summary/importance/highlights:** Bacterial vaginosis (BV) and human papillomavirus (HPV) infections mostly occur in sexually active and reproductive-age women, adversely affecting their quality of life and predisposing them to sequelae of reproductive disorders. In this study, the protective effects of *Lactobacilli* against BV and HPV infections to maintain a healthy cervicovaginal microbiome were investigated. *Lactobacillus crispatus, L. gasseri,* and *L. jensenii* were common in clinical specimens that had no or relatively lower concentrations of hrHPV, BV, and cervico-vaginal cell abnormalities and vice versa. This shows how important these *Lactobacilli* are in maintaining a healthy cervix and vagina. Age and BV were found to be strongly associated with the incidence of HPV infections and epithelial cell abnormalities (ECAs). Some States had higher prevalence of BV and ECAs than others. Co-infection with multiple hrHPV and/or hrHPV-BV pathogens segued into negative health effects; some interactions were beneficial for the microbiome, albeit those were few. This makes multiple HPV genotype infections dangerous.

## 1. INTRODUCTION

The normal vaginal microbial flora is comprised of abundant H_2_O_2_-producing *Lactobacillus* sp. and a low abundance of other anaerobic bacteria, which keep the vaginal microbiome acidic (pH < 4.5) ^1^. Under the regulation of circulating estrogen, in a concentration-dependent manner, glycogen is deposited into the vaginal lumen by vaginal and cervical epithelial cells, which is metabolized by *Lactobacillus* sp. into lactic acid ^1,2^. This results in an acidic vaginal pH, which inhibits the proliferation of other microbial species, keeping the diversity of the vaginal microbiome low ^3^. A change in estrogen levels, as observed during menopause, menstruation, puberty, and pregnancy, affects vaginal glycogen levels and consequently, the vaginal pH, *Lactobacillus* sp. abundance, and microbiota composition ^3,4^. Further, the use of broad-spectrum antibiotics, which can deplete *Lactobacillus* sp., also causes vaginal microbiome dysbiosis ^5,6^.

Depletion of *Lactobacillus sp.* and increase in vaginal pH permit other microbial species to proliferate, increasing the microbial diversity and causing sequelae of vulvovaginal candidiasis and bacterial vaginosis (BV). This further predisposes the vagina and cervix to sexually transmitted infections (STIs) such as gonorrhea, human papillomavirus (HPV) and HIV infections, obstetric complications, and cervical cancer ^3,7^. Notably, BV infections have been implicated as a risk factor for cervical HPV infections: BV affects almost 30% of reproductive age women globally and is the most common cause of vaginal discharge ^8^ while HPV infections, which is a major cause of cervical cancer, affects around 26.8-38.4% of women aged 15-59 years ^9,10^ and 31% of men ^11^.

Of the over 200 HPV types, not all are associated with cervical cancer; those that are, are called high-risk HPV (hrHPV) while those that are not, are classified as non-high risk, being associated with anogenital warts ^11^. There are about 12 hrHPV types, including HPV-18, -16, -31, -33, -45, -52, and –58, which are implicated in 99% of cervical cancers. Type 16 causes approximately 50% of cervical cancers while both types 16 and 18 are responsible for 66% of cervical cancers ^12^. The other hrHPV types are associated with 15% of cervical cancers and 11% of HPV-associated cancers. Although most women with hrHPV do not develop cervical cancer, HPV infection can cause cervical (epithelial) cell abnormalities (ECA) ^12^. Based on cytology findings, the cervix’s squamous cell epithelium can be classified as negative for intraepithelial lesion or malignancy (NILM) for those with no ECA while ECA are classified into three major categories: atypical squamous cells of undetermined significance (ASC-US), atypical squamous cells–cannot exclude HSIL (ASC-H), low-grade squamous intraepithelial lesions (LSIL), high-grade squamous intraepithelial lesions (HSIL), and squamous cell carcinoma (SCC) ^13^.

Several studies have reported the benefits of *Lactobacilli* against BV, HPV infections, and ECAs. Yet, none of these studies have used a larger cohort of datasets comprising of demographics, qPCR, and pathological data to understand the prevalence, age distribution, associations, correlation, interaction effects, and determinants of BV, HPV, and cervical cytology in the presence and absence of *Lactobacilli.* This study retrospectively reviewed previously reported laboratory test results obtained from 15,607 specimens sent from health care providers across the United States to a single laboratory.

## 2.0 Methods

### 2.1 Sampling

This study was conducted using clinical electronic data pulled from the database of Medical Diagnostic Laboratories (MDL), LLC, based in Hamilton Township, New Jersey. The clinical data comprised of the demographic and qPCR records obtained from molecular analyses of vaginal and cervical swab (and urine, anal, and semen from males) specimens that were received at the laboratory between August 1, 2021, and April 5, 2023. The data consisted of (1) the demographics i.e., gender, age, and healthcare provider State, (2) qPCR CT scores and extrapolated DNA concentrations of bacterial species and HPV subtypes, (3) bacterial vaginosis diagnosis, (4) high-risk HPV (hrHPV) results, and (5) pap smear cervical cytology results (Supplemental Table 1). The demographic data were obtained from the presenting physicians’ records uploaded to the MDL requisition website while the molecular and pathological data were obtained from the routine clinical laboratory analysis of clinical specimens.

### 2.2 Microbiology and pathology analyses

The cervical and vaginal specimen were subjected to qPCR testing to determine the presence of *Fannyhessea vaginae, Gardnerella vaginalis,* bacteria-vaginosis associated bacteria (BVAB-2), *Megasphaera sp. Type 1, Megasphaera sp. Type 2, Lactobacillus iners, Lactobacillus gasseri, Lactobacillus jensenii, L. crispatus,* HPV 16 , HPV 18, HPV 31, HPV 33, HPV 35, HPV 39, HPV 45, HPV 51, HPV 52, HPV 56, HPV 58 HPV 59, and HPV 68 using already described methods. Based on the outcome of the qPCR, the specimens were classified as BV positive, BV negative, or transitional BV using the relative abundance concentrations of the bacterial species. The presence of the HPV types was also used to classify the specimen as hrHPV abnormal (or normal if no HPV is present).

The cervical cytology analysis of the samples is undertaken using already described methods to identify the types of cell pathologies found in the cervix and classify them as negative for intraepithelial lesion or malignancy (NILM) or epithelial cell abnormality (ECA). Some specimens classified as NILM cells were further classified as reactive cellular changes (RCC); RCC refers to the appearance of cervical cells when there is inflammation or infection nearby. RCC is a benign cellular change that is associated with inflammation. ECA specimens were further classified as atypical glandular cells (AGC), atypical squamous cells of undetermined significance (ASCUS), atypical squamous cells, cannot exclude high-grade squamous intraepithelial lesion (ASC-H), high-grade squamous intraepithelial lesion (HSIL), Low-grade squamous intraepithelial lesion (LSIL) (Table S1).

### 2.3 Data cleaning and restructuring

The data were arranged into arrays and schema, placing each item into a unique row and column for easier downstream analyses. Texts in the gender, sex, BV Status, hrHPV results, and pap smear results columns were encoded into numbers for easy statistical analysis and machine learning processing. Duplicate entries and specimens with no qPCR results for any of the bacterial species, resulting in an indeterminable BV diagnosis, were deleted.

### 2.4 Data analysis

The data analysis was divided into four steps. (1) Epidemiological analysis encompassing descriptive and basic statistics such as counts, means, distribution, and standard deviations. (2) Association, correlation, and interaction effect analysis, which delved into the relationships between demographics (gender, age, State), incidence and prevalence of the nine bacterial species and HPV subtypes, BV diagnosis, and pap smear/cervical cytology outcomes. (3) Principal component analysis and K-means clustering to determine the variance and distribution of the bacterial species and HPV types across the specimens. (4) Machine learning models to identify important features that determine or impact the incidence of BV and cervical cytology outcomes.

All data analyses were conducted using GraphPad prism 10.2.2 and Python3 (version 3.10.13) on Ubuntu 22.04.2 LTS using the following libraries: SciPy and Statsmodels (for statistics and linear regression), Seaborn (for correlations, heatmaps, and distribution plots), Pandas (for handling data in DataFrame, frequency counts and mean concentrations), Matplotlib (plots and charts generation), Scikit-learn (principal component analysis, PCA, and K-Means clustering, machine learning models), XGBoost (machine learning algorithms), NumPy (numerical calculations, array operations, and statistical analysis).

*P*-values of < 0.05 was defined as significant in all calculations and statistics.

## 3.0 Results

### 3.1 Demographics

We obtained 19, 105 clinical specimen records from the MDL database within the period of study (August 1, 2021, and April 5, 2023); 3,498 specimen data were removed, resulting in a final dataset of 15, 607 specimens from 15607 persons. The specimens were obtained from healthcare providers located in 32 States and the District of Columbia (DC). Most samples were obtained from Texas (n = 2327), Arizona (n = 1790), Illinois (n = 1784), Florida (n = 1660), Louisiana (n = 1517), Michigan (n = 1511), and California (n = 1052). Females were mostly (n = 15541, 99.57%) the source of these specimens, with males (n = 13, 0.08%) and unknown gender (n = 53, 0.34%) being in the minority. The ages of the patients were between 14 and 95 years: most samples were obtained from persons within 20-60 years; the highest number of samples was from age 30 (n = 689) (Dataset 1.1-1.3; Fig. 1).

**Figure 1.**
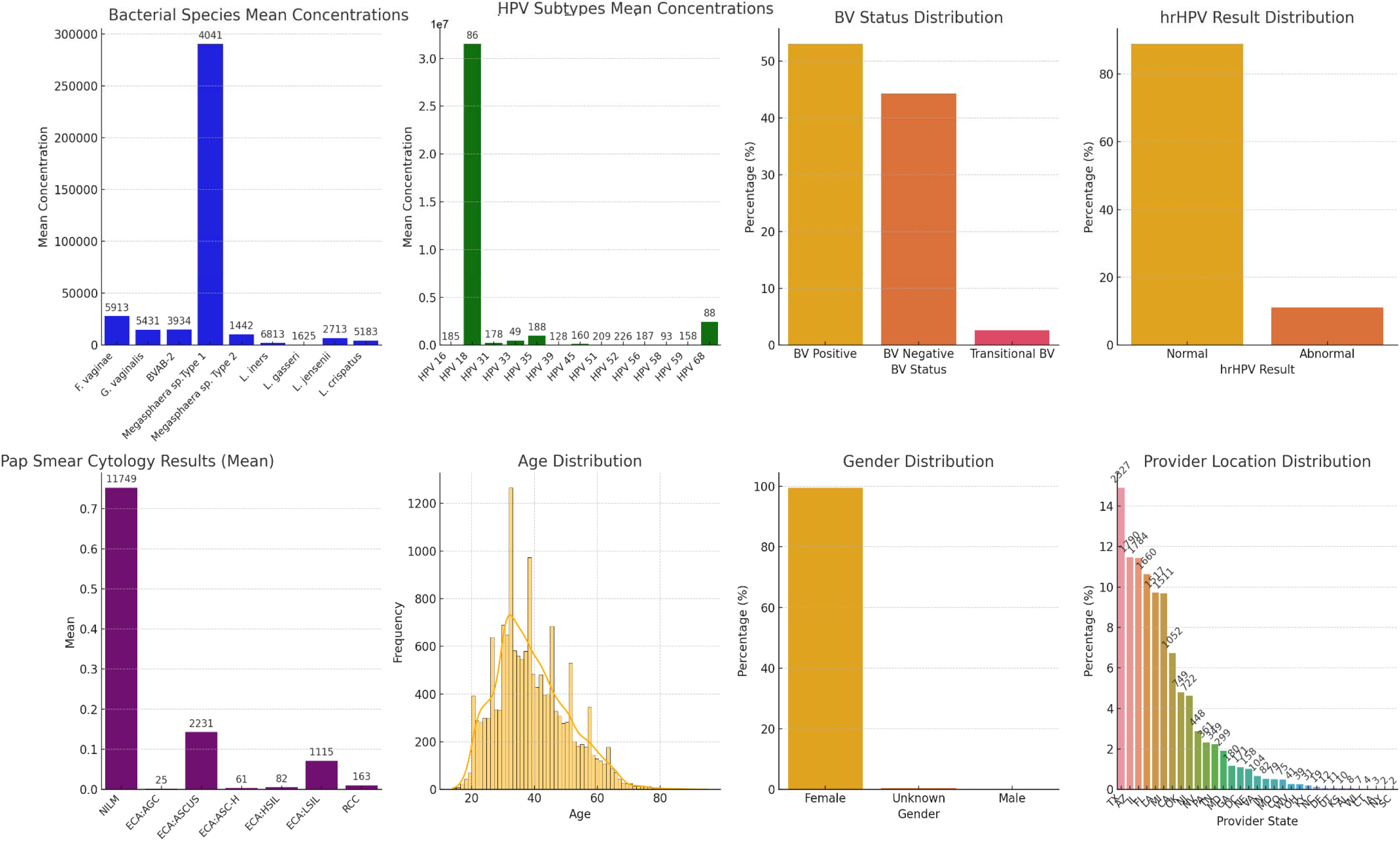
Epidemiological analysis showing the counts, mean concentrations, and distributions of bacterial species, HPV subtypes, BV diagnosis, hrHPV results, cervical cytology results, and age, gender, and provider location distribution. The annotations on the bars are the counts to show the actual prevalence/occurrence in all the samples. ASCUS and LSIL were the most common epithelial cell abnormality diagnoses among the samples while most of the samples were collected from patients aged between 30 and 40.

### 3.2 Molecular (qPCR) and cervical cytology outcomes

Nine bacterial species and 13 HPV subtypes were tested in each sample, resulting in concentrations and Ct scores (Table S1): Megasphaera sp. Type 1 and HPV-18 had the highest mean concentrations among the bacterial species and HPV types, respectively. However, *L. iners* (n = 6813, 43.65%), *F. vaginae* (n = 5913, 37.89%)*, G. vaginalis* (n = 5431, 34.80%), and *L. crispatus* (n = 5183, 33.21%) were most prevalent bacteria among the samples (Dataset 1.4) while HPV-52 (n = 226, 1.45%), HPV-51 (n = 209, 1.34%), HPV-35 (n = 188, 1.20%), HPV-56 (n = 187, 1.20%) and HPV-16 (n = 185, 1.19%) were most prevalent HPV types (Dataset 1.5).

There were more BV positive (n = 8282, 53.07%) diagnosis than BV negative (n = 6919, 44.33%) and transitional BV (n = 406, 2.60%) (Dataset 1.6). Contrarily, 1726 (11.06%) samples/patients were abnormal/positive for hrHPV while 13881 (88.94%) were normal/negative for hrHPV (Dataset 1.7). Further, the Pap smear cervical cytology results had 75.28% NILM (n = 11749) diagnosis followed by 14.29% ASCUS (n = 2231), 7.14% LSIL (n = 1115), 1.04% RCC (n = 163), 0.53% HSIL (n = 82), 0.39% ASC-H (n = 61), and 0.16% AGC (n = 25) (Dataset 1.8).

### 3.2 Significant associations, interactions, and correlations

#### Age associations

There was a significant distribution of age across the different States, BV diagnosis, and hrHPV outcomes. Notably, age per bacteria species and BV status had little variations except for *L. gasseri* that was mostly detected in patients aged between 35 and 50. HPV types 18 and 33 mostly occurred in older populations than the rest while samples from Delaware, Missouri, Colorado, and DC were from older patients than the rest. Further, AGC, HSIL, and RCC were mostly in patients older than the other outcomes. In all, hrHPV-positive samples were in a younger population than normal ones (Fig. S1; Datasets 1.9 – 1.15).

#### Gender associations

Gender was only significantly associated with provider location (States). The ages of the different gender were mostly within the same brackets. It’s notable that all the bacterial species (except *L. gasseri* in males) were also found in specimens from males and unknown gender patients, with 8249, 4, and 29 BV-positive specimens being from females, males, and unknown gender respectively. Comparatively, fewer HPV were found in males (HPV-31 and -39) and unknown gender (HPV-16, -45, -52, and -59). ECA such as ASCUS and AGC were found in males and unknown gender while LSIL was found in unknown gender (Fig. S2 – S4; Datasets 1.16 – 1.23).

#### Provider location (State) associations

Except for the HPV subtypes, there was significant association between the provider’s location (State) and cervical cytology, hrHPV and BV outcomes, and detected bacteria species and HPV subtypes. Notably, there were more BV positive (except in Colorado, California, Delaware, Michigan, Oklahoma, and Tennessee) and hrHPV normal diagnosis in most States, reflecting the distribution of BV-causing bacteria and *Lactobacillus species,* while ASCUS and LSIL were relatively more prevalent in Arizona, Florida, Illinois, Louisiana, Michigan, Oklahoma, and Texas; these States also had substantial presence of almost all the HPV types. The nine bacteria species were commonly detected in relatively higher numbers in samples from States including Arizona, California, DC, Florida, Georgia, Illinois, Louisiana, Maryland, New Jersey, Nevada, Oklahoma, Pennsylvania, Tennessee, and Texas; reflecting the higher prevalence of bacteria species in the specimens than HPV types (Dataset 1.24-1.27; Fig. S5-S8).

#### BV associations

There were substantially higher and significant presence (association) of *F. vaginae* (90.48%)*, G. vaginalis* (86.60%)*, BVAB-2* (96.54%)*, Megasphaera sp. Type 1* (98.42%)*, Megasphaera sp. Type 2* (97.92%), and *L. iners* (65.54%) in BV-positive samples; specifically, 2201 BV-negative samples (32.31%) were *L. iners-*positive compared to 4465-positive *L. iners* (65.54%) in BV-positive samples. Furthermore, HPV-positive samples had higher BV-positive diagnosis (between 51.90% and 68.60%) than BV-negative diagnosis. Contrarily, *L. gasseri* (79.75%)*, L. jensenii* (82.68%)*, L. crispatus* (90.91%) were mostly found in BV-negative specimens (Fig. 2 & S9; Datasets 2.1-2.6).

**Figure 2.**
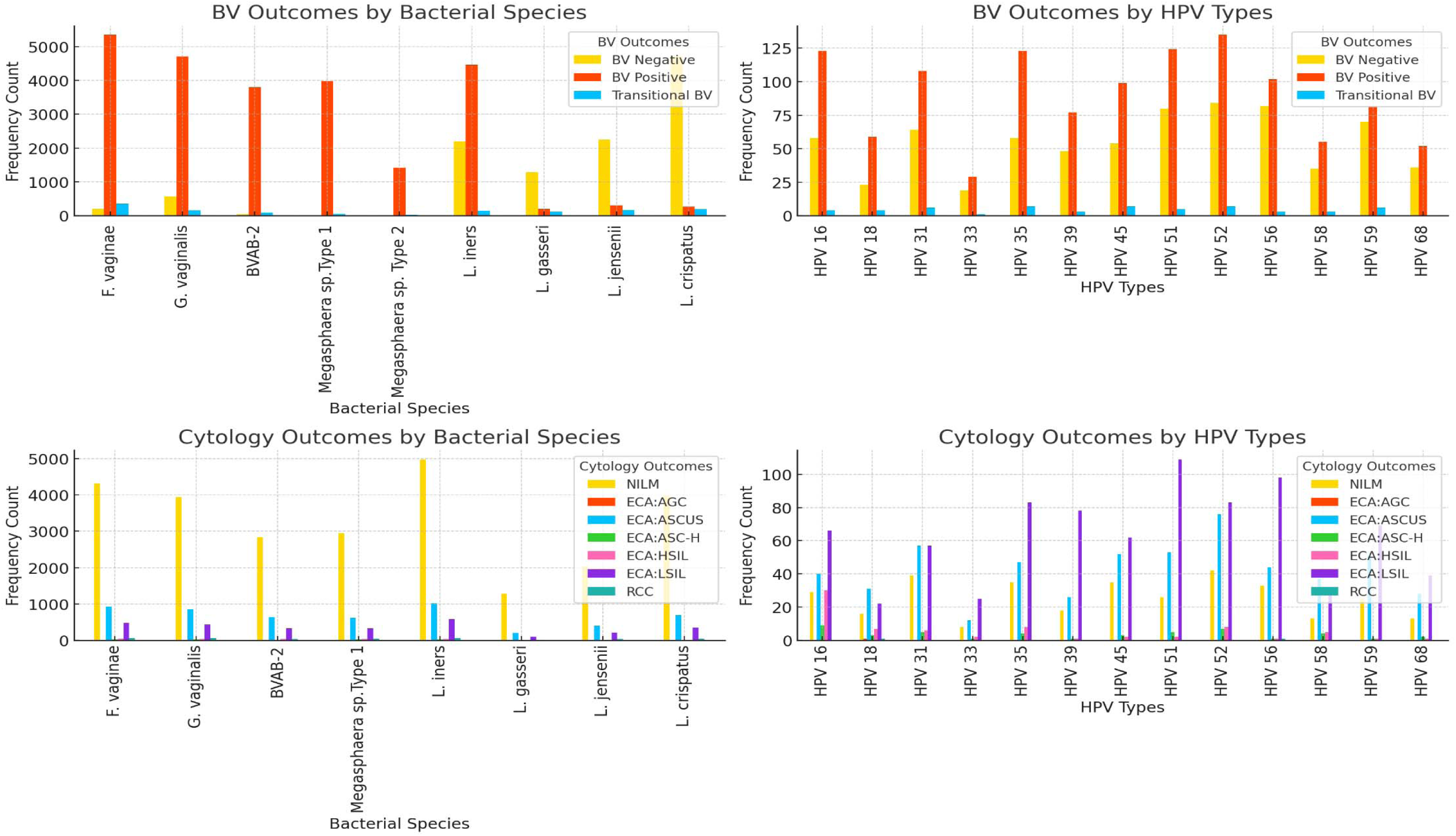
Association of bacterial species and HPV subtypes’ frequency with BV status and cervical cytology outcomes. F. vaginae, G. vaginalis, BVAB-2, Megasphaera sp., and L. iners s well as all HPV subtypes were associated with BV-positive samples while L. gasseri, L. jensenii, and L. crispatus were strongly associated with BV-negative samples. There were higher ssociation/presence of all HPV types with ECA-types (ASCUS, HSIL and SIL) than with NILM while NILM was highly associated with all the bacterial species than the other cervical cytology utcomes.

Correspondingly, there were higher mean concentrations of *F. vaginae* (89.60%)*, G. vaginalis* (93.23%)*, BVAB-2* (97.80%)*, Megasphaera sp. Type 1* (99.87%)*, Megasphaera sp. Type 2* (99.01%), and *L. iners* (58.85%) in BV-positive samples. Further, the mean concentrations of *L. gasseri* (56.33%)*, L. jensenii* (83.37%)*, L. crispatus* (80.57%) were high in BV-negative specimens and very minimal (1.75% – 0.3%) in BV-positive ones (Fig. S10-S15; Datasets 2.1-2.6). Contrary to the HPV subtypes’ counts per BV status however, the mean concentrations of HPV subtypes were not generally higher in BV-positive samples except HPV-33 (99.96%), HPV-35 (92.73%), HPV-39 (63.12%), and HPV-51 (59.89%); the remaining subtypes had percentage mean concentrations of 0.17% to 48.48% in BV-positive samples. None of the mean concentrations of HPV subtypes per BV status was significant (Fig. S10-S15; Datasets 2.1-2.6).

#### Cervical cytology (Pap smear) outcome associations

Whereas the bacterial species were all commonly detected in NILM, ASCUS, and LSIL-positive samples in descending order, their mean concentrations varied by species with *F. vaginae, G. vaginalis, BVAB-2,* and *Megasphaera sp.* Types 1-2 being relatively higher in HSIL, ASCUS, LSIL, and AGC than in NILM in many cases. Generally, the concentrations of the above-listed species were higher than those of the *Lactobacillus* species in the HSIL, ASCUS, ASC-H, LSIL, and AGC-positive samples. However, the *Lactobacillus* species had higher mean concentrations in the NILM-positive specimens. The HPV-positive specimens, however, were mostly HSIL, ASCUS, ASC-H, LSIL, and AGC-positive than NILM-positive and except for HPV-18, HPV-33, HPV-35, had higher concentrations in ECA-type pathologies (Fig. S10-S15; Datasets 2.1-2.6). A heatmap showing the chi-square p-values of these associations is shown in Fig. S16.

#### Correlation and pairwise analyses

The correlation matrix plot (Fig. S17; Dataset 3.1) confirmed the above findings: the bacterial pathogens were strongly and positively correlated with the presence of BV while the three *Lactobacillus* species (*gasseri, jensenii,* and *crispatus*) were strongly negatively correlated. There was a strong negative correlation between NILM and ASCUS, SIL, HSIL, AGC and ASC-H, and a positive correlation between the HPV subtypes and LSIL. HSIL was strongly positively correlated with HPV-16. NILM was positively correlated with age while all BV, the ECA-types, HPV subtypes, and bacterial species were weakly negatively correlated with age. A Chi-square and ANOVA pairwise analyses of the various factors showed significant association between age, State, BV status, ASCUS, and LSIL and most of the other factors tested, confirming several observations from the correlation matrix (Fig. S18).

### 3.3 Lactobacillus effect on HPV and pathogenic bacteria

We tested the effect of each *Lactobacillus* species on the five bacterial pathogens and 13 HPV subtypes. As shown in Fig. 3, there was a significant reduction in bacterial pathogens and HPV types in the presence of *L. gasseri, L. jensenii,* and *L. crispatus* than in the presence of *L. iners.* A Spearman correlation matrix showed stronger negative correlation between *L. crispatus* and *L. gasseri* and the bacterial pathogens in terms of their concentrations (Fig. S19). Notably, A chi-square analysis showed significant association of *L. iners, L. gasseri, and L. crispatus* with most of the HPV subtype and bacteria pathogens influencing BV and cervical cytology while *L. jensenii* had mostly insignificant associations with HPV. The odds ratio forest plot further showed the protective effect of the different *Lactobacillus* species (odd ratio < 1) against the occurrence of certain bacteria species, HPV types, and cervical cytology outcomes (Fig. 4; Datasets 3.3).

**Figure 3.**
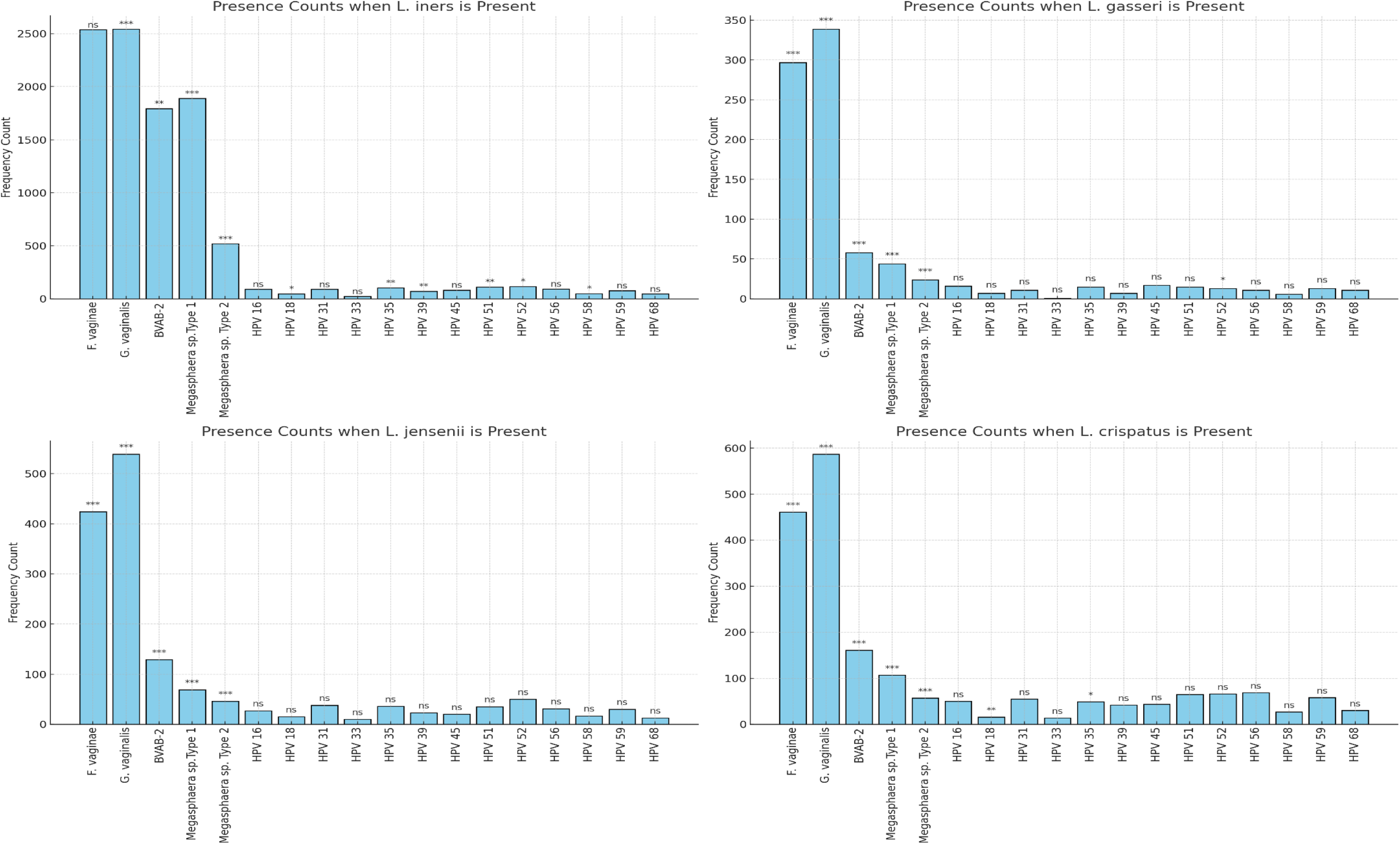
Frequency count of various bacterial pathogens and HPV types in samples that are positive for the various Lactobacillus species. Except for L. iners, the other Lactobacillus species were associated with lower counts of pathogenic bacteria and HPV in the samples. The associations were mostly significant for the bacterial species and some HPV subtypes.

**Figure 4.**
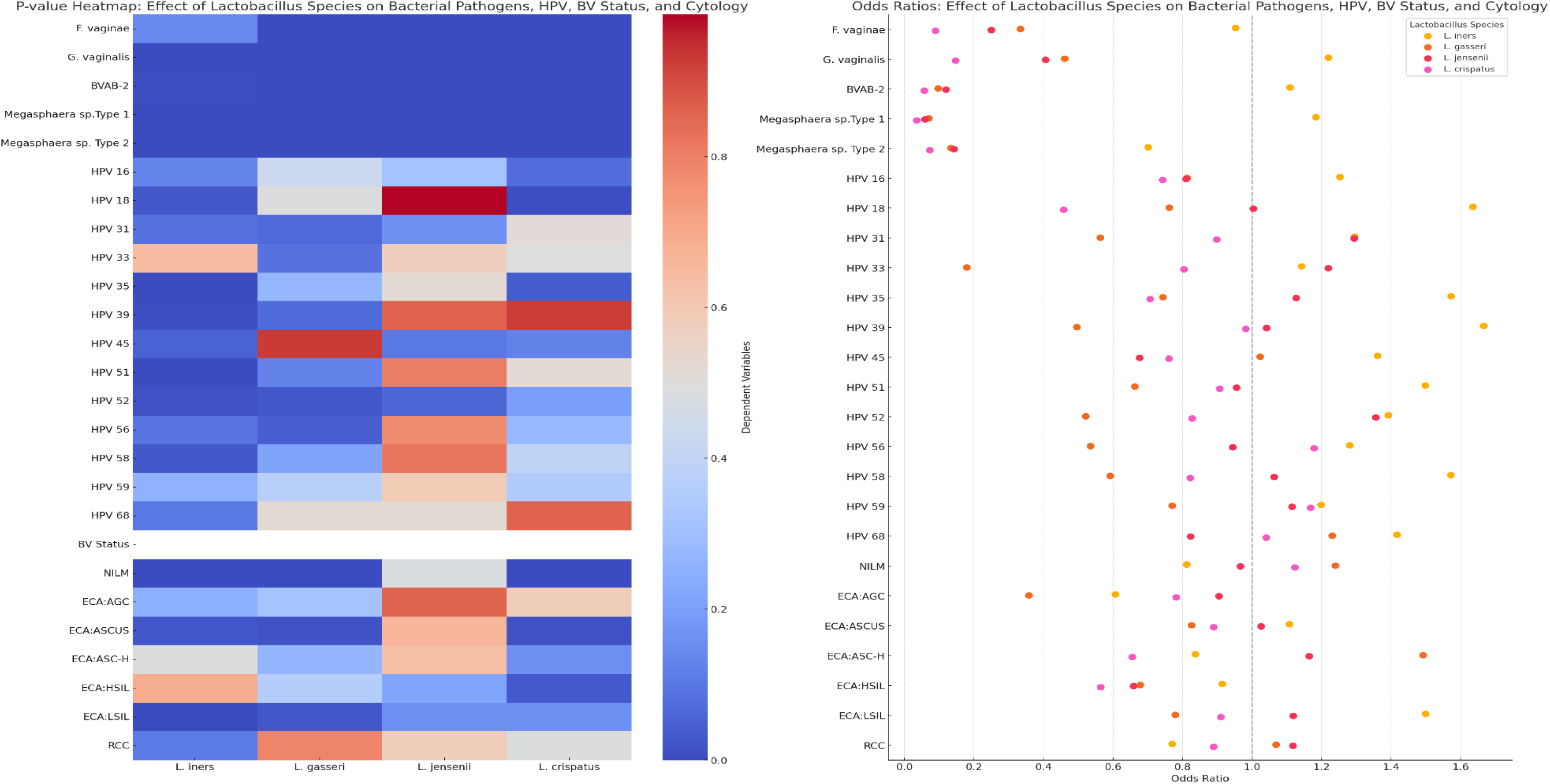
Chi-square statistical significance heatmap (left) and forest plot of odds ratios (right) showing the protective effect of the various Lactobacillus species against the various HPV subtypes, pathogenic bacteria, and cervical cytology outcomes. **Odds ratios greater than 1** indicate an increased likelihood of the outcome in the presence of the Lactobacillus species. **Odds ratios less than 1** indicate a decreased likelihood (protective effect) of the outcome in the presence of the Lactobacillus species. The vertical line at **1** indicates no effect.

Notably, for several HPV subtypes (HPV 16) and bacterial pathogens (*G. vaginalis*), *L. crispatus* had odds ratios < 1, indicating a protective effect. *L. iners*, however, had odds ratios > 1 for several pathogens, particularly BVAB-2 and Megasphaera sp. Type 1, suggesting that its presence may increase the likelihood of detecting these harmful bacteria. *L. gasseri* and *L. jensenii* showed mixed effects, with *L. gasseri* tending toward a protective role (odds ratios < 1) against HPV 45 and some bacterial pathogens, while *L. jensenii* showed potential protection against HPV 35 (Fig. 4).

### 3.4 Lactobacillus-pathogens interaction effects

An interaction effect analysis of all the bacterial species and HPV subtypes showed significant negative effect on vaginal and cervical health when certain HPV subtypes and/or bacterial pathogens co-occurred together. Specifically, HPV-16 and HPV-31, HPV-45 and HPV-51, HPV-52 and HPV-58, HPV-16, HPV-16 and *G. vaginalis,* and HPV-68 and *F. vaginae,* among others, were associated with ECAs. However, most of the non-*L. iners Lactobacillus-*pathogen interactions with bacterial and HPV pathogens resulted in BV negative or NILM outcomes. A detailed breakdown is shown in Supplemental Datasets 3.4 – 3.9 and 5 (Fig. S20 – S25).

### 3.5 Principal Component and K-means clustering Analysis

Principal component analysis (PCA) was undertaken to determine the variance and spread of the various bacterial species towards the two components (outcomes), P1 and P2. The loadings of PC1 are dominated by species such as *F. vaginae, G. vaginalis*, and BVAB-2, suggesting that these bacterial species contribute heavily to the variance along PC1. *L. iners* had a significant loading on PC2, meaning it influences variability in a different direction compared to other species. HPV types, such as HPV-59 and HPV-45, also had notable loadings on PC2, indicating their contribution to variance along that component (Fig. S26; Dataset 4.1).

An average bacterial and HPV subtype concentrations per K-means clusters 0, 1, and 2 (Fig. S27 – S28) show that cluster 1 is associated with ASCUS, HSIL, BV-positive or transitional BV cases owing to the higher concentrations of *F. vaginae, G. vaginalis,* BVAB-2, *Megasphaera sp.* Type 1, *L. iners,* HPV-16, HPV-35, HPV-45, HPV-59, and HPV-68; other subtypes such as HPV-18, HPV-31, and HPV-33 also had considerable concentrations. However, cluster 2 had elevated concentrations of *L. iners, L. crispatus, L. jensenii*, and lower HPV concentrations, except HPV 31 and HPV 56, suggesting that this cluster is likely associated with a healthy vaginal microbiome, normal cytology (NILM), or HPV-negative cases (Fig. S27 – S28; Dataset 4.2).

### 3.6 Feature importance on cervical cytology and BV outcomes

Based on their precision, recall, and F1 metrics, XGBoost and Random Forest were the strongest performers across both outcomes (BV and Cervical Cytology). The four models identified *L. crispatus, L. jensenii, L. gasseri, F. vaginae,* and *G. vaginalis,* as most important factors in predicting the occurrence of BV. There was also agreement among the models on the importance of age, hrHPV outcome, *L. iners, L. crispatus, L. jensenii,* HPV-51, *G. vaginalis, F. vaginae,* and *Megasphaera sp. Type 1* on the occurrence/prediction of cervical cytology outcomes (Fig. 5). However, the accuracy of all the models were higher in BV than in cervical cytology predictions (Datasets 4.3 – 4.11).

**Figure 5.**
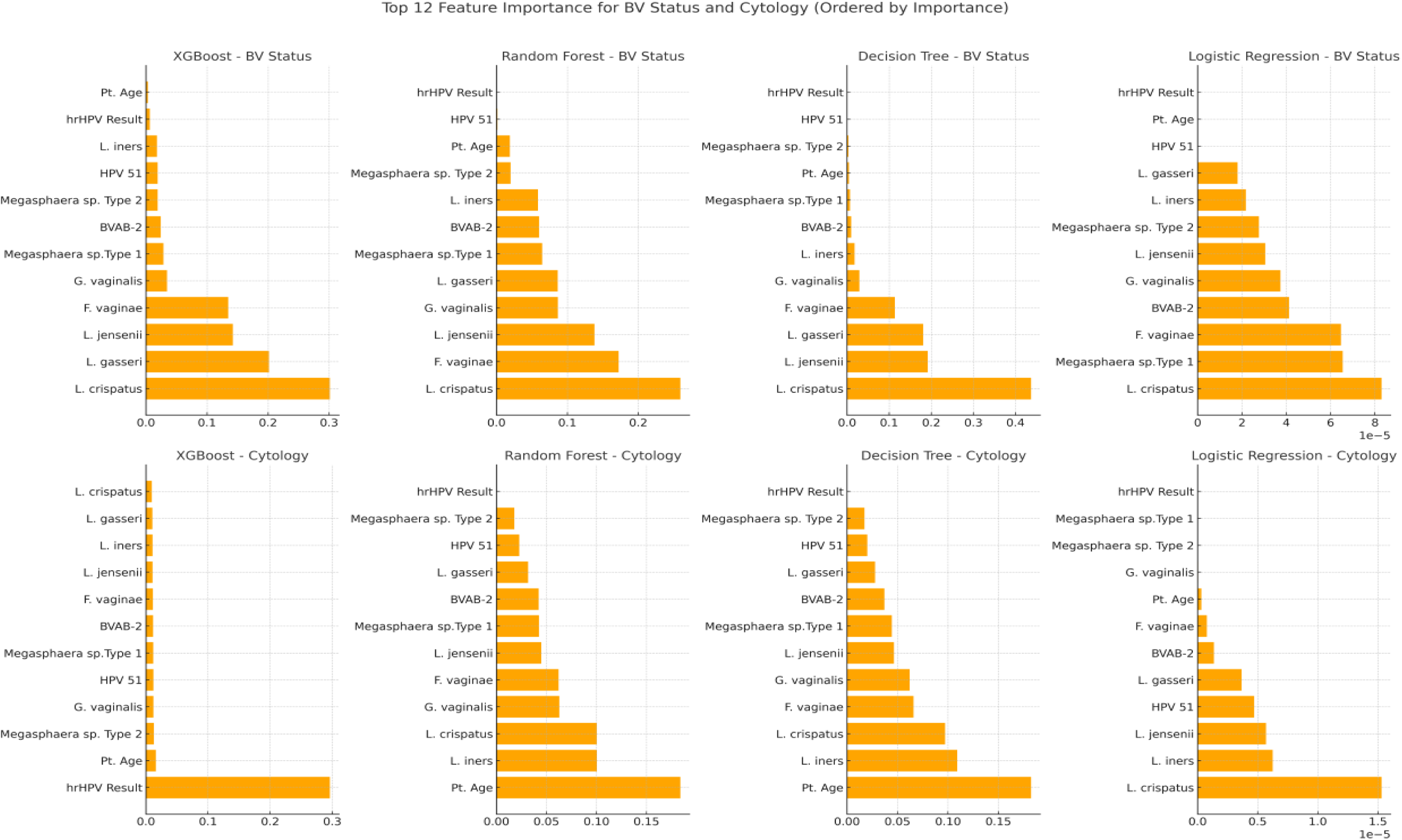
Feature importance of various determinants/factors predicting the outcome of BV and cervical cytology as determined using XGBoost, Random F rest, Decision Tree, and Logistic Regression Machine-Learning models. Regardless of their differences in accuracy and performance indices, the four models agree on the importance of common bacterial species, HPV types, and age that influence the outcomes of BV and cervical cytology.

## 4.0 Discussion

The clinical effect of *L. crispatus, L. gasseri,* and *L. jensenii* on the incidence of hrHPV subtypes, BV, and ECAs were determined using 15601 clinical data from 15601 anonymized patients that reported to healthcare providers in 32 states and the D. C, USA. The series of molecular, pathological and statistical analyses undertaken on this large cohort of samples all point to the protective effect of *L. crispatus* and to some extent, *L. gasseri* and *L. jensenii.* Specifically, *L. crispatus* was consistently the most protective Lactobacillus species across all visualizations. It reduces the likelihood of bacterial pathogens such as *G. vaginalis* and *F. vaginae,* as well as hrHPV subtypes such as HPV 16 and HPV 18. The odds ratios (OR < 1), significant p-values, and lower presence counts all support this protective effect.

*L. iners* is not protective. In fact, it is associated with higher frequencies of bacterial pathogens such as *F. vaginae, G. vaginalis,* BVAB-2 and *Megasphaera sp.* Type 1, which are often associated with bacterial vaginosis and dysbiosis. The odds ratios > 1 and higher presence counts suggest that *L. iners* does not contribute to a healthy microbiome*. L. gasseri* and *L. jensenii* showed protective effects, particularly against some HPV subtypes such as HPV 45 and HPV 35, but their impact was less pronounced compared *to L. crispatus*. Indeed, these findings are not new as other studies have already shown these protective effects.^14–16^ However, this study uses a larger number of cervicovaginal samples from different racial backgrounds and States in the United States to establish these earlier findings.

The higher relative abundance of *L. iners* in the vaginal microbiome is already established, as well as the association of *F. vaginae, G. vaginalis,* BVAB-2 and *Megasphaera sp.* Type 1 and 2 with BV.^14,17^ This study expands on these by showing that these pathogens also have higher concentrations and prevalence in ECAs and could be risk factors for getting cervical cancer (as shown by the four machine learning models) as they indicate vaginal dysbiosis.^17,18^ Furthermore, while the effect of hrHPV on ECAs (also found in this study) is known, this study further shows a higher prevalence of HPV-subtypes in BV-positive specimens, with most hrHPV types having higher concentrations in BV-positive specimens than BV-negative ones. Indeed, the higher prevalence and concentrations of *Lactobacillus sp.* in NILM-positive samples evince their protective effect against ECAs.^15,19–21^

Interactive (co-existence) effects of multiple hrHPV (n = 1719 patients) and/or bacterial species led to significant increases in BV and ECA pathologies as shown in Dataset 5.2 (Fig. S20-S25). This shows that having multiple HPV infections and HPV-bacterial pathogens predisposes the host to higher risk of BV and ECA pathologies.^20^ The protective effects of *Lactobacilli* might explain why many studies are proposing the use of probiotics and prebiotics to fight BV and ECA instead of using chemotherapy alone.^18,19,21,22^ In clinical trials, *L. crispatus* has been found beneficial than placebos in reducing BV and vulvovaginal candidiasis.^15^

Demographic factors such as age and State were also significantly associated with the prevalence BV and hrHPV subtypes: AGC, HSIL, and RCC were found in populations older than the average.^14,23–25^ The higher presence of BV, hrHPV, and cervical cytology outcomes in some of the States should be investigated further to ascertain why those States are reporting higher proportions of these pathologies and infections. Most of the BV-and hrHPV-positive cases falling within ages 20-50 agree with other studies as this age group is the most sexually active.^26–29^ The presence of BV-causing bacteria, hrHPV (HPV-31 and 39) and associated ECAs (ASCUS and AGC) in males, albeit smaller in number (n = 13), is concerning as it suggests these pathogens can be sexually transmitted to female partners.^30^ hrHPV in males has been reported to be increasing globally, with a prevalence of 21%, which is more than the prevalence found in this study.^11^ An expanded study using more clinical samples from males will provide a better prevalence data on hrHPV among males in the USA.

Fortunately, the overall prevalence of ECAs was less than 15% albeit BV was 53.07%; this higher BV prevalence could be because the samples were from patients suspected of having BV. Moreover, HPV-16 and HPV-18, which are the most common hrHPV etiologies of ECAs,^11,31,32^ were not the most prevalent hrHPV types: HPV-52, 51, 35, and 56 were the most prevalent, followed by HPV-16. Lebeau et al. (2022) recently showed that HPV infections alter the cervicovaginal microbiome through down-regulation of the host’s innate peptides used by *Lactobacilli* to biosynthesize amino acids.^33^ This finding suggests that not only BV predisposes women to hrHPV infections, but the reverse can also hold true. In either case, the higher association of HPV and BV pathogens with BV and ECAs show their inter-relatedness in BV and ECA pathogenesis.^9,18,32^ Notwithstanding, not all hrHPV-positive specimens were classified as ECA, showing that not all hrHPV infections affect the cervical cytology negatively; particularly when the immune system can resolve many HPV-infections efficiently.^9,10^

In summary, *L. crispatus,* followed by *L. gasseri* and *L. jensenii*, play the strongest role in maintaining a healthy cervicovaginal microbiome by reducing the presence of harmful bacterial pathogens and high-risk HPV subtypes, while *L. iners* may have an opposing or neutral effect. Coinfection with multiple hrHPV and/or hrHPV-BV pathogens increased the risk of BV and ECA outcomes and should be prioritized. Age and State had a significant effect on the prevalence of BV and cervical cytology outcomes and the public health departments of States with higher BV and ECA outcomes must take notice.

## Acknowledgements

The authors are grateful to the technicians at Medical Diagnostic Laboratories for their direct and indirect assistance. The material and financial resources provided by Medical Diagnostic Laboratories towards this project are warmly acknowledged and deeply appreciated. We are also grateful to Annette Daughtry for assisting with the review of the initial draft.

## Transparency declaration

The authors are scientists working at Medical Diagnostic Laboratories, LLC.

## Ethics statement

Ethics approval was not required for this study as only anonymized electronic records were used, posing no risk of identifying individual patients. According to the Common Rule (45 CFR 46) and HIPAA (45 CFR 164.514(b)(2)), research involving anonymized data does not constitute human subjects research and thus does not require IRB approval. Consent procedures: Written informed consent was not obtained as the study used anonymized electronic records and the study did not involve direct interaction with patients.

## Supplementary files

**Supplemental Table**. Anonymized dataset containing the raw data of the demographics, bacteria and hrHPV qPCR data, hrHPV, BV and cervical cytology diagnoses.

**Supplemental Datasets 1**. Datasets containing demographic, descriptive, and exploratory statistical analyses.

**Supplemental Datasets 2**. Datasets containing statistics, counts and mean concentrations of bacterial species and HPV subtypes per BV and cytology outcomes.

**Supplemental Datasets 3.** Datasets containing association, pairwise statistics, correlation, and interactive effects analyses of the demographics, bacteria, HPV, BV and cervical cytology outcomes.

**Supplemental Datasets 4.** Datasets containing principal component analysis (PCoA), k-mean clusters, and machine learning outputs.

**Supplemental Dataset 5.** Appendices of definitions, detailed results and analysis of interactions effects, PCoA, K-means clustering heatmaps, and machine learning metrics.

**Supplemental Figures S1 - S28.** Supplemental Figures S1 to S28 are shown in this file.

---

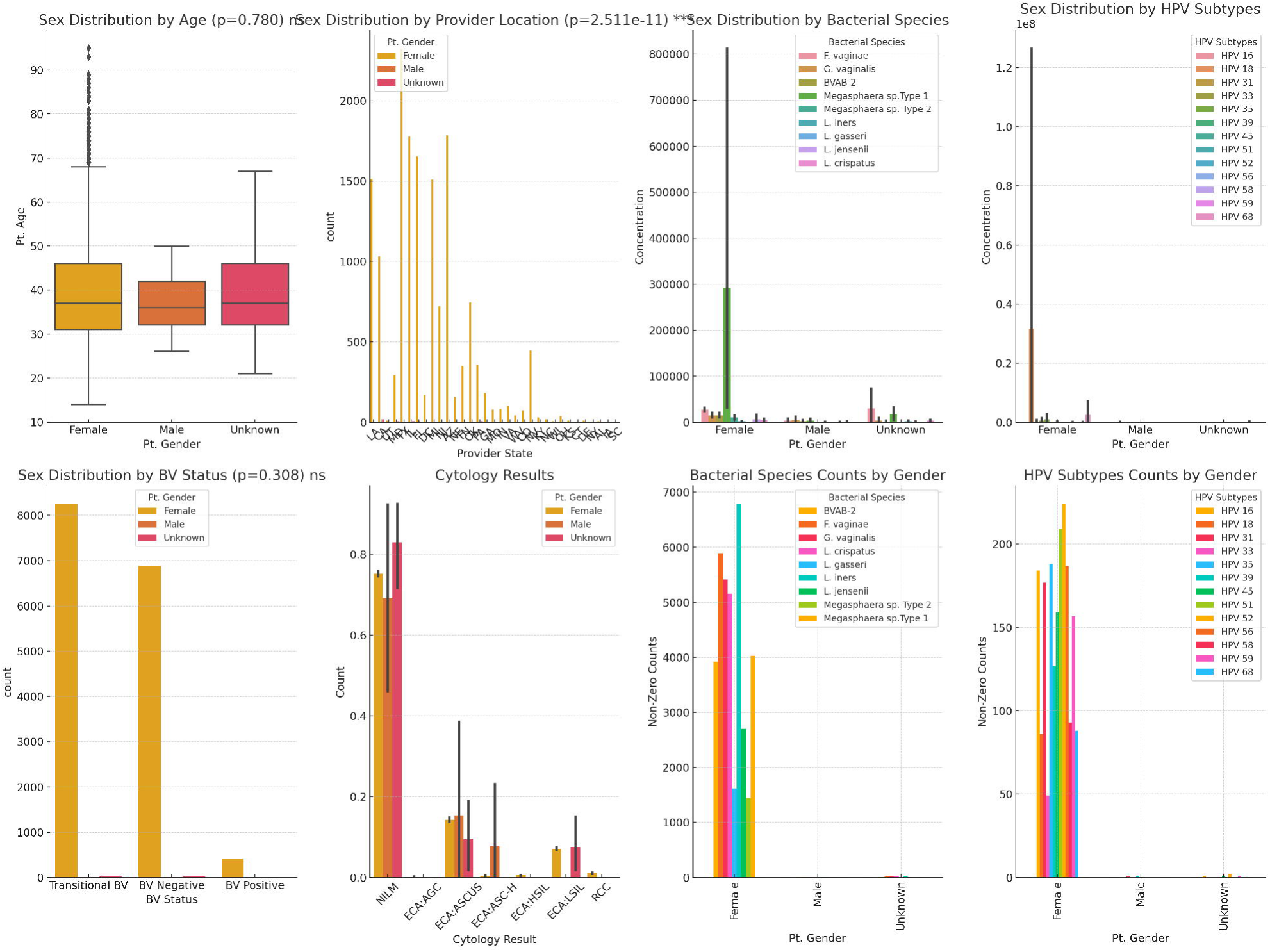

