## Supplemental Figures 1-28 for "Molecular and Pathological Characterization of Cervicovaginal Specimens Evince Significant Protection Against HPV Infections, Bacterial Vaginosis, and Epithelial Cell Abnormalities by *Lactobacillus* species"

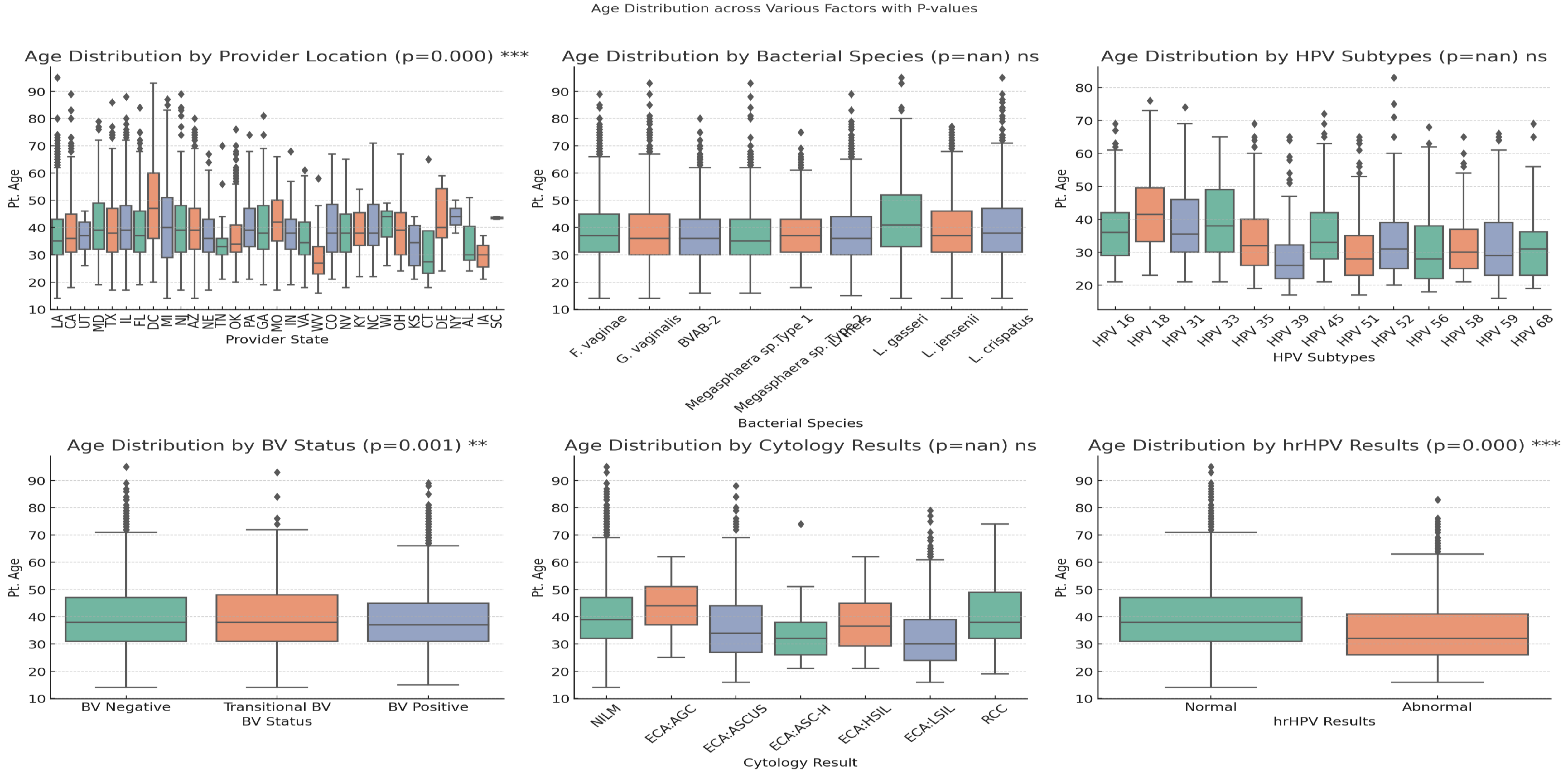


Fig. S1. Age distribution across States (provider location), bacterial species, HPV subtypes, BV status, cytology results, and hrHPV results. Only provider location, BV status, and hrHPV results had significant association with age. Notably, age per bacteria species and BV status had little variations except for L. gasseri that was mostly detected in patients aged between 35 and 50. HPV types 18 and 33 mostly occurred in older populations than the rest while samples from Delaware, Missouri, Colorado, and DC were from older patients than the rest. Further, AGC, HSIL, and RCC were mostly in patients older than the other outcomes. In all, hrHPV-positive samples were in a younger population than normal ones.

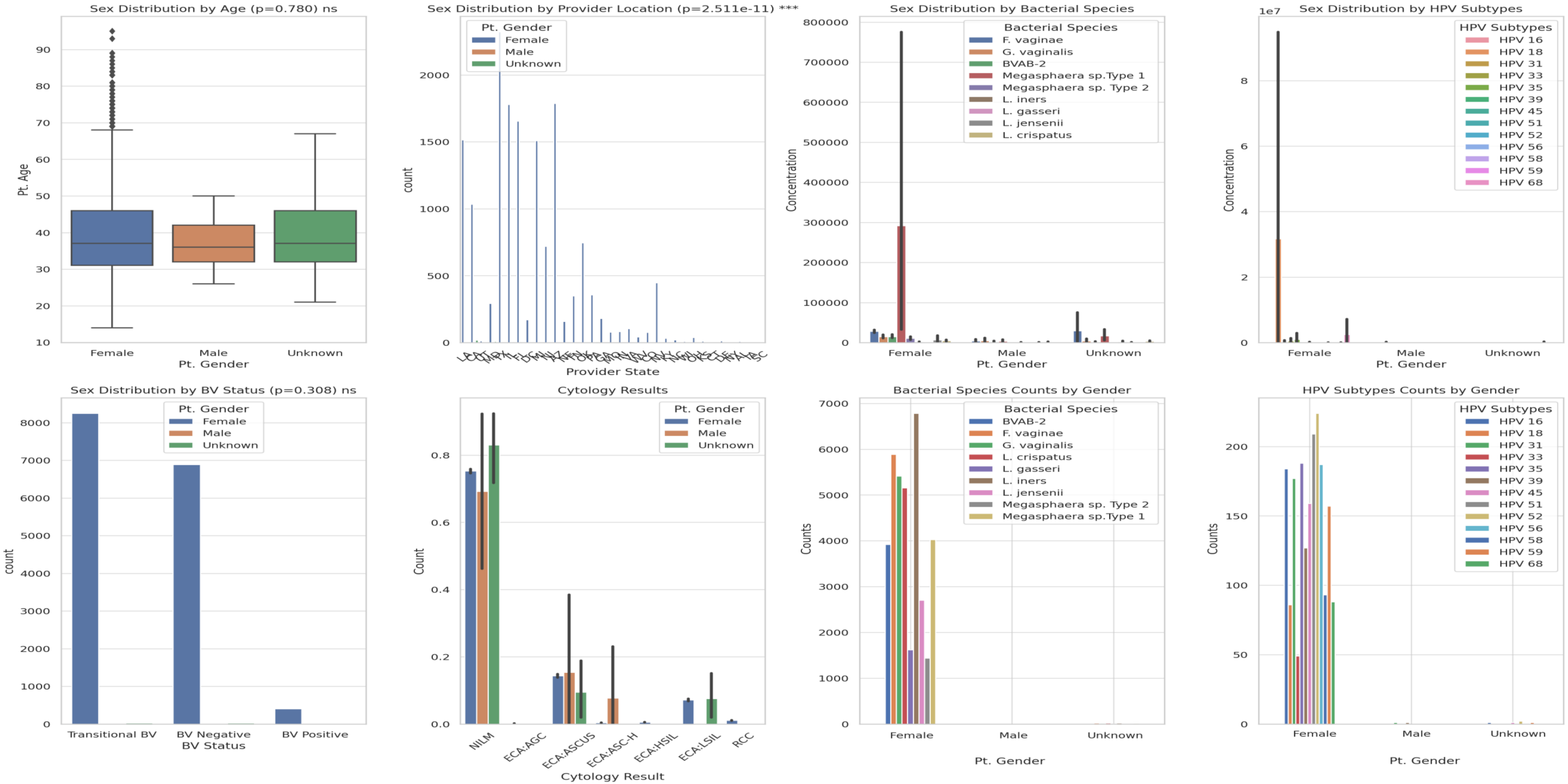


Fig. S2. Distribution of gender across age, provider location (States), bacteria species, HPV types, BV status, and cytology results. Only provider location was significantly associated with gender while the rest were not. Most bacterial species and a few HPV types were found in specimens from males and unknown gender. ECA such as ASCUS and AGC were found in males and unknown while LSIL was found unknown genders.

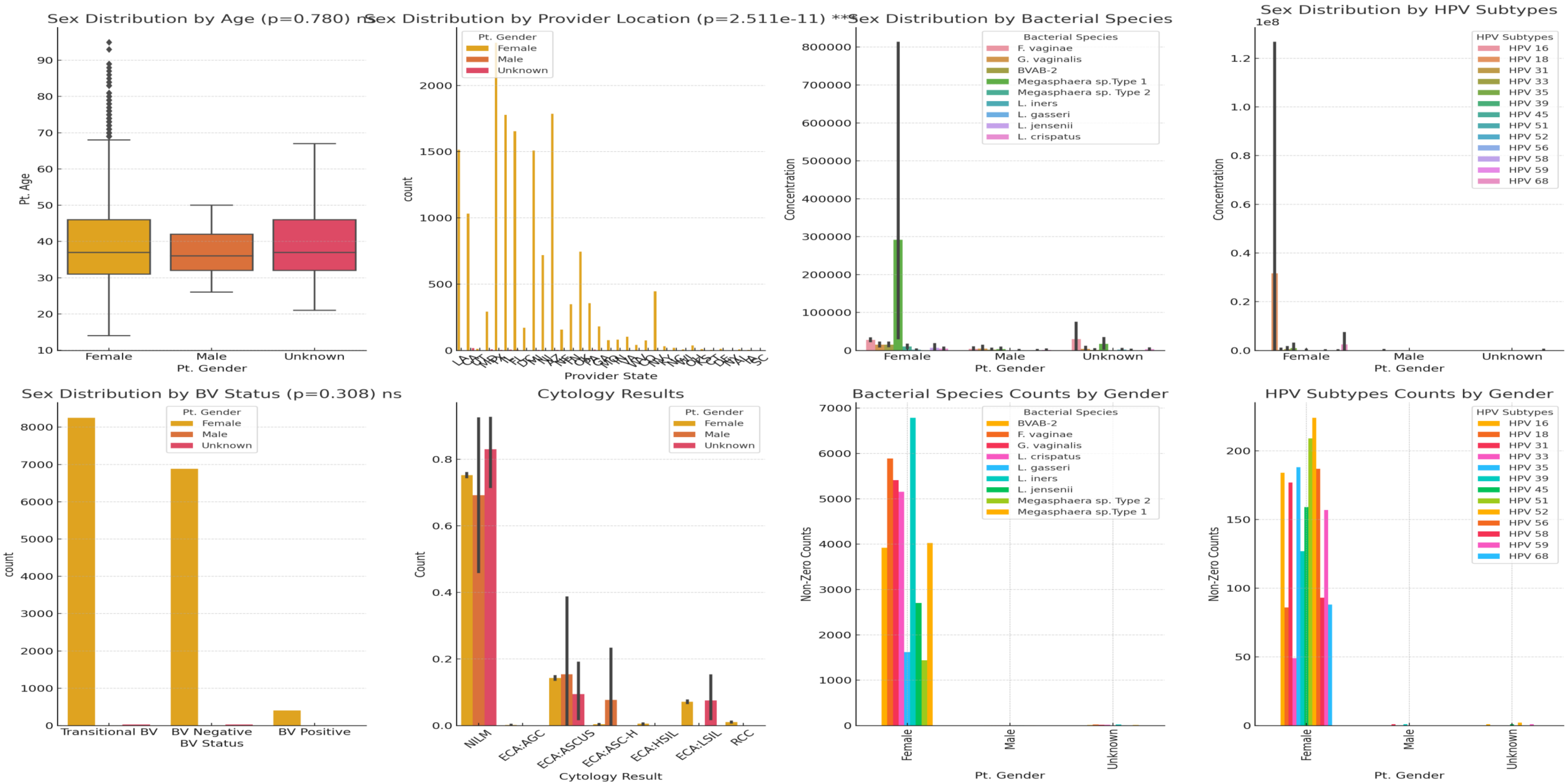


Fig. S2b. Distribution of gender across age, provider location (States), bacteria species, HPV types, BV status, and cytology results. Only provider location was significantly associated with gender while the rest were not. Most bacterial species and a few HPV types were found in specimens from males and unknown gender. ECA such as ASCUS and AGC were found in males and unknown while LSIL was found unknown genders.

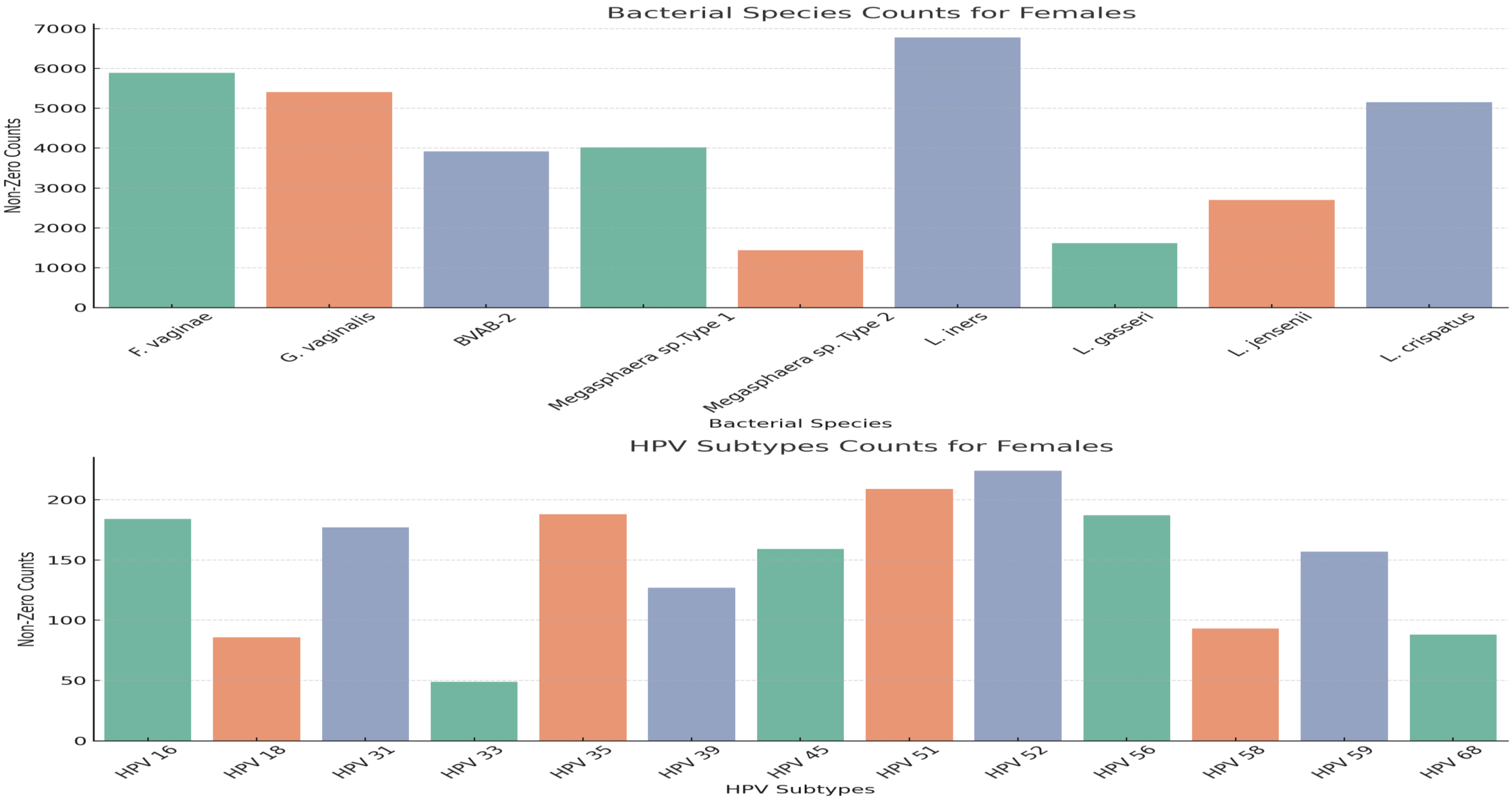


Fig. S3. Expanded view of counts of bacterial species and HPV types found in females

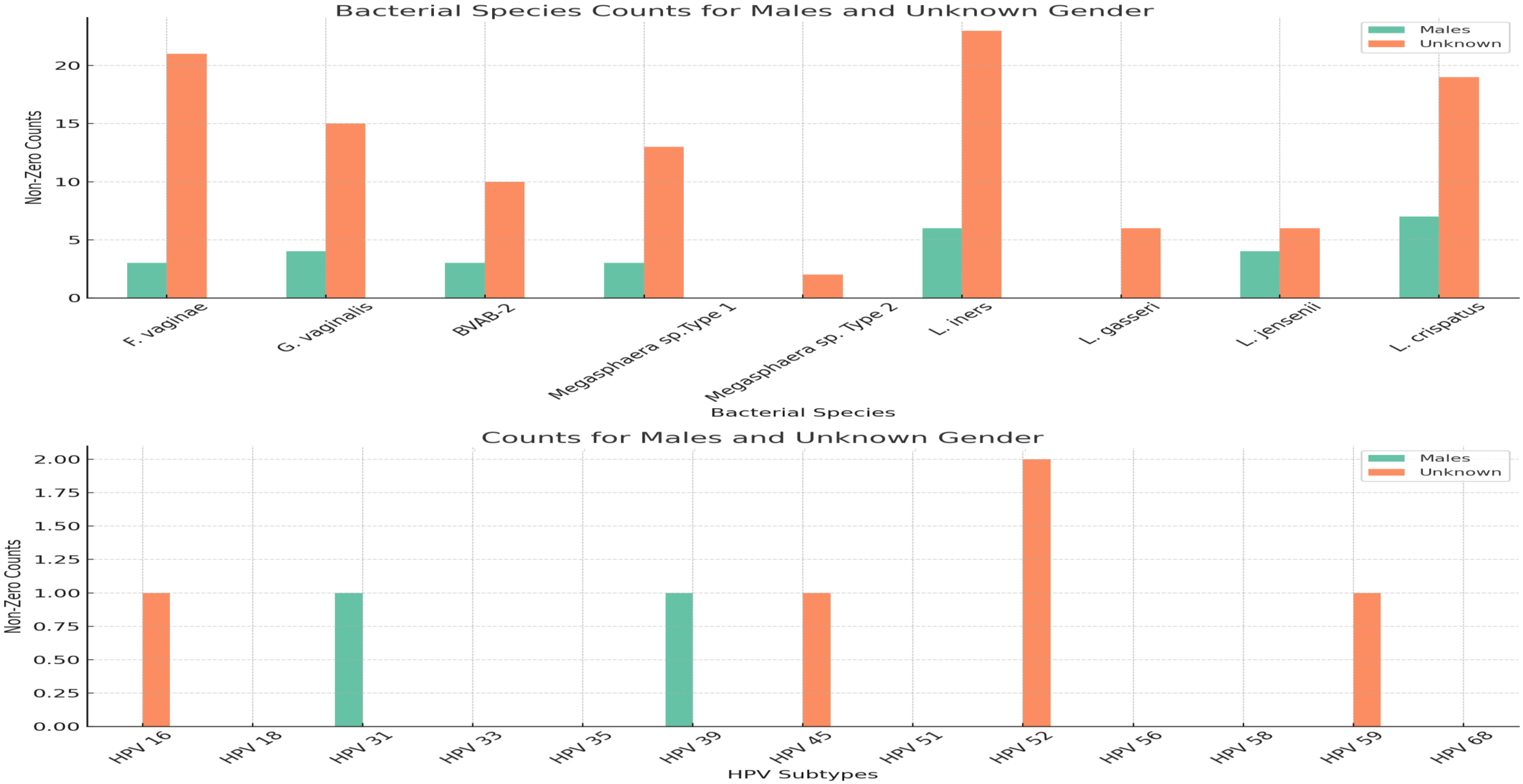


Fig. S4. Expanded view of counts of bacterial species and HPV types found in males and unknown gender

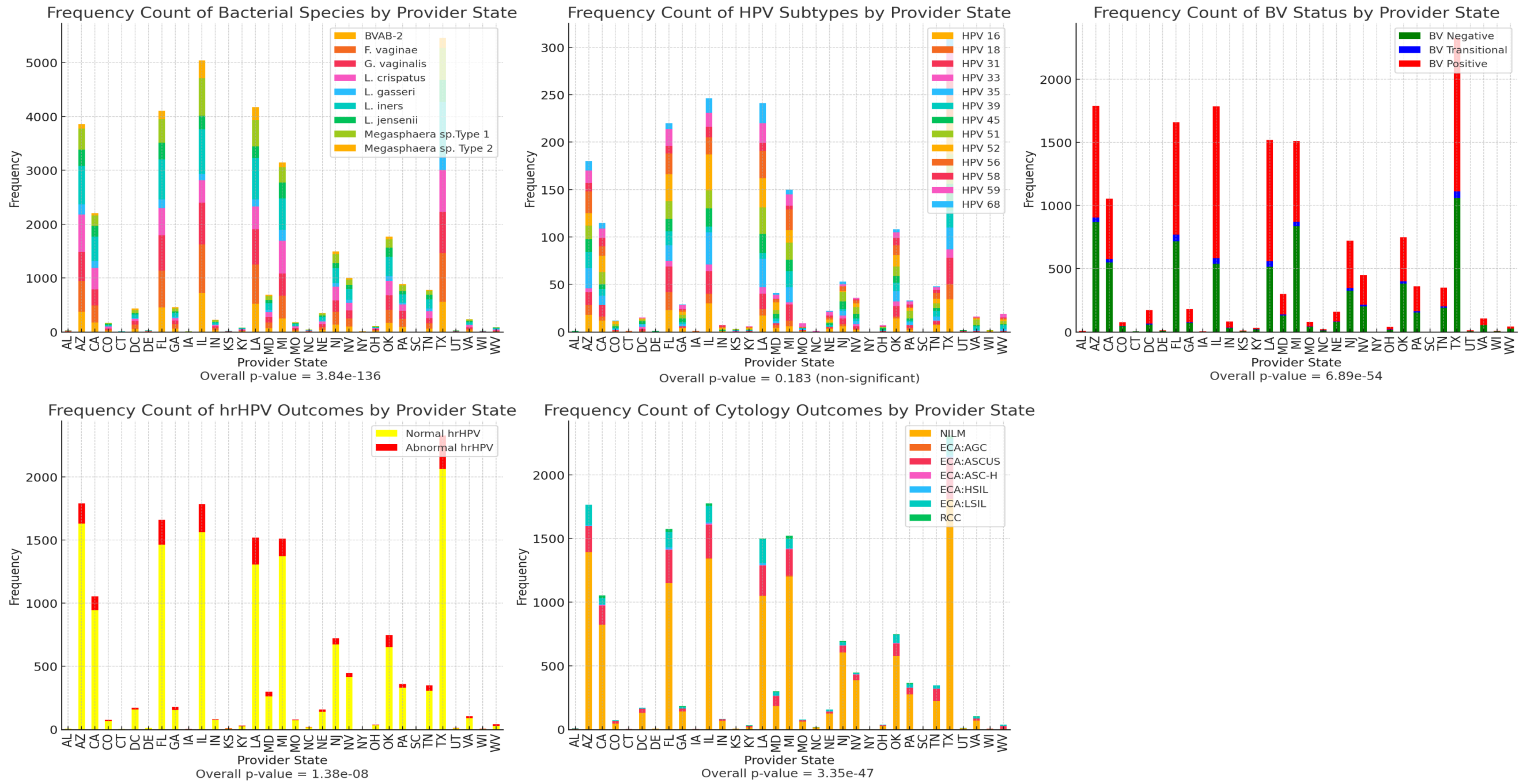


Fig. S5. Distribution of bacterial species, HPV subtypes, BV status, hrHPV outcome, and cervical cytology (pap smear) diagnosis per State (healthcare provider location). *L. iners, F. vaginae, G. vaginalis,* and *L. crispatus* were dominant in samples from almost all States. Samples from most states were BV positive, NILM, and hrHPV negative. States are abbreviated.

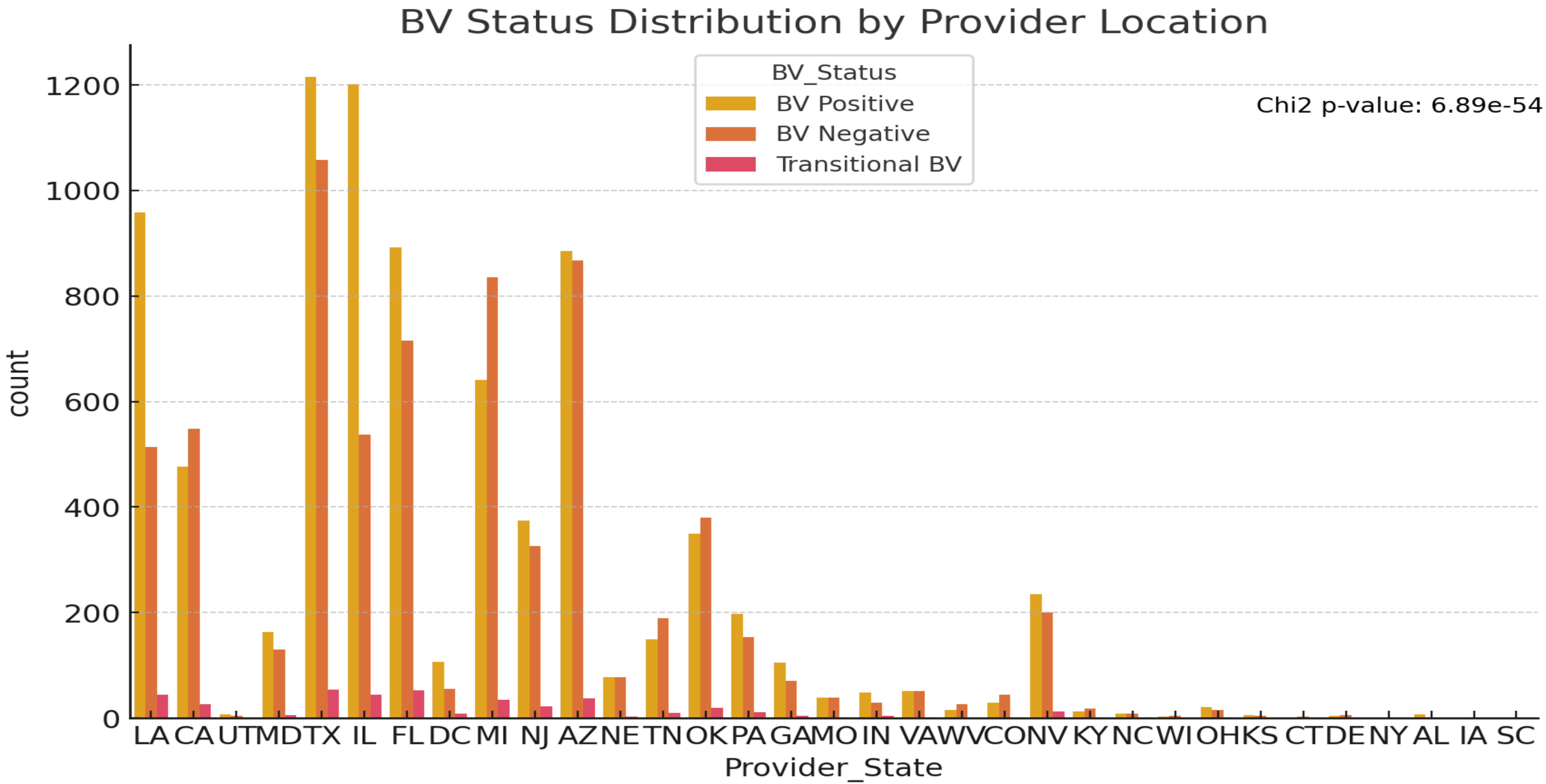


Fig. S6. BV Status distribution per State (provider location). The counts of BV-positive, BV-negative, and Transitional BV per healthcare provider State is shown above in different column colours. States are abbreviated.

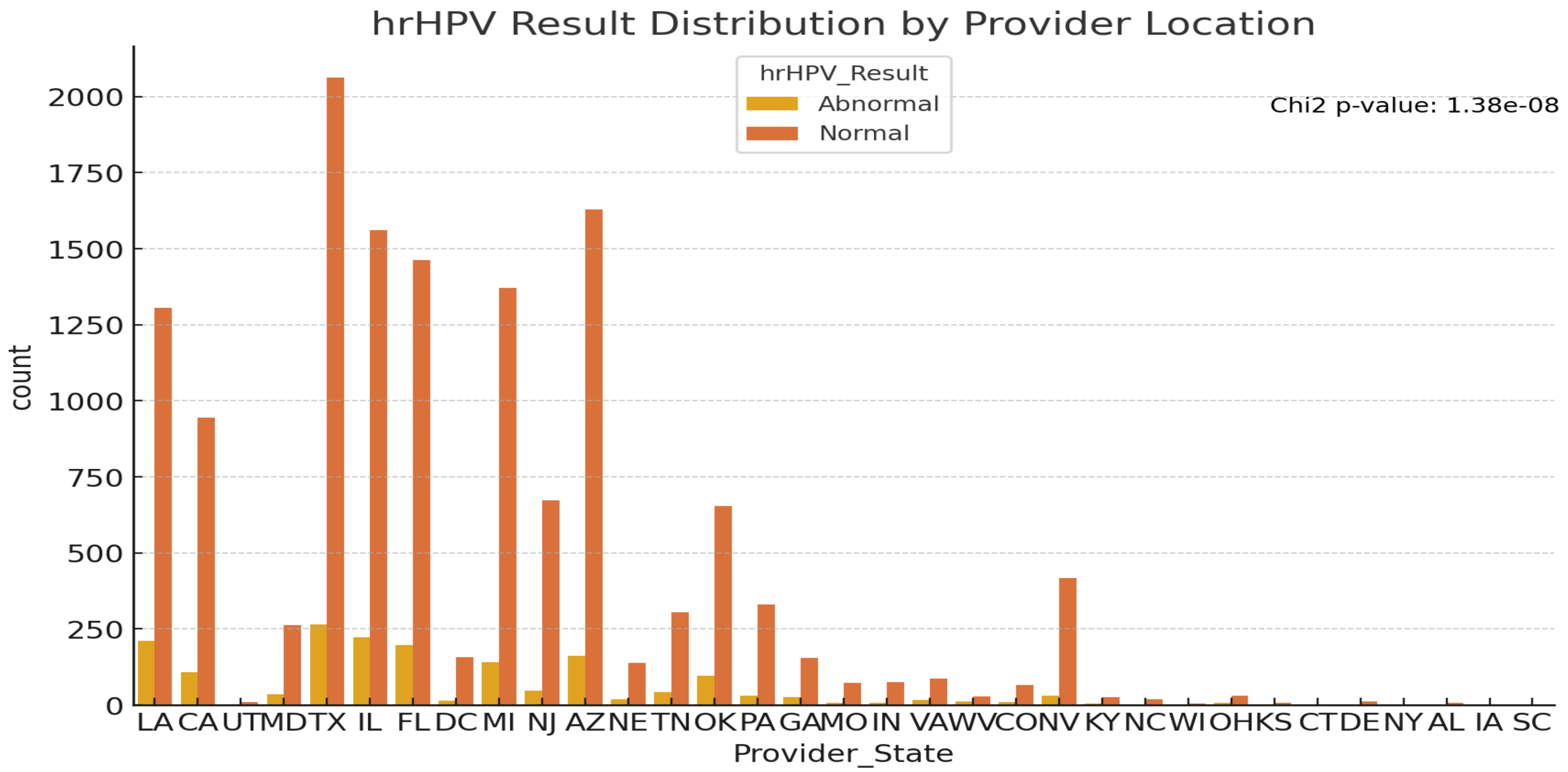


Fig. S7. hrHPV results distribution per provider location (State). Abnormal hrHPV (i.e., hrHPV-positive) specimens were substantially lower than normal (hrHPV-negative) specimens in most states, reflecting the generally lower prevalence of hrHPV in the USA.

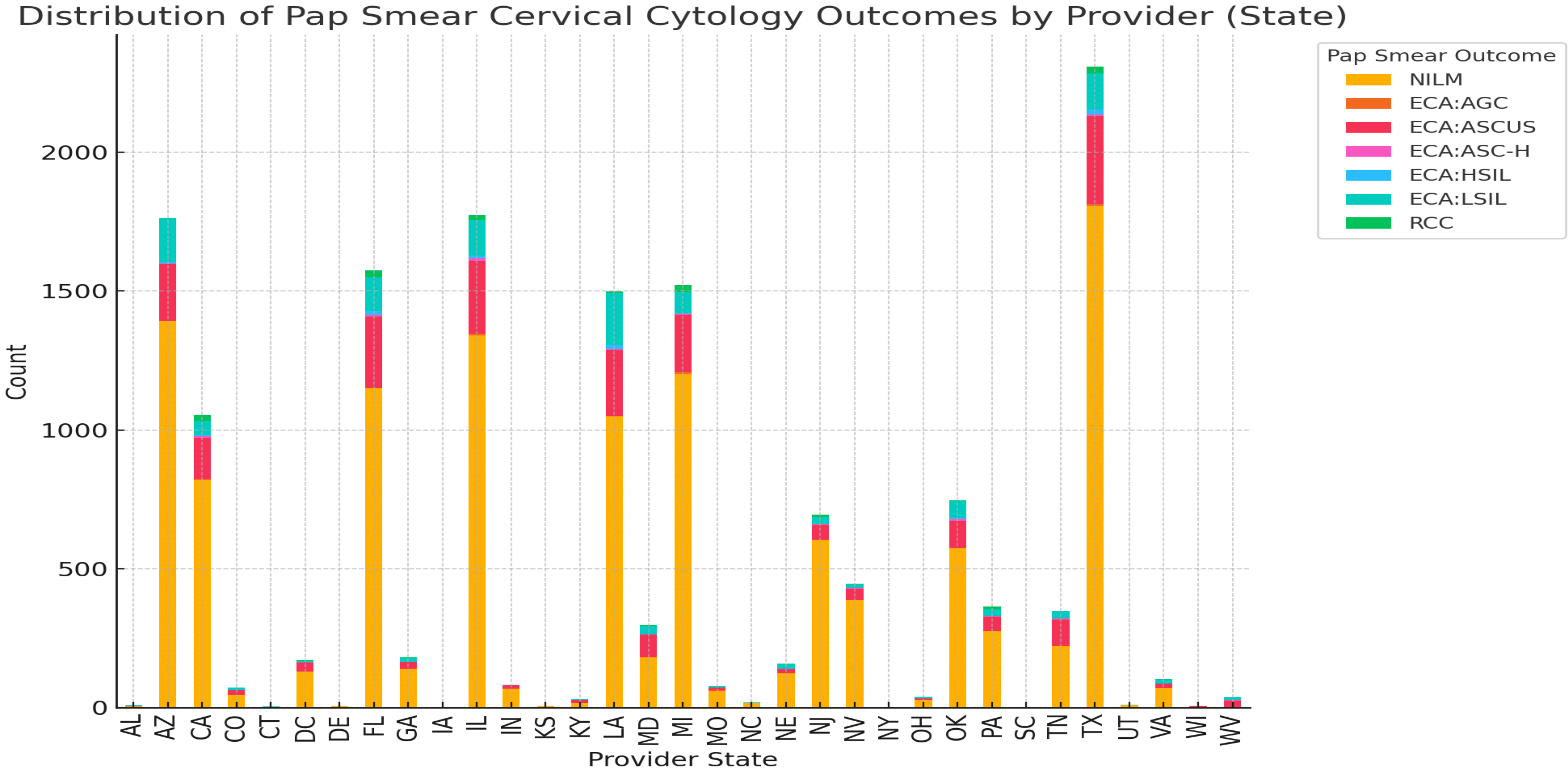


Fig. S8. Cervical cytology results from Pap smear analysis. The different cervical cytology outcomes are represented by different colours per State. Similar to the hrHPV results, most States had a higher prevalence of NILM outcomes than ECA.

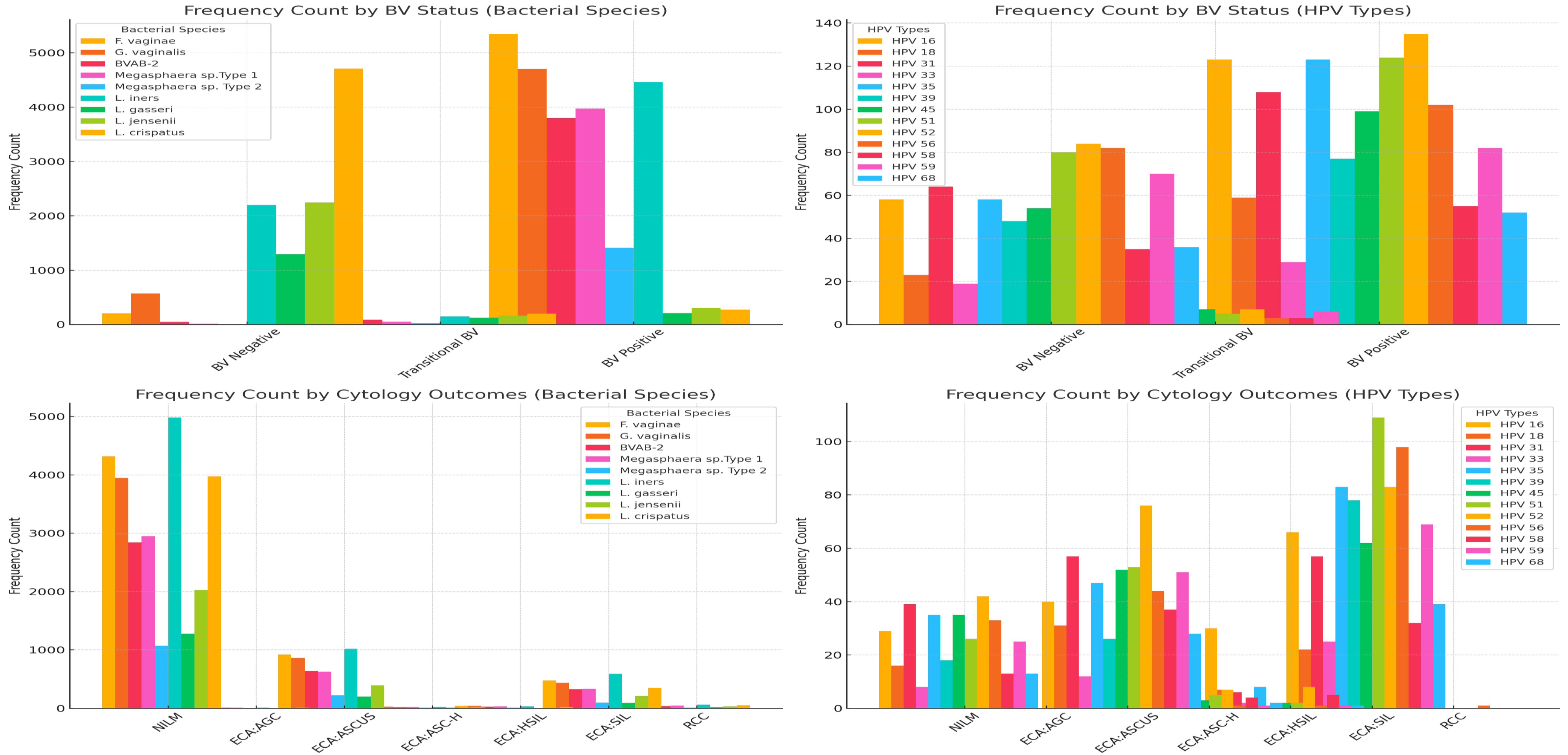


**Fig. S9.** ***Association of bacterial species and HPV subtypes' frequency with BV status and cervical cytology outcomes.*** *F. vaginae, G. vaginalis, BVAB-2, Megasphaera sp., and L. iners as well as all HPV subtypes* were associated with BV-positive samples while *L. gasseri, L. jensenii, and L. crispatus were strongly associated with BV-negative samples. There were higher association/presence of all HPV types with ECA-types (ASCUS, HSIL and SIL) than with NILM while NILM was highly associated with all the bacterial species than the other cervical cytology outcomes.*

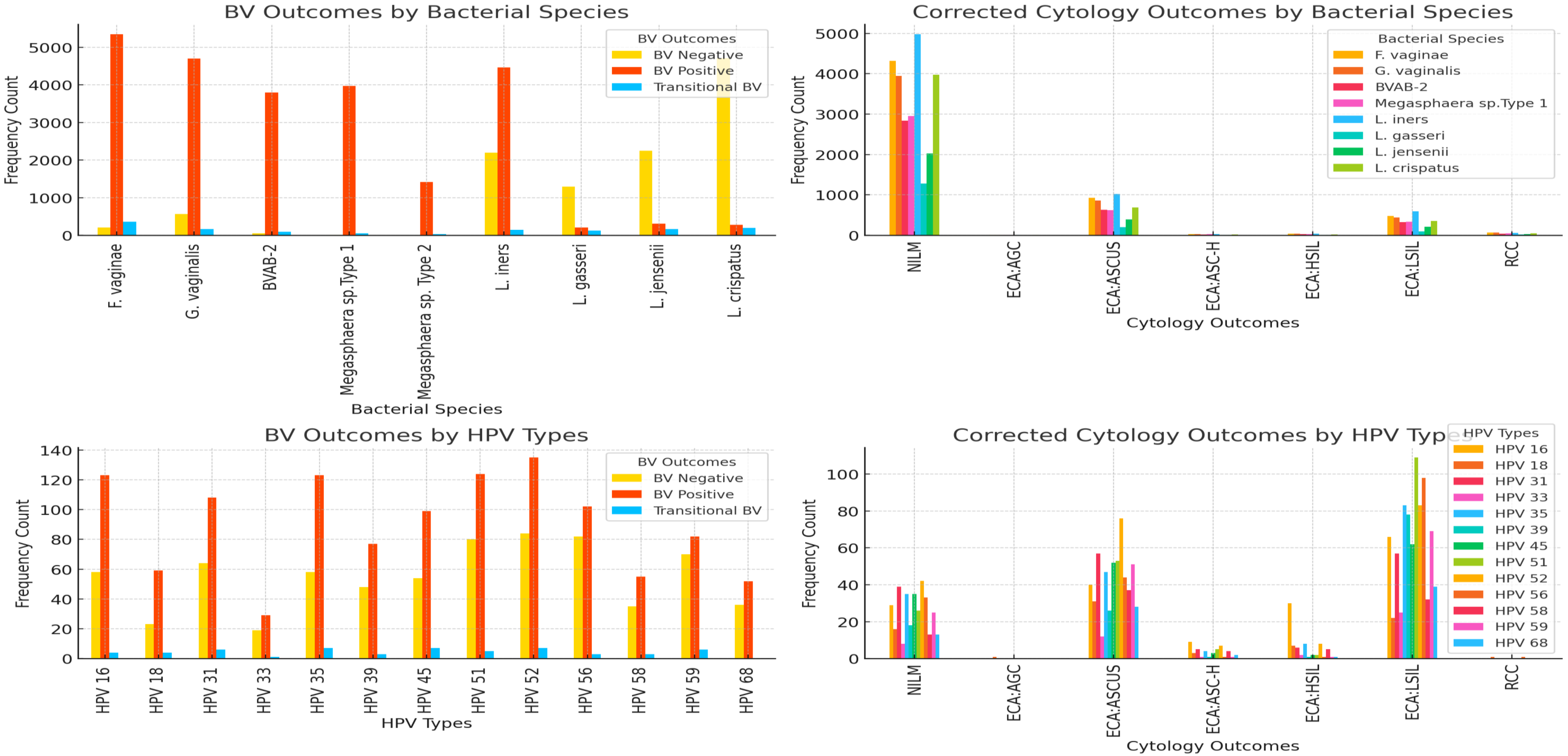


**Fig. S9b.** ***Association of bacterial species and HPV subtypes' frequency with BV status and cervical cytology outcomes.*** *F. vaginae, G. vaginalis, BVAB-2, Megasphaera sp., and L. iners as well as all HPV subtypes* were associated with BV-positive samples while *L. gasseri, L. jensenii, and L. crispatus were strongly associated with BV-negative samples. There were higher association/presence of all HPV types with ECA-types (ASCUS, HSIL and SIL) than with NILM while NILM was highly associated with all the bacterial species than the other cervical cytology outcomes.*

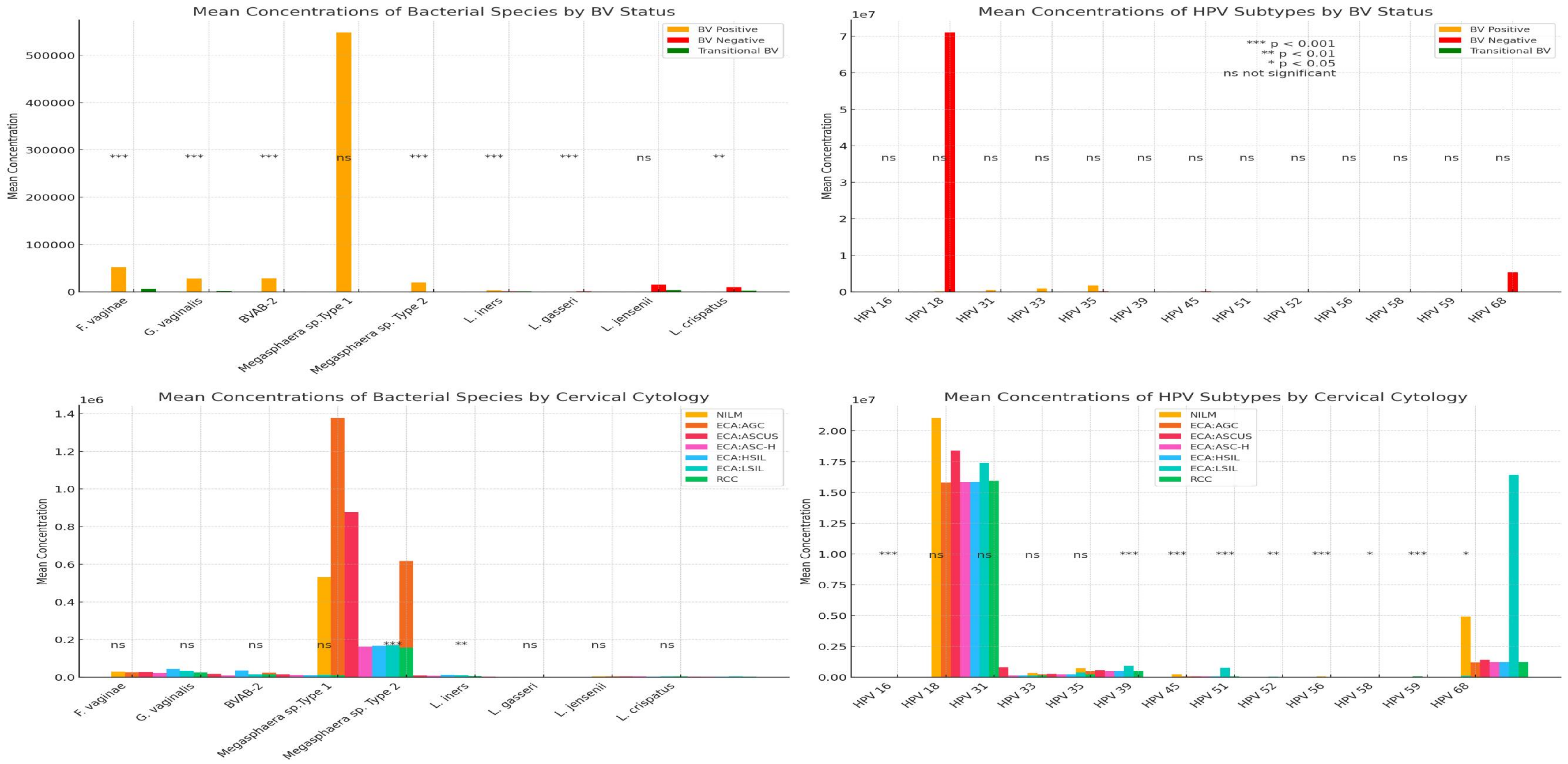


**Fig. S10. Mean concentration of bacterial species and HPV subtypes found in BV-negative, BV-positive, Transitional BV, NILM, RCC, and other ECA-type samples.** Higher concentrations of *F. vaginae, G. vaginalis, BVAB-2, Megasphaera sp.,* HPV-31, HPV-33, and HPV-35 were associated with BV-positive and ECA-type samples (ASCUS, HSIL, LSIL) while higher concentrations of *L. jensenii, L. crispatus, HPV-18 and HPV-68 were associated with BV-negative samples.* Most HPV subtypes had higher concentrations in ECA-positive samples than in NILM-positive samples.

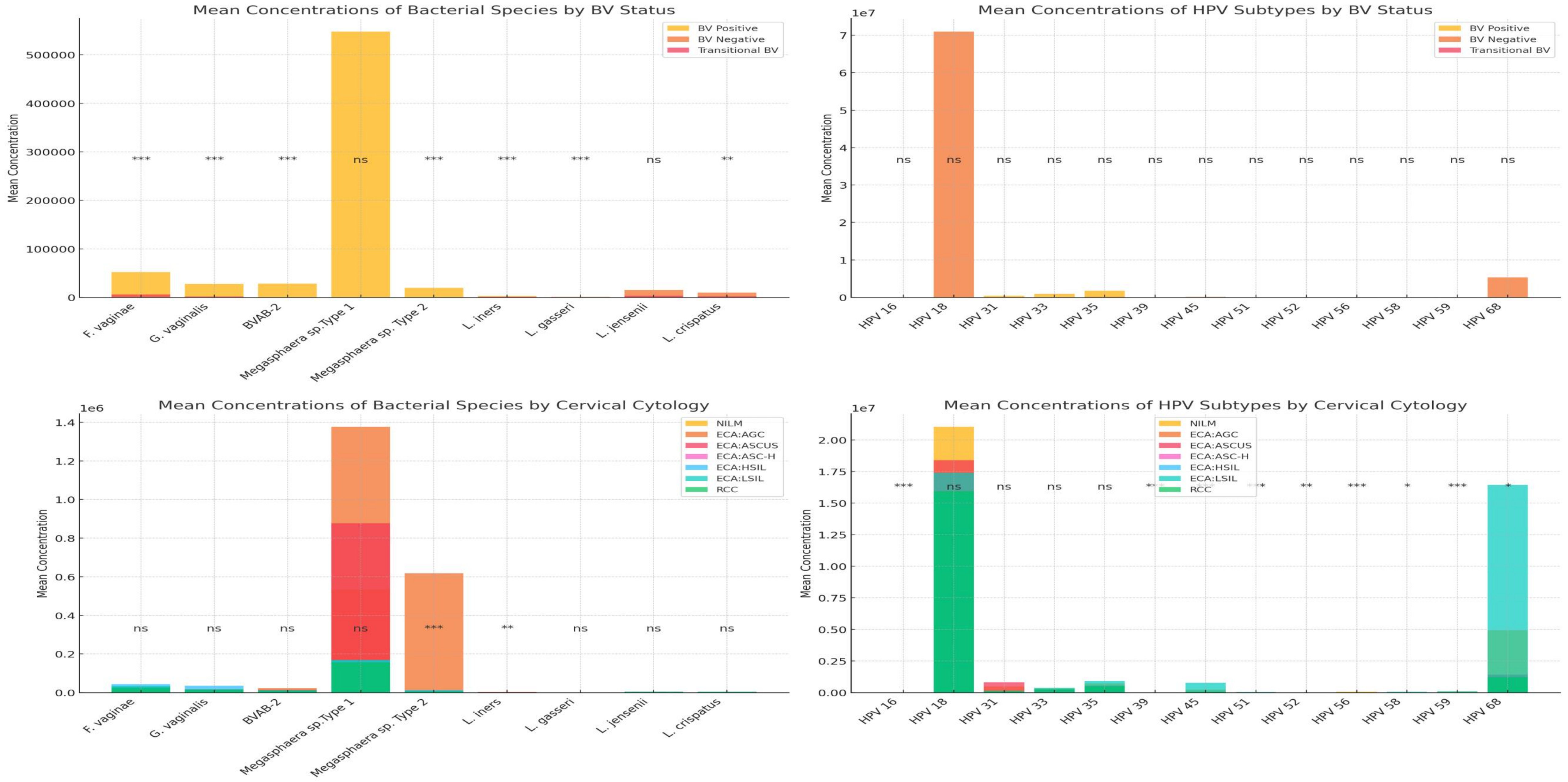


**Fig. S11. Mean concentration of bacterial species and HPV subtypes found in BV-negative, BV-positive, Transitional BV, NILM, RCC, and other ECA-type samples.** Higher concentrations of *F. vaginae, G. vaginalis, BVAB-2, Megasphaera sp.,* HPV-31, HPV-33, and HPV-35 were associated with BV-positive and ECA-type samples (ASCUS, HSIL, LSIL) while higher concentrations of *L. jensenii, L. crispatus, HPV-18 and HPV-68 were associated with BV-negative samples.* Most HPV subtypes had higher concentrations in ECA-positive samples than in NILM-positive samples.

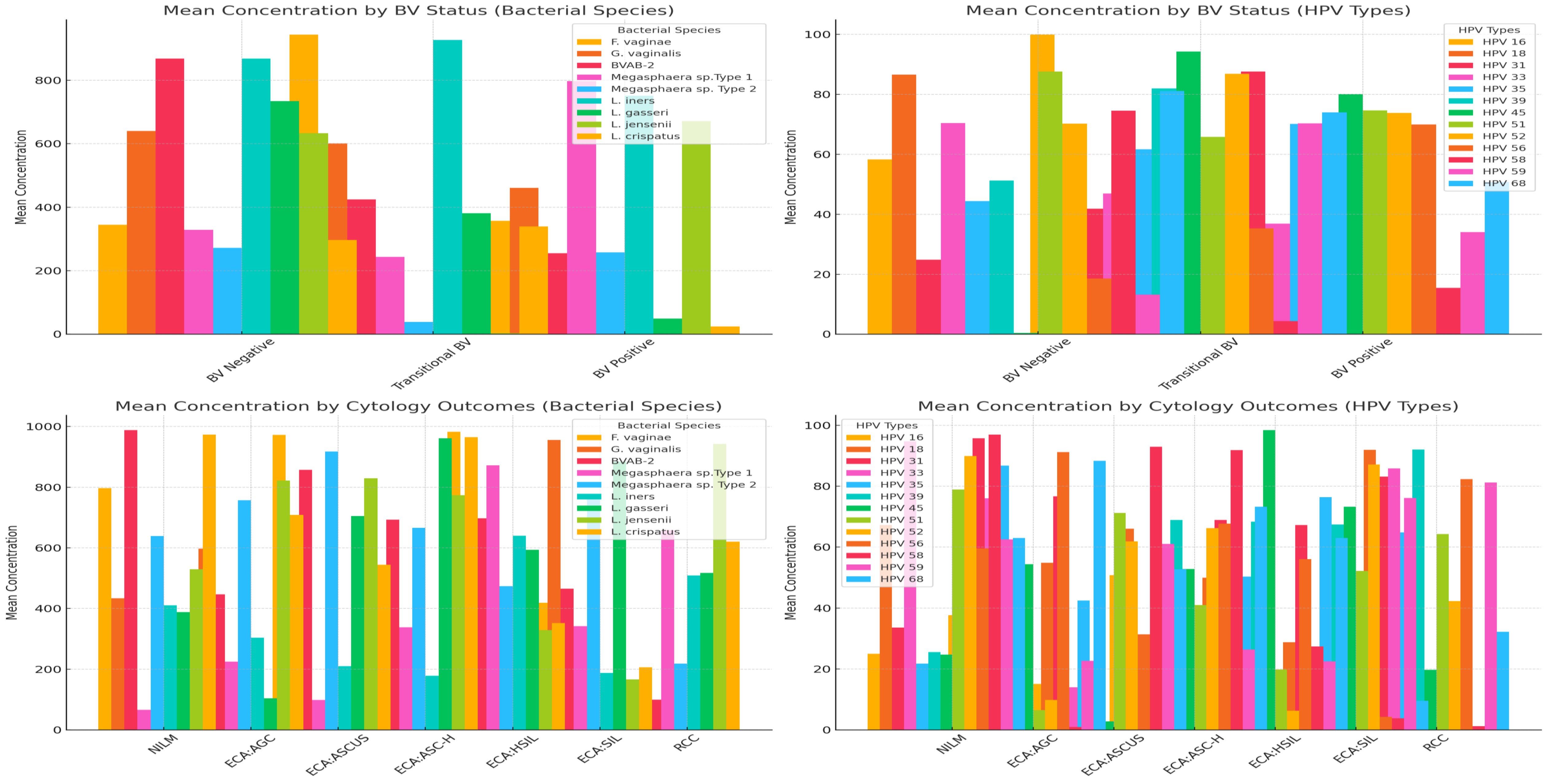


**Fig. S12. Mean concentration of bacterial species and HPV subtypes found in BV-negative, BV-positive, Transitional BV, NILM, RCC, and other ECA-type samples.** Higher concentrations of *F. vaginae, G. vaginalis, BVAB-2, Megasphaera sp.,* HPV-31, HPV-33, and HPV-35 were associated with BV-positive and ECA-type samples (ASCUS, HSIL, LSIL) while higher concentrations of *L. jensenii, L. crispatus, HPV-18 and HPV-68 were associated with BV-negative samples.* Most HPV subtypes had higher concentrations in ECA-positive samples than in NILM-positive samples.

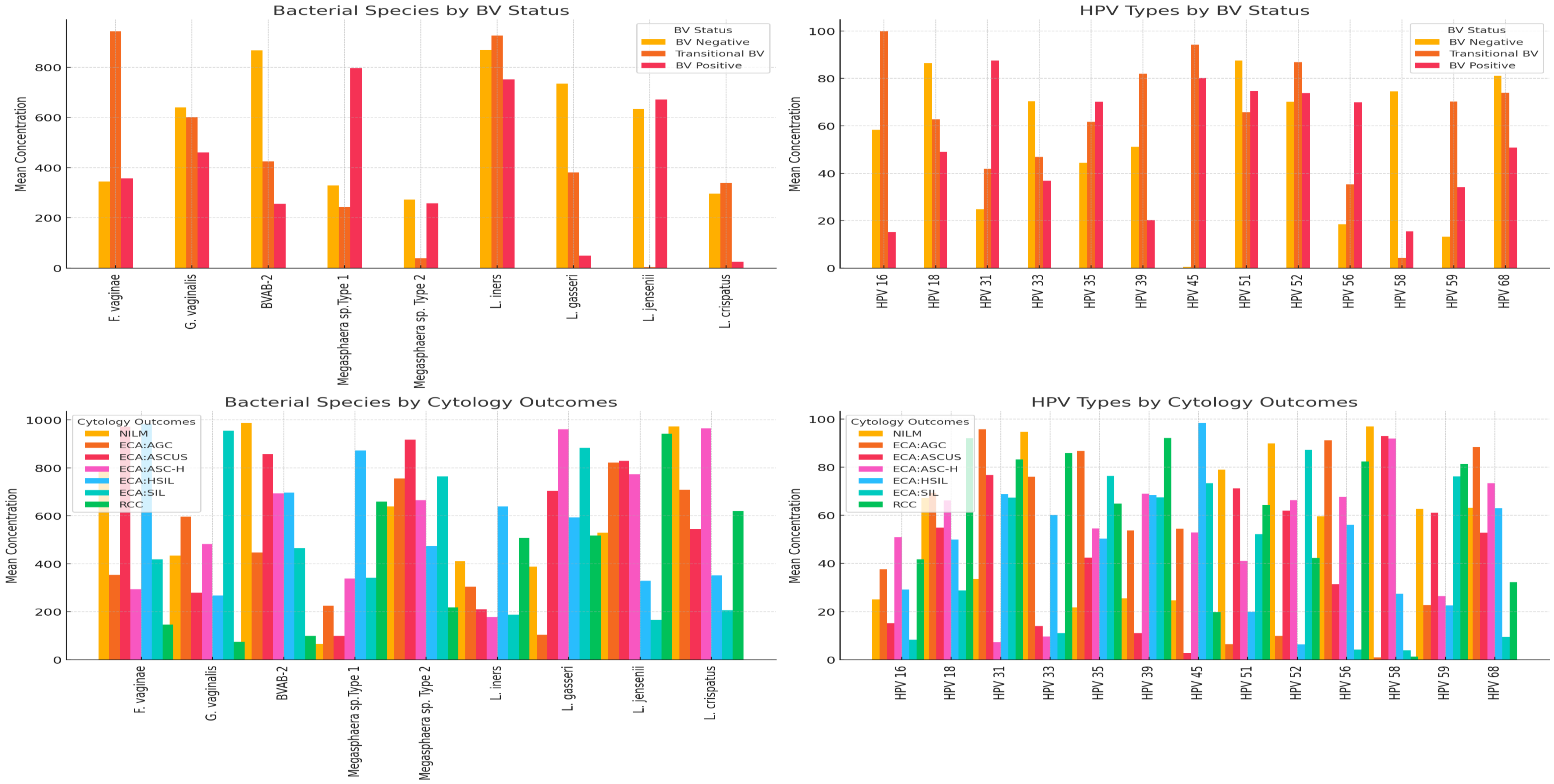


**Fig. S13. Mean concentration of bacterial species and HPV subtypes found in BV-negative, BV-positive, Transitional BV, NILM, RCC, and other ECA-type samples.** Higher concentrations of *F. vaginae, G. vaginalis, BVAB-2, Megasphaera sp.,* HPV-31, HPV-33, and HPV-35 were associated with BV-positive and ECA-type samples (ASCUS, HSIL, LSIL) while higher concentrations of *L. jensenii, L. crispatus, HPV-18 and HPV-68 were associated with BV-negative samples.* Most HPV subtypes had higher concentrations in ECA-positive samples than in NILM-positive samples.

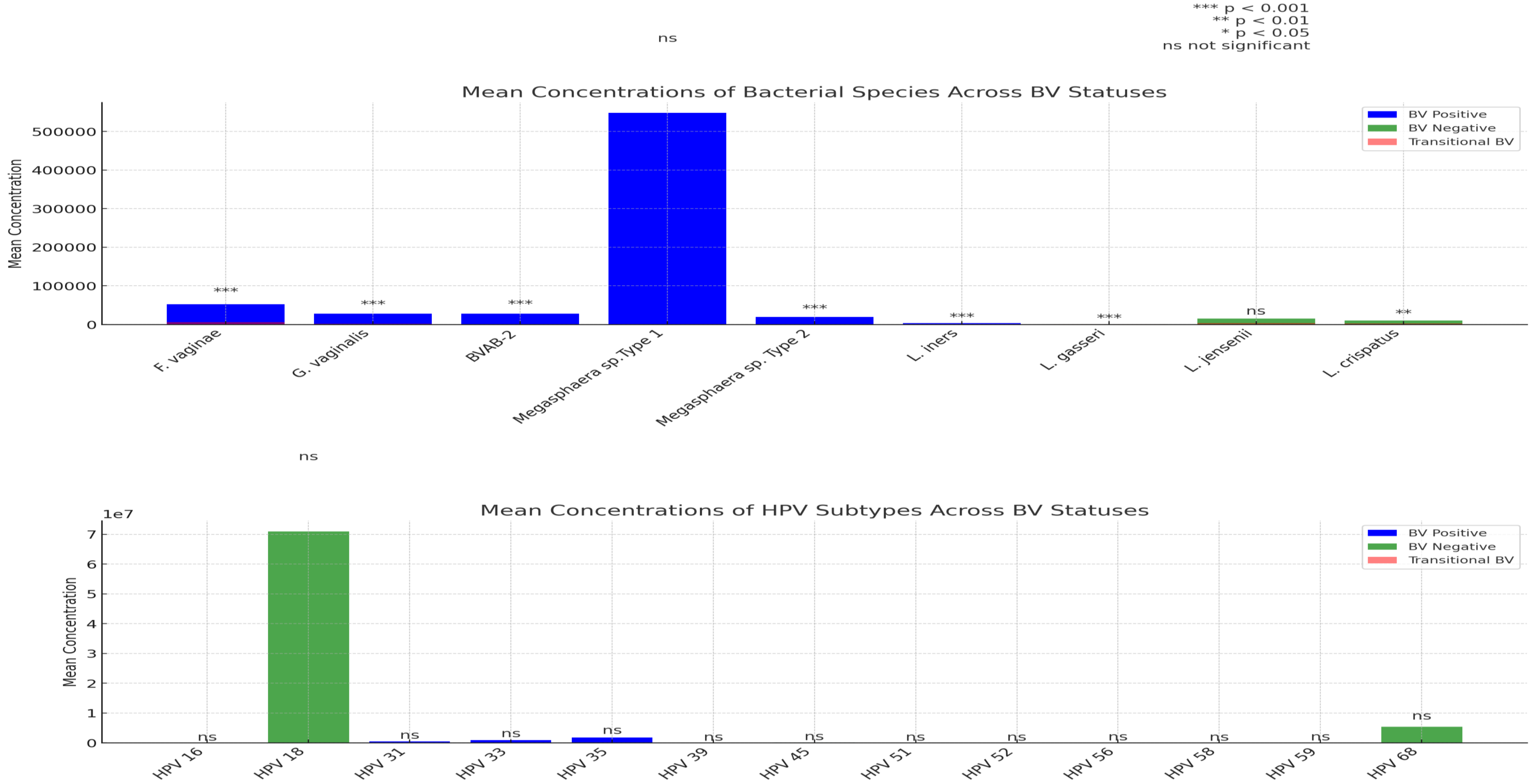


**Fig. S14. Mean concentration of bacterial species found in BV-negative, BV-positive, and Transitional BV samples.** Higher concentrations of *F. vaginae, G. vaginalis, BVAB-2, Megasphaera sp.,* HPV-31, HPV-33, and HPV-35 were associated with BV-positive samples while higher concentrations of *L. jensenii, L. crispatus, HPV-18 and HPV-68 were associated with BV-negative samples.*

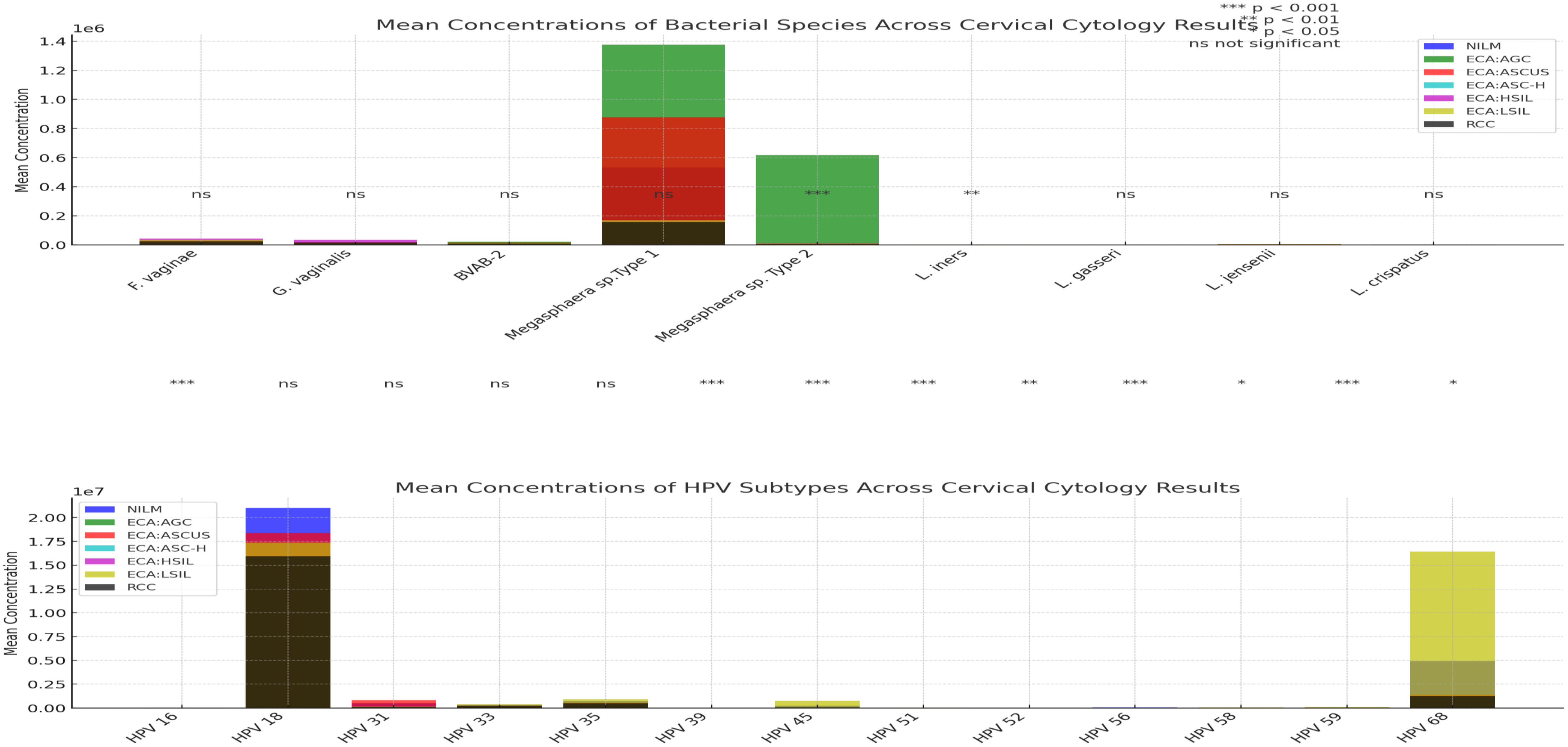


**Fig. S15. Mean concentration of bacterial species and HPV subtypes found in NILM, RCC, and other ECA-type samples.** Higher concentrations of *F. vaginae, G. vaginalis, BVAB-2, Megasphaera sp.*, HPV-31, HPV-33, and HPV-35 were associated with ECA-type samples (ASCUS, HSIL, LSIL). Most HPV subtypes had higher concentrations in ECA-positive samples than in NILM-positive samples.

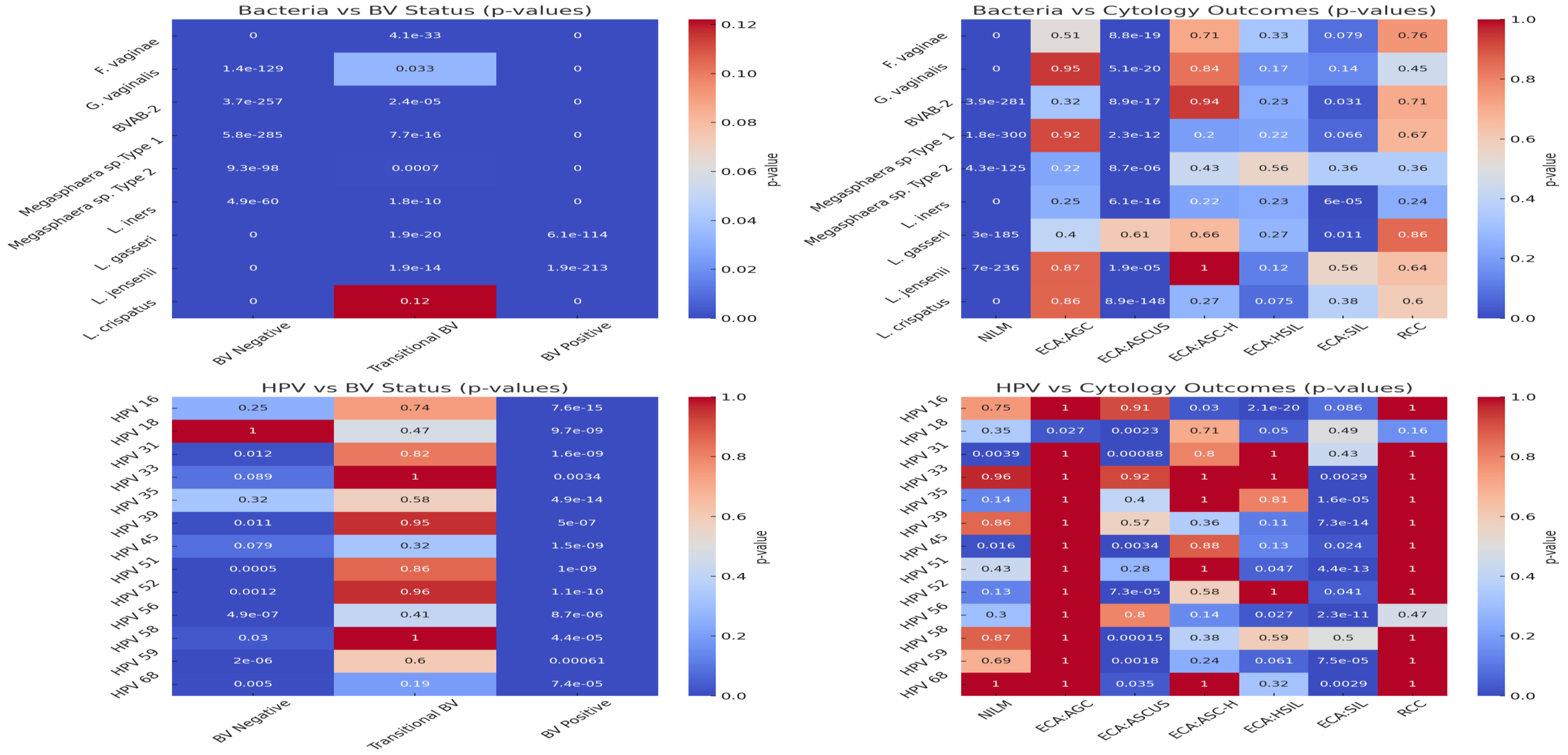


**Fig. S16. Heatmap of p-values showing the significance of the association between the bacterial species and HPV subtypes' frequencies, and BV-negative, BV-positive, Transitional BV, NILM, RCC, and ECA-positive samples**. While most bacteria were significantly associated with BV (except *L. crispatus* and Transitional BV) and NILM, there were significant between fewer species and ASCUS and SIL. Most HPV subtypes were significantly associated with BV-negative and BV-positive, ASCUS, HSIL, and SIL-positive samples.

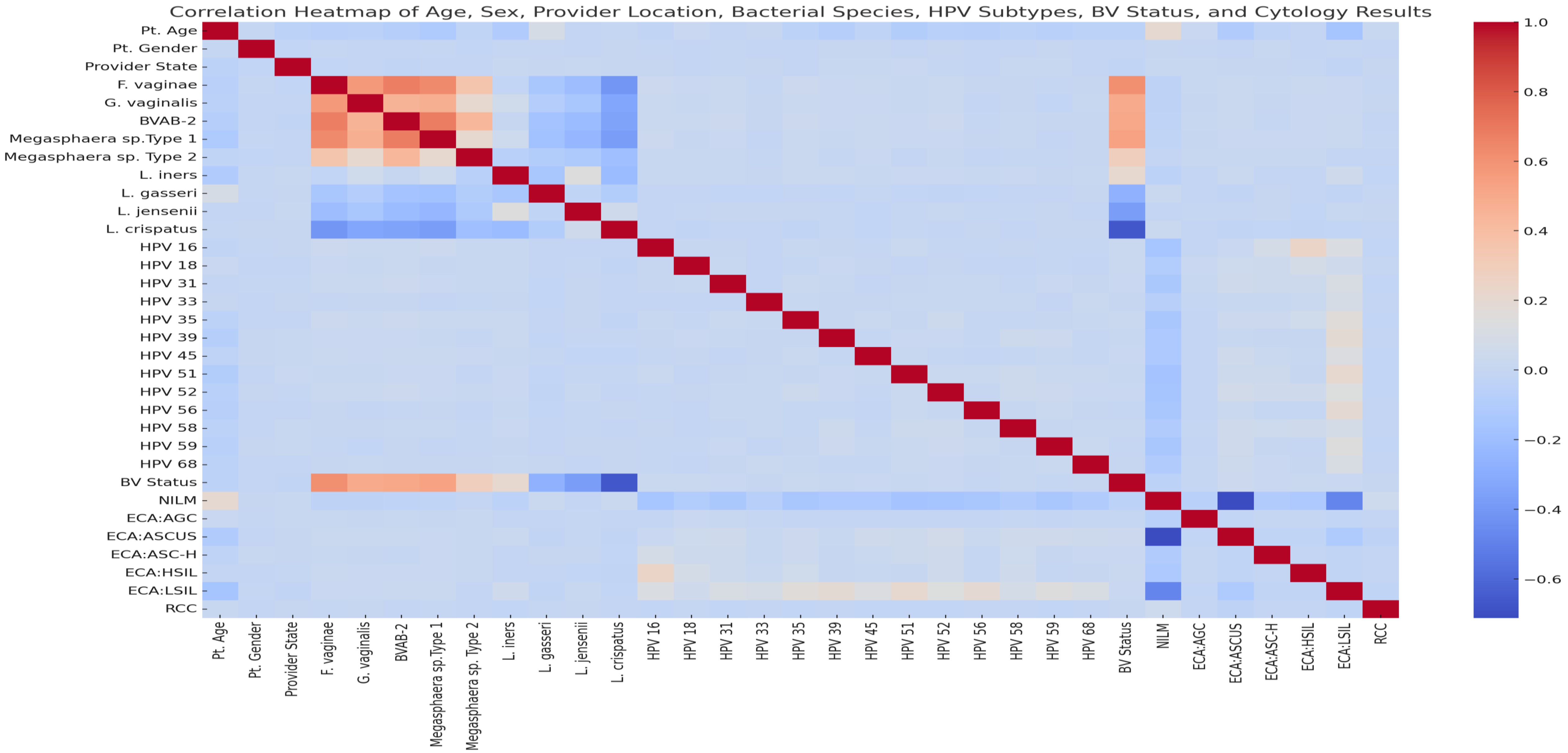


**Fig. S17. Correlation matrix showing the correlation between the demographics, bacterial species and HPV subtypes prevalence, and BV and cervical cytology outcomes**. The heatmap provides insights into how different variables (age, sex, provider location, bacterial species, HPV subtypes, BV status, and cervical cytology outcomes) are correlated with each other. Dark red indicates a strong positive correlation (closer to +1). Dark blue indicates a strong negative correlation (closer to -1). Lighter colors (light blue or light red) represent weaker correlations (closer to 0). The diagonal is dark red because any variable is perfectly correlated with itself (correlation = 1).

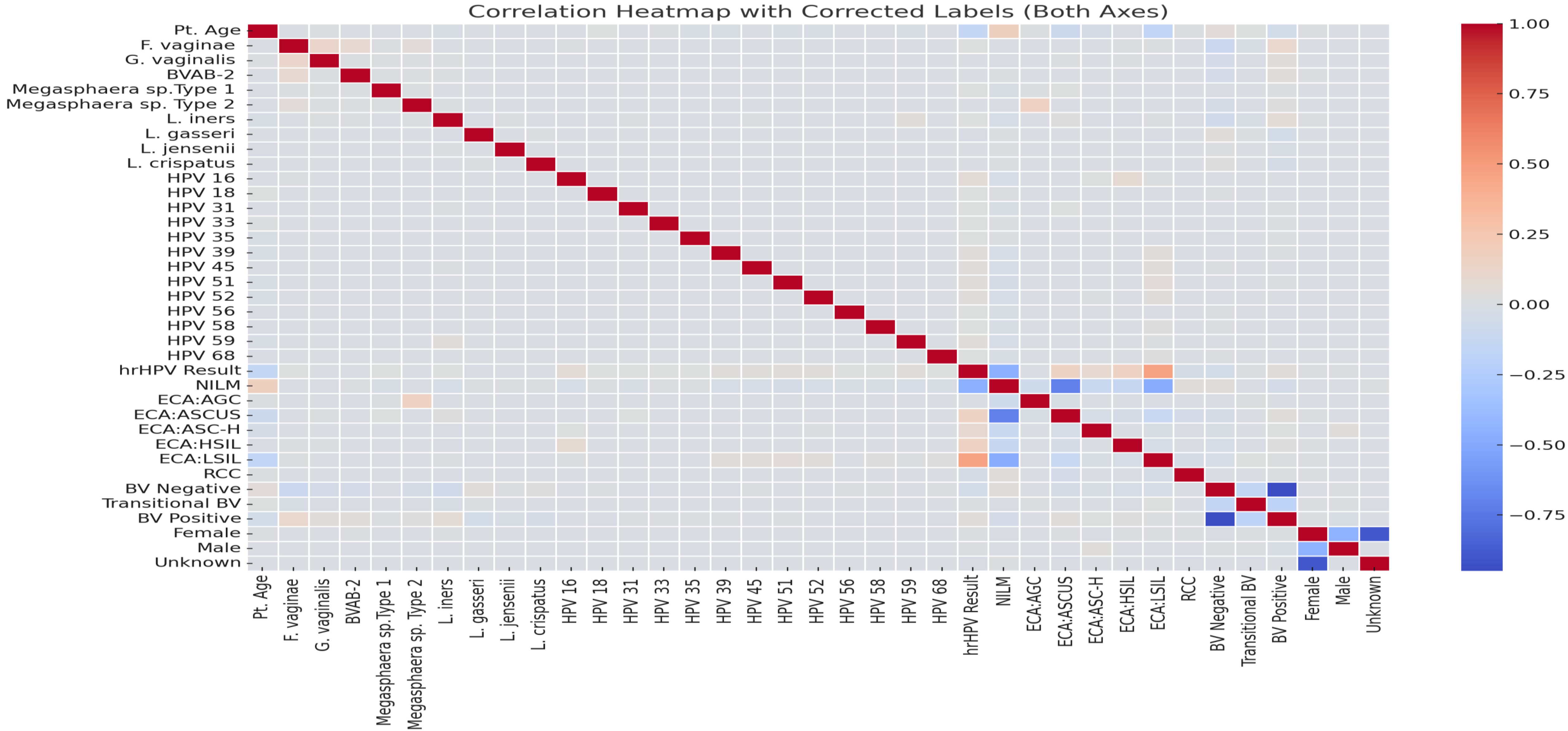


**Fig. S17b. Expanded correlation matrix showing the correlation between the demographics, bacterial species and HPV subtypes prevalence, and BV and cervical cytology outcomes**. The heatmap provides insights into how different variables (age, sex, provider location, bacterial species, HPV subtypes, BV status, and cervical cytology outcomes) are correlated with each other. Dark red indicates a strong positive correlation (closer to +1). Dark blue indicates a strong negative correlation (closer to -1). Lighter colors (light blue or light red) represent weaker correlations (closer to 0). The diagonal is dark red because any variable is perfectly correlated with itself (correlation = 1). Gender and BV is expanded to show their individual components (male, female, unknown, BV positive, BV negative, and Transitional BV)

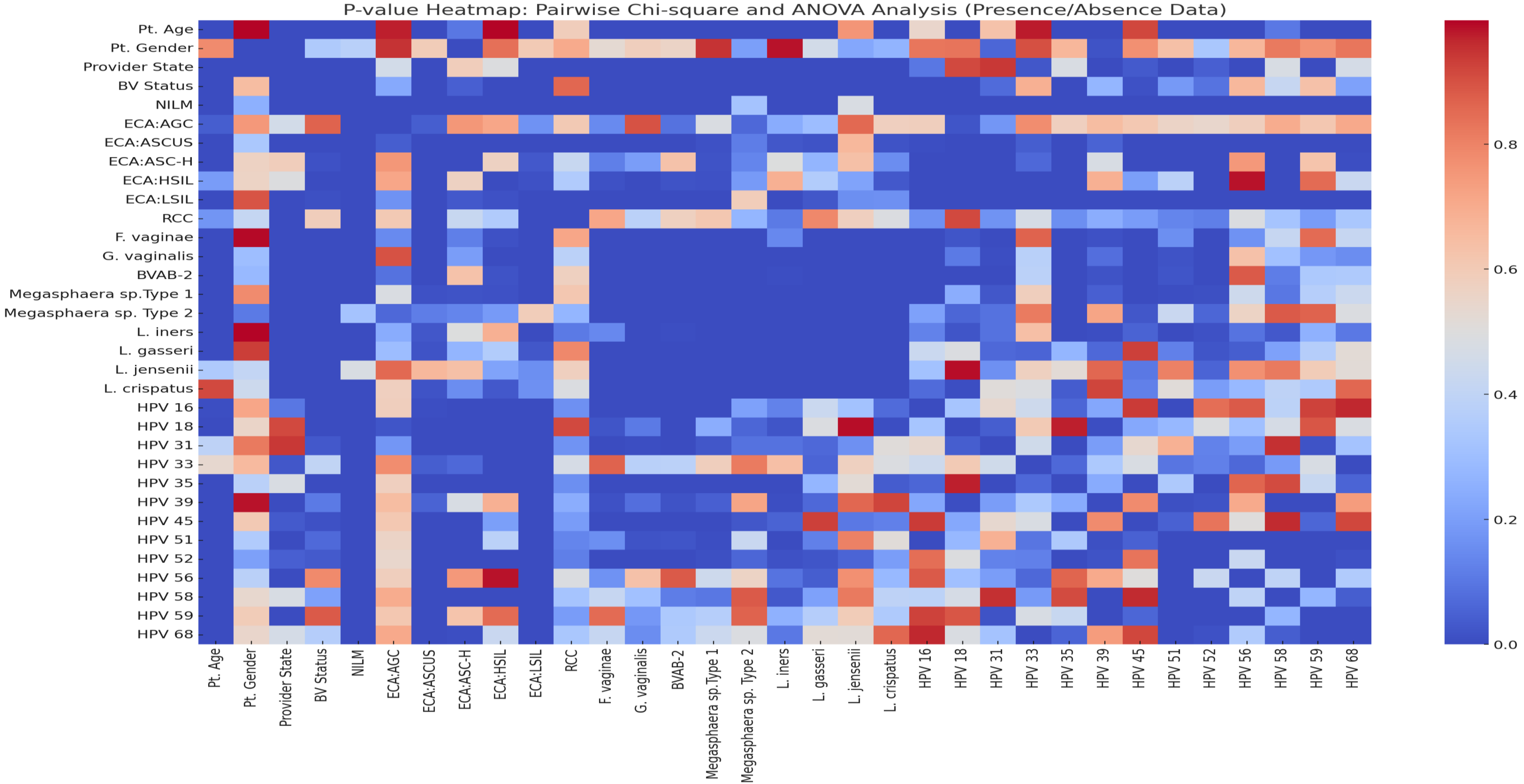


**Fig. S18. A chi-square and ANOVA analysis heatmap plot showing significant and non-significant associations between the various demographics, BV and cervical cytology outcome, bacterial species and HPV subtypes.** Categorical variables were analysed using Chi-square while non-categorical ones such as concentration values of HPV and bacteria species were analysed using ANOVA.

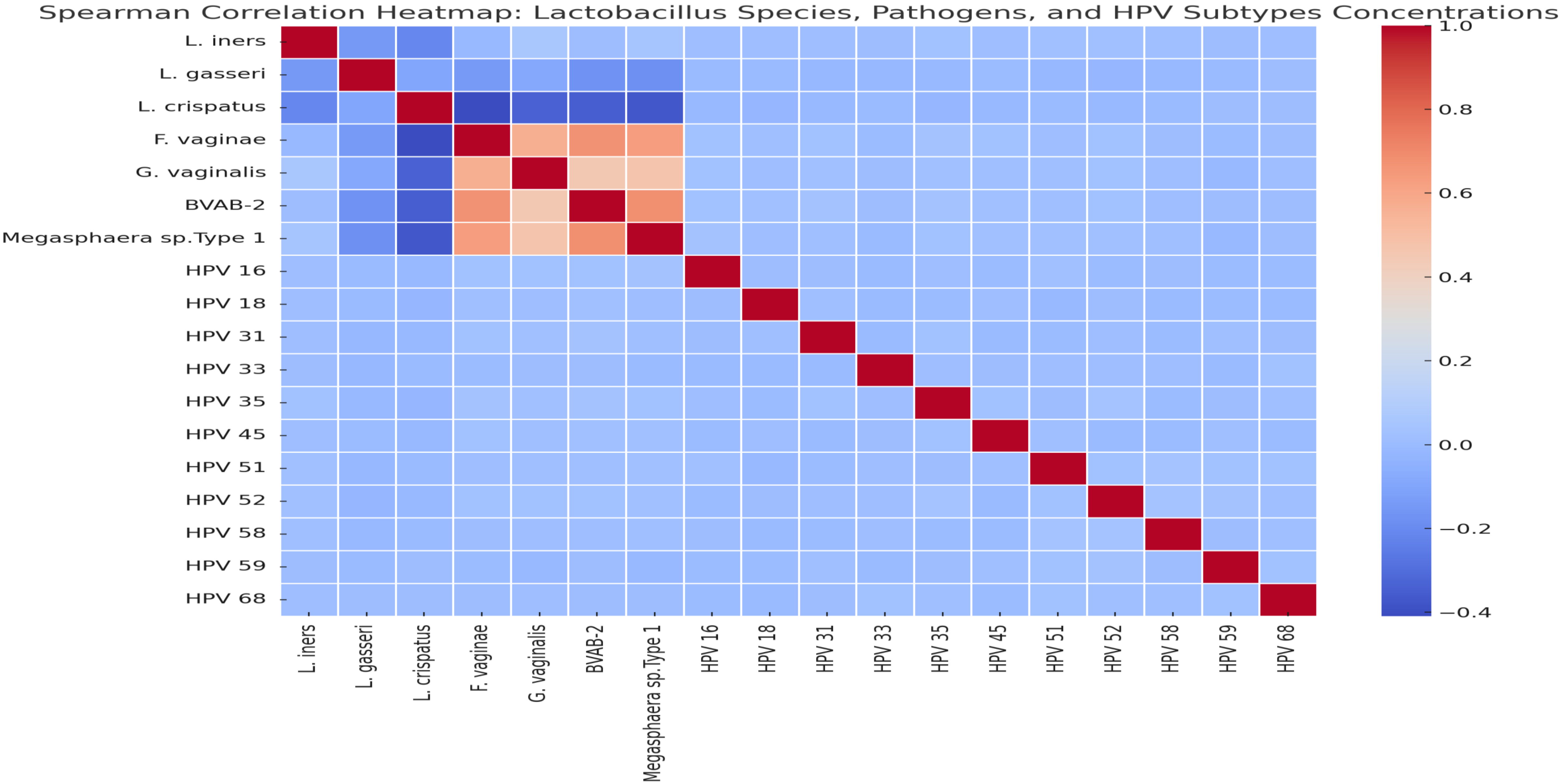


**Fig. S19. A Spearman correlation matrix that shows the correlation between *Lactobacillus and non-lactobacillus bacteria species* and HPV sub types in terms of their concentrations.**

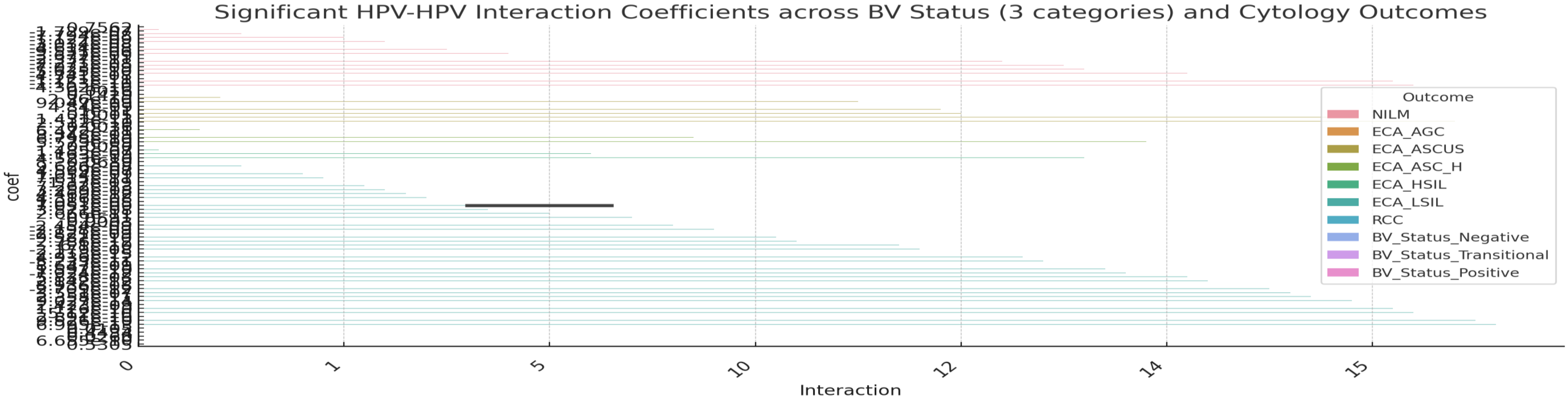


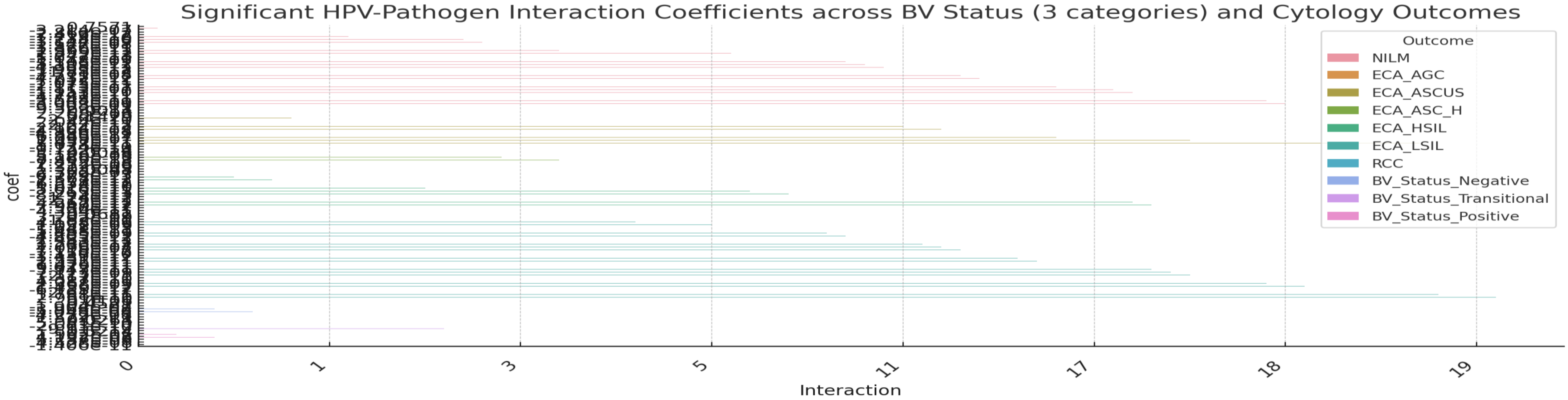


**Fig. S20. HPV-HPV and HPV-bacterial pathogens interactions effects on cervical cytology outcomes.** The upper plot is the co-efficient of the interactions showing the strength and direction of the interaction effects.

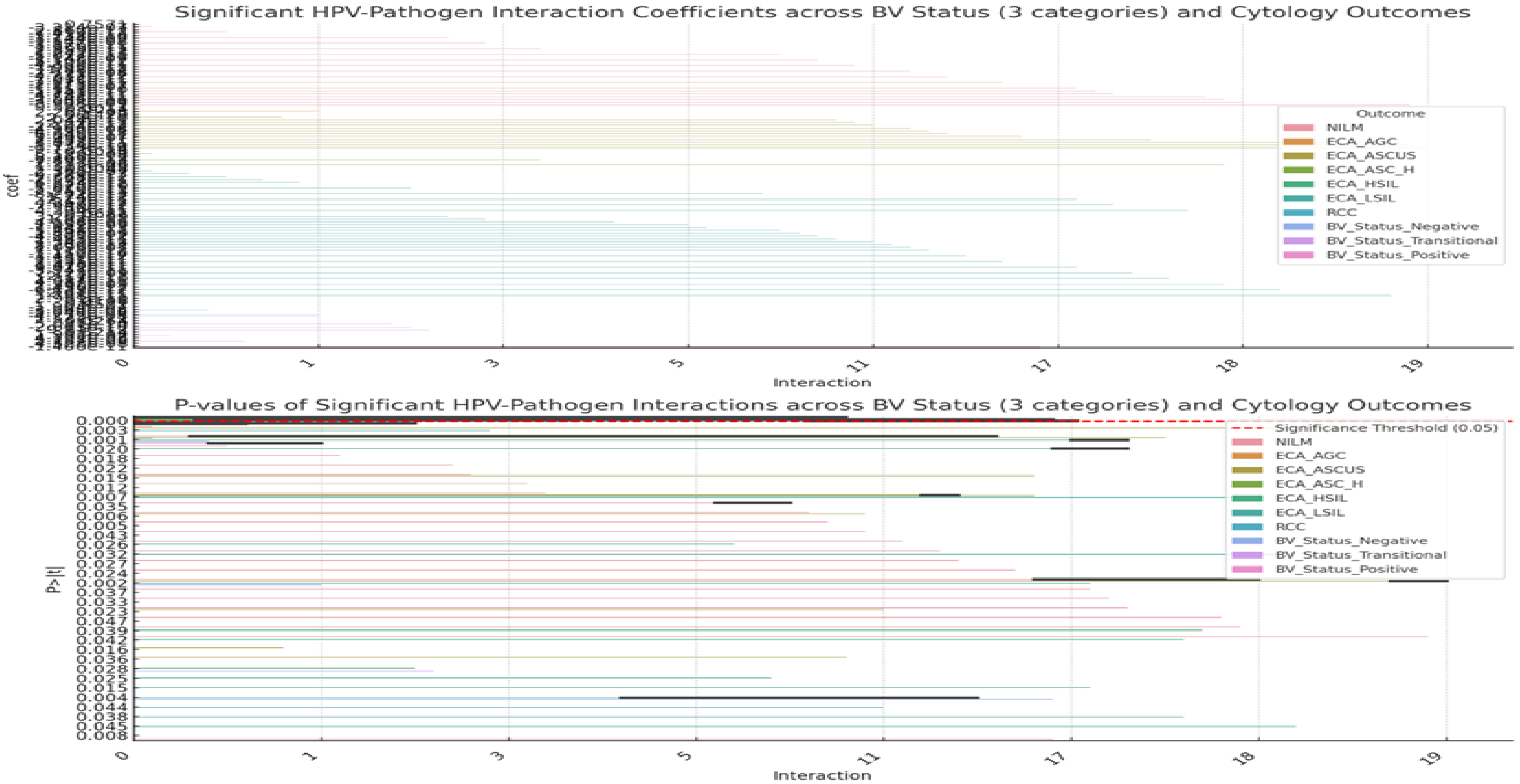


**Fig. S21. HPV-bacterial pathogens interactions effects on BV outcomes.** The upper plot is the co-efficient of the interactions showing the strength and direction of the interaction effects and the lower plot shows the significance (p-values) of the interactions.

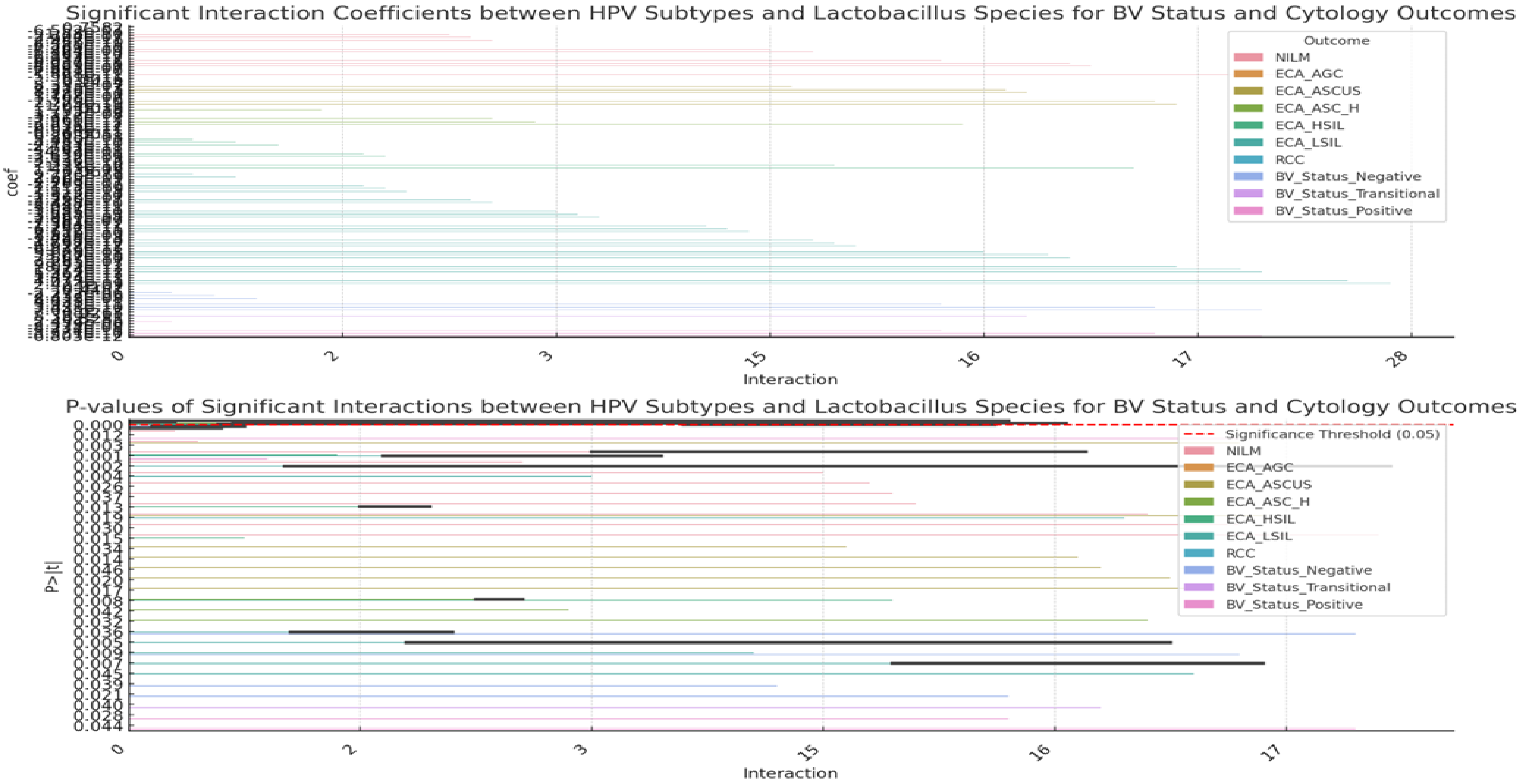


**Fig. S22. HPV-Lactobacillus species interactions effects on BV outcomes.** The upper plot is the co-efficient of the interactions showing the strength and direction of the interaction effects and the lower plot shows the significance (p-values) of the interactions.

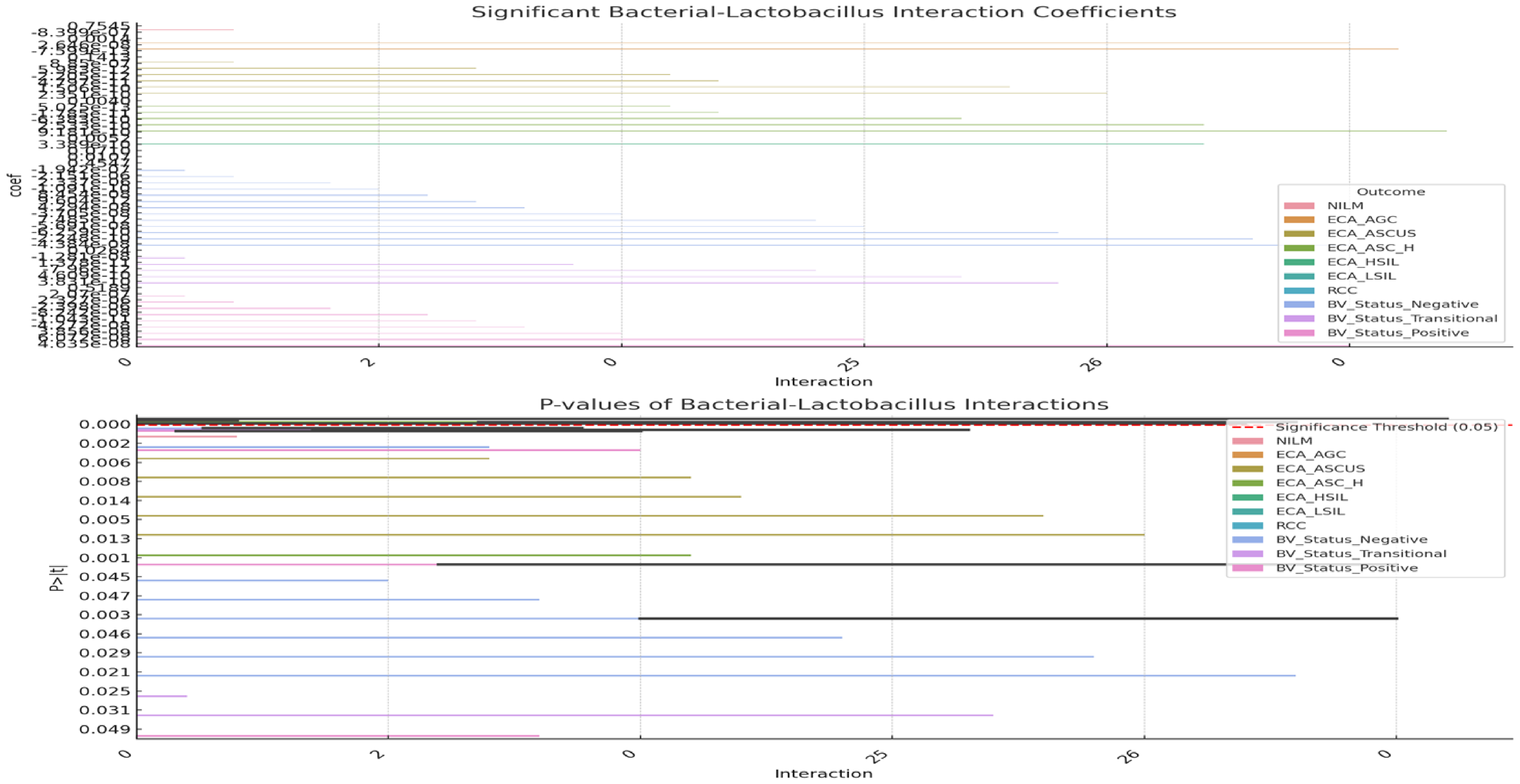


**Fig. S23. lactobacillus-bacterial pathogens interactions effects on BV outcomes.** The upper plot is the co-efficient of the interactions showing the strength and direction of the interaction effects and the lower plot shows the significance (p-values) of the interactions.

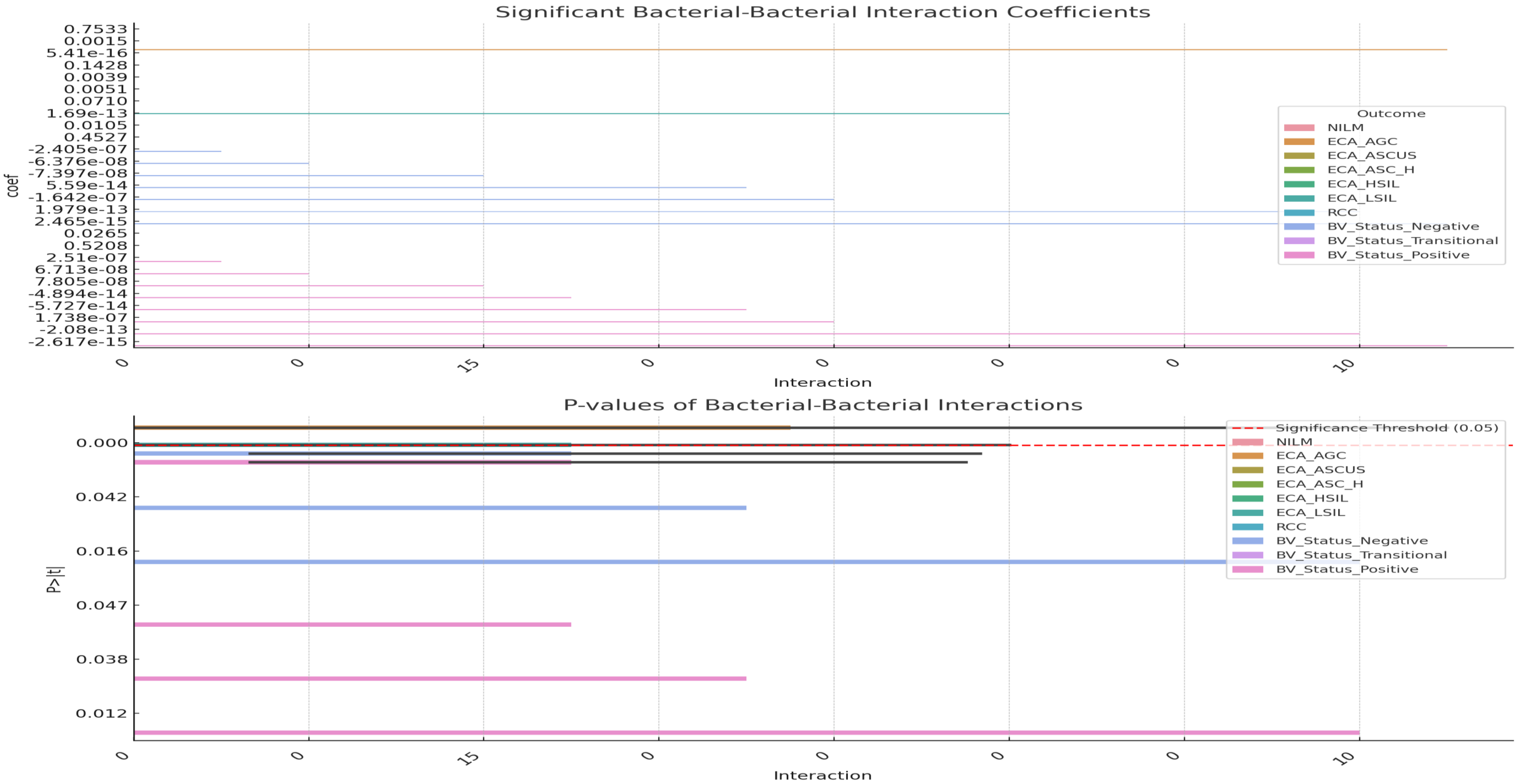


**Fig. S24. Bacteria-bacterial pathogens interactions effects on BV outcomes.** The upper plot is the co-efficient of the interactions showing the strength and direction of the interaction effects and the lower plot shows the significance (p-values) of the interactions.

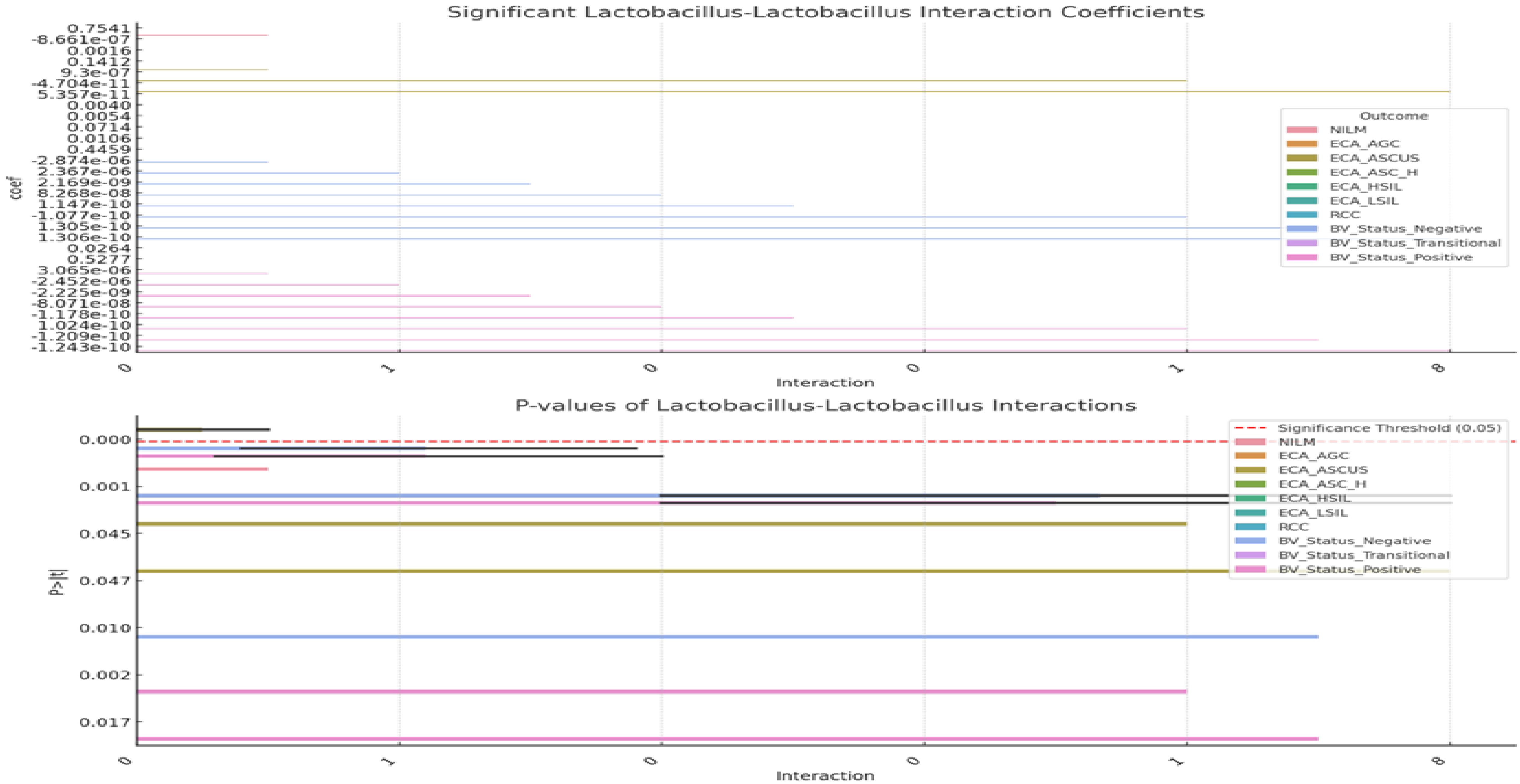


**Fig. S25. lactobacillus-Lactobacillus interactions effects on BV outcomes.** The upper plot is the co-efficient of the interactions showing the strength and direction of the interaction effects and the lower plot shows the significance (p-values) of the interactions.

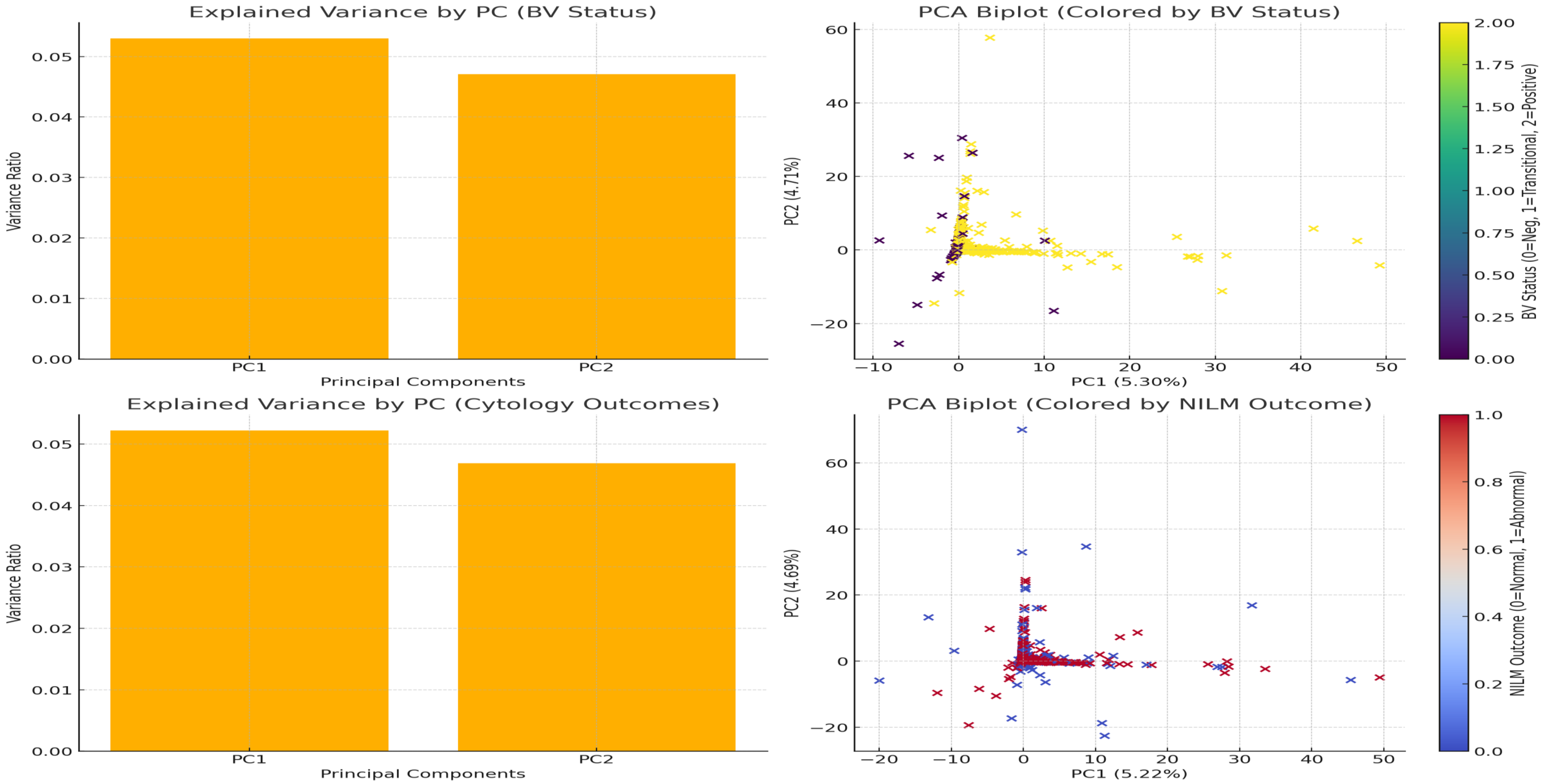


**Fig. S26. Principal component analysis plots showing the variance and biplot of the samples' separation towards the two components. Top Left**: Explained variance by the first two principal components (PC1 and PC2) for the **BV Status** analysis. **Top Right**: PCA biplot colored by **BV Status**, showing how the samples separate based on bacterial species and HPV subtypes. **Bottom Left**: Explained variance for **Cytology Outcomes** (using NILM as an example). **Bottom Right**: PCA biplot colored by **NILM** outcome, illustrating how the data clusters based on this cervical cytology result.

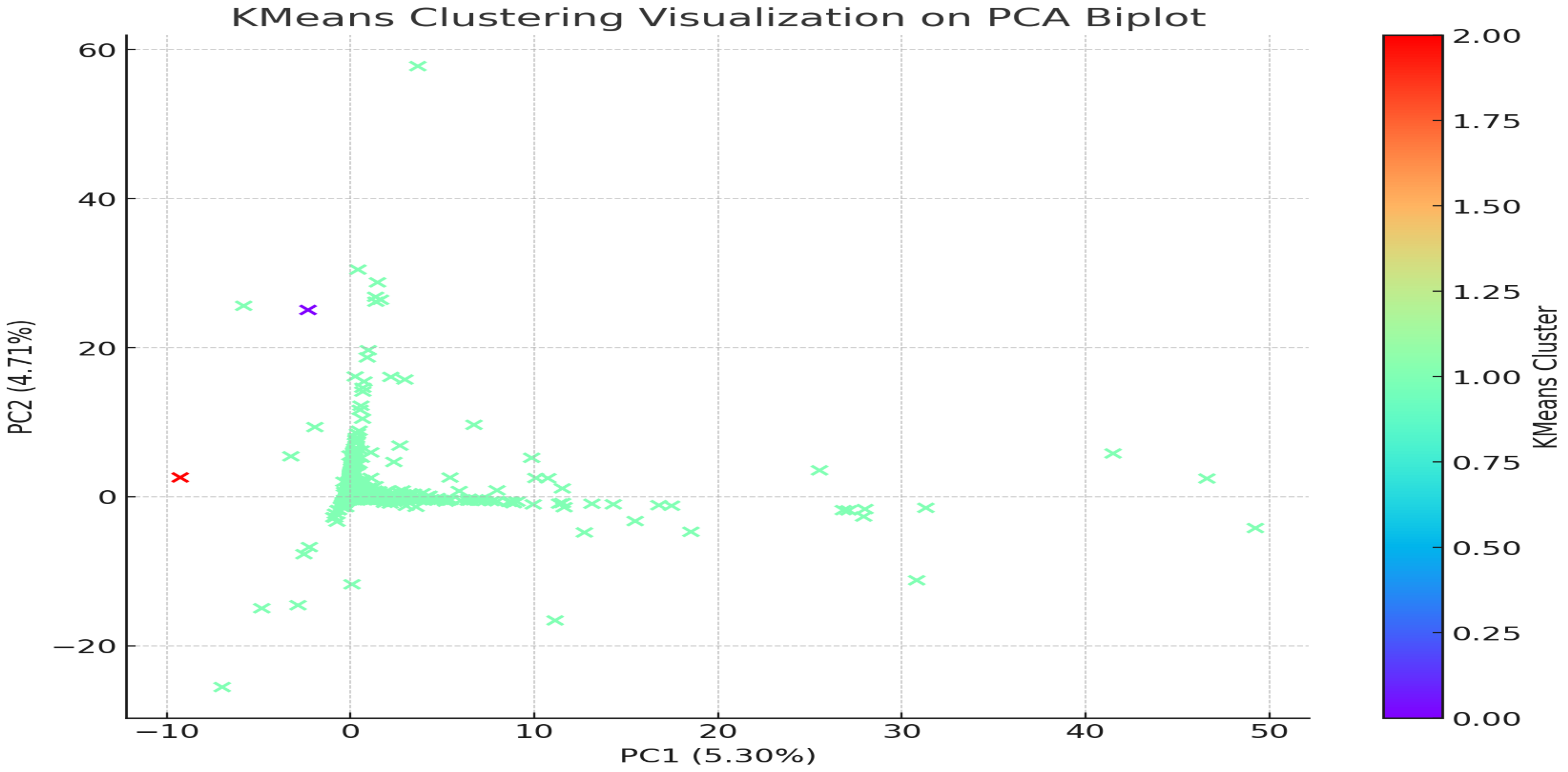


**Fig. S27. K. means clustering data plotted unto two components to show the distribution of the HPV subtypes and bacterial species across the samples.** The red and blue samples (stars) are outliers while the green samples are spread towards the two different components.

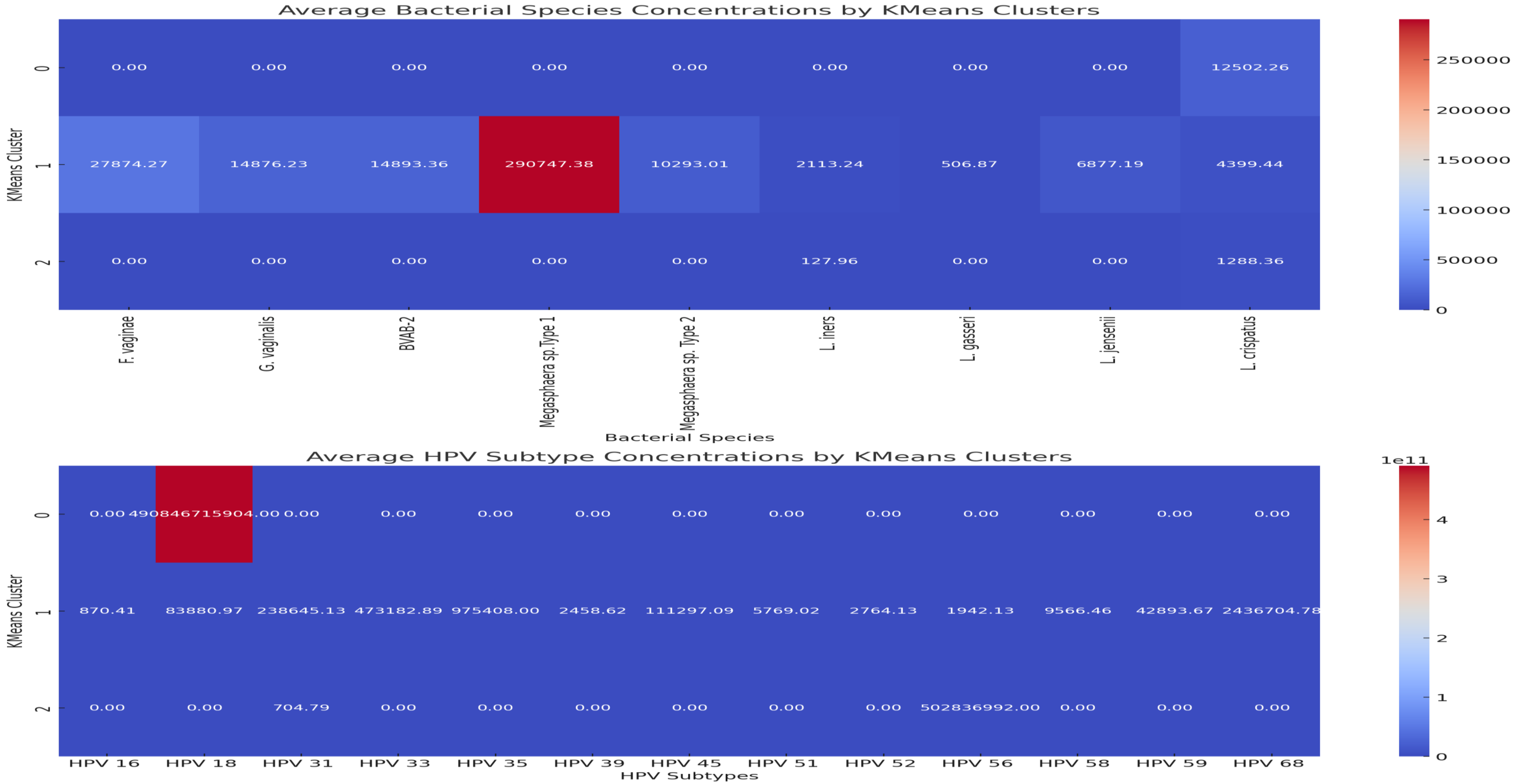


**Fig. S28. A combined heatmap showing the average concentrations of bacterial species and HPV subtypes across the three K-means clusters. Cluster 1 in both cases is associated with BV-positive and ASCUS/HSIL conditions owing to the higher concentrations of bacterial pathogens and HPV subtypes in that cluster.**
