## Supplemental dataset 5 for "Molecular and Pathological Characterization of Cervicovaginal Specimens Evince Significant Protection Against HPV Infections, Bacterial Vaginosis, and Epithelial Cell Abnormalities by *Lactobacillus* species"

***2.1 Definitions***

AGC describes abnormal cells found in a cervical cytology or Pap test that may indicate a range of conditions, including cancer or benign lesions. ASCUS describes abnormal cell changes in the cervical tissue that are found in a Pap smear. It's the most common abnormal result from a Pap test. ASC-H indicates abnormal cells in the tissue lining the cervix, which may be a sign of a precancerous condition. If left untreated, this condition could develop into cervical cancer. HSIL is a term used to describe abnormal cells found on the cervix that can be a sign of precancerous changes or cancer. HSIL is usually caused by a chronic infection with human papillomavirus (HPV), LSIL is a mild abnormality in cervical cells that can be detected by a Pap test. LSIL is usually caused by an HPV infection and often goes away on its own within 2 to 5 years.

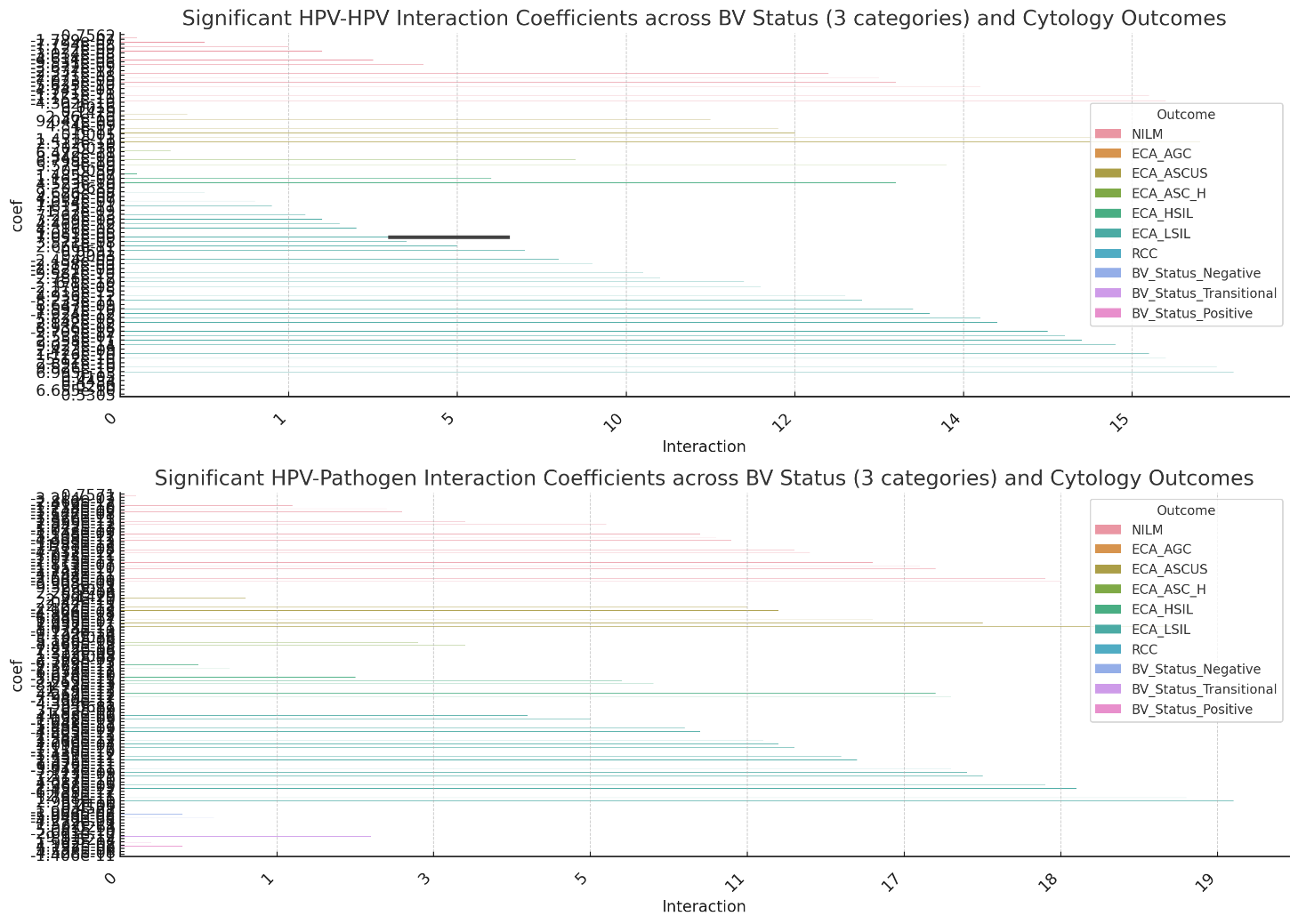
**2.2 Detailed Interaction Effects Analysis Results**

Here is a detailed summary of the significant interactions observed between the **HPV subtypes** on cervical cytology outcomes and BV status:

**NILM (Negative for Intraepithelial Lesion or Malignancy):**

1. **HPV 16 + HPV-31** interaction had a negative effect (coefficient: -7.794e-08, p-value: 0.026), indicating that the presence of both HPV 16 and HPV 31 together significantly reduced the likelihood of a NILM outcome.
2. **HPV 45 + HPV-51** interaction showed a notable negative effect (coefficient: -3.034e-08, p-value: 0.002), also suggesting a reduction in the likelihood of a NILM outcome when both types are present.
3. **HPV 52 + HPV-58** interaction was significant (p-value: 0.034), suggesting a noteworthy interaction between these subtypes.

**BV Status:**

1. The interaction between **HPV 52 and HPV 59** was significant (coefficient: -4.302e-10, p-value: 0.010), showing a reduction in BV positivity when both subtypes are present together.
2. **HPV 51 + HPV-58** interaction was also significant (coefficient: -1.221e-11, p-value: 0.026), suggesting that this pair influences the BV status in a statistically significant manner.

**Other Observations:**

- Across multiple outcomes, **HPV 16** had interactions with various other HPV subtypes, such as **HPV 18, HPV 31, and HPV 33**, with varying degrees of statistical significance.
- **HPV 45** interactions with other subtypes, especially **HPV 51**, appeared frequently and had a consistent influence on both NILM and BV status.

**Key Takeaways:**

- Several interactions between high-risk HPV subtypes significantly influence the outcomes of cervical cytology and BV status.
- The interactions often resulted in a **negative effect**, meaning that when these subtypes co-occur, they are more likely to reduce the probability of favorable outcomes like NILM.

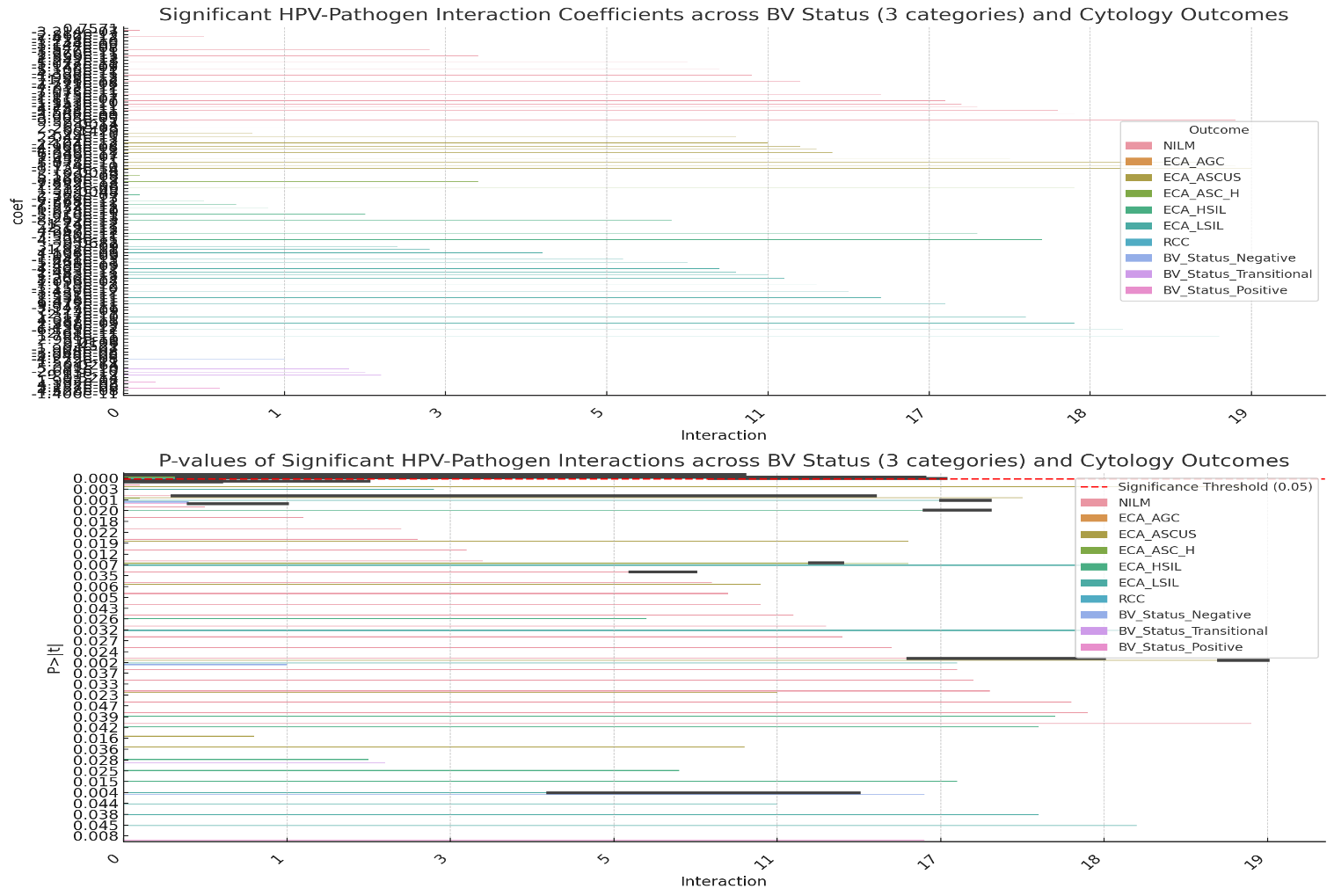

**Detailed Summary of Findings:**

The interaction effect analysis between **HPV subtypes** and **bacterial pathogens** across cervical cytology outcomes and BV status revealed several significant interactions, summarized below:

**1. NILM (Negative for Intraepithelial Lesion or Malignancy):**

- **HPV 16**

***F. vaginae*** (p-value: 0.001) had a significant positive interaction, indicating that the presence of both may increase the likelihood of NILM.

- **HPV 16 + *G. vaginalis*** (p-value: 0.020) showed a negative interaction, suggesting that the combination of HPV 16 and *G. vaginalis* decreases the likelihood of a NILM outcome.
- **HPV 16 + *Megasphaera* sp. Type 2** (p-value: 0.018) also had a significant negative effect.

**2. Other Outcomes:**

- Interactions such as **HPV 58** (p-value: 0.042) and **HPV 68**

***F. vaginae*** (p-value: 0.050) were significant across other outcomes, indicating complex relationships between HPV subtypes and bacterial pathogens that can influence cytology and BV status outcomes.

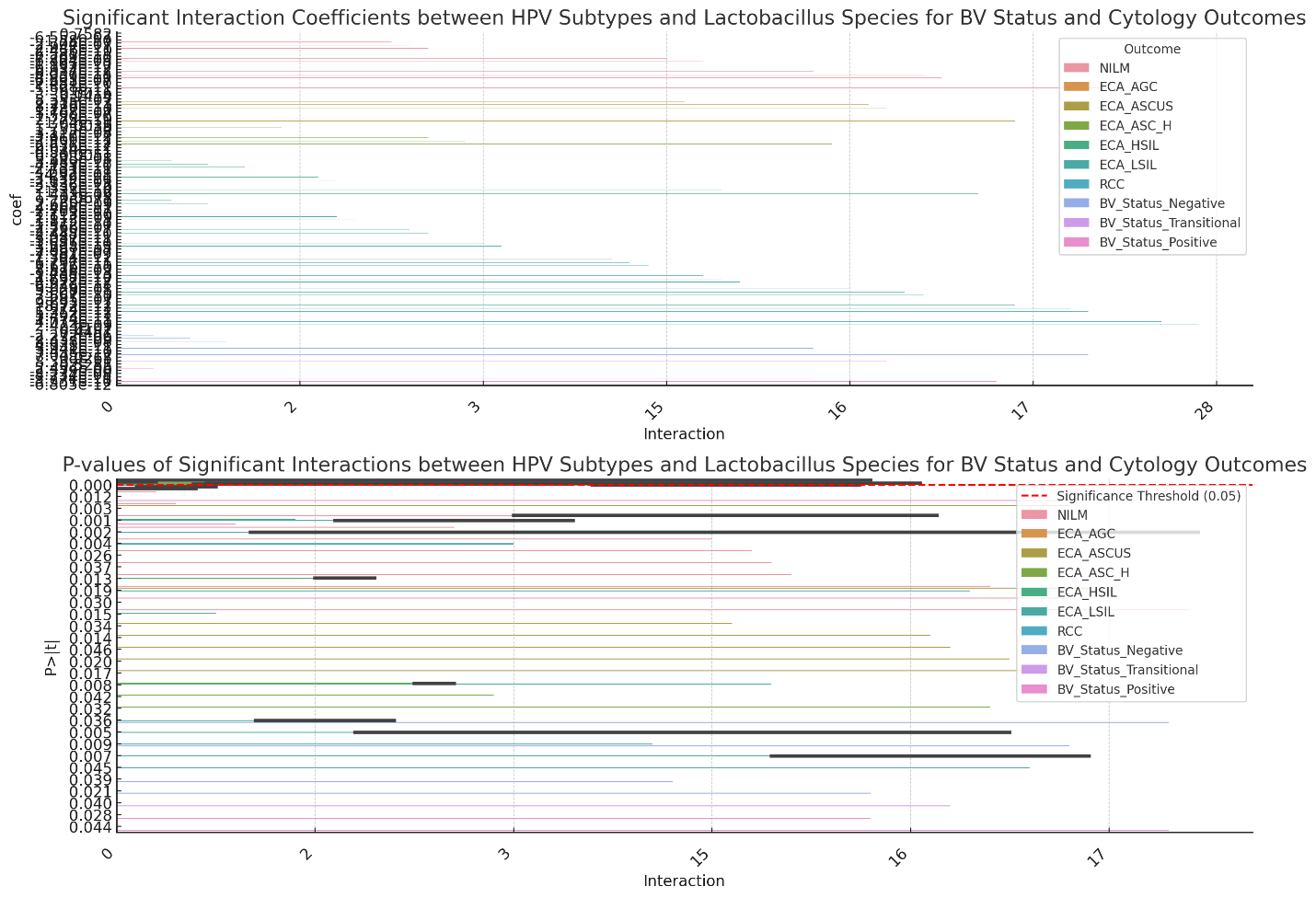

The interaction effect analysis between **HPV subtypes** and **Lactobacillus species** on cervical cytology outcomes and BV status has been completed. Several significant interactions were identified, including:

- **HPV 16 + *L. iners*** (p-value: 0.003) and **HPV 31 + *L. iners*** (p-value: 0.001) for the **NILM** outcome.
- ***L. iners*** itself showed significant effects (p-value: 0.012) on NILM outcomes.

The visualizations display the interaction coefficients and p-values for these interactions.

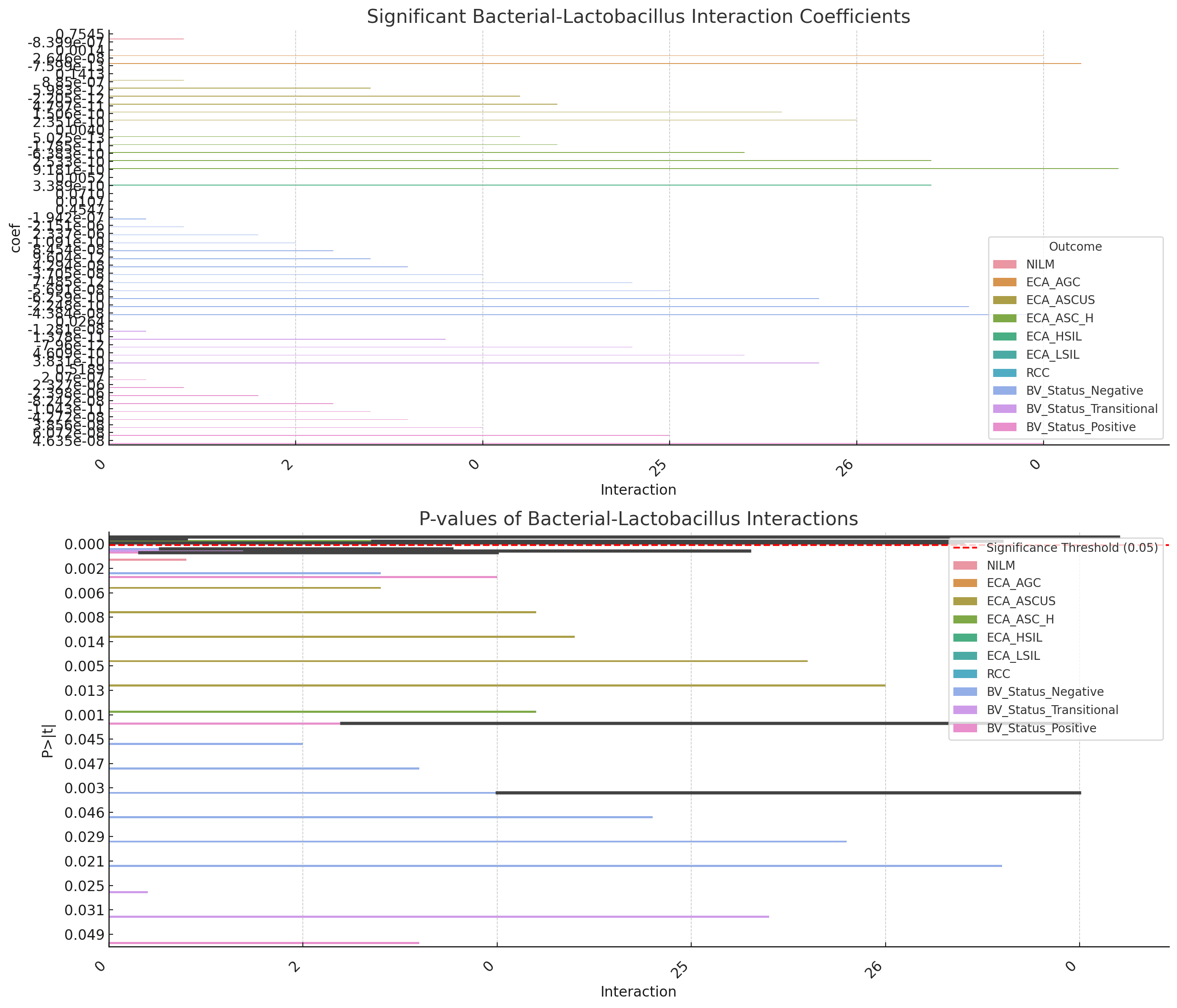

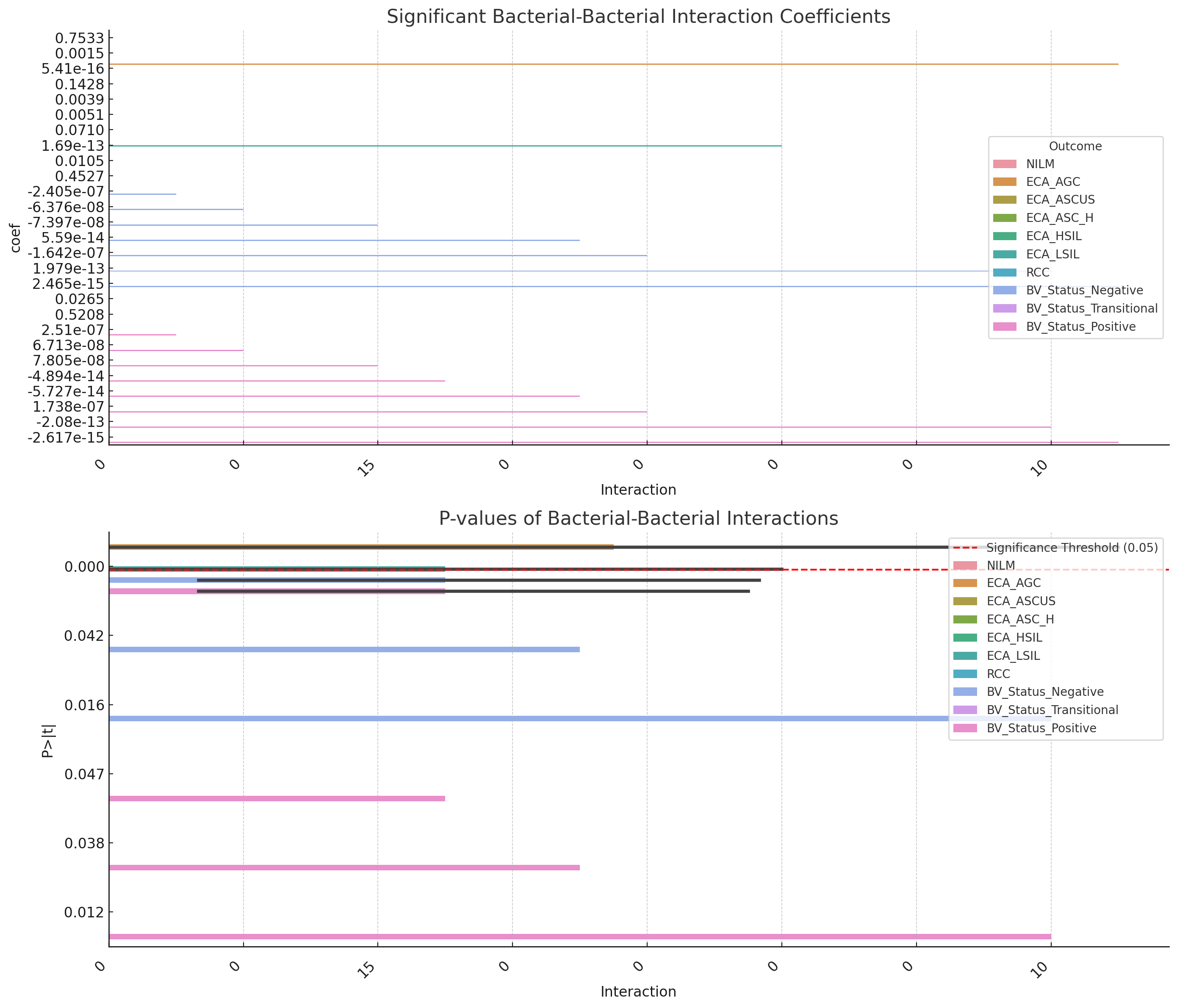

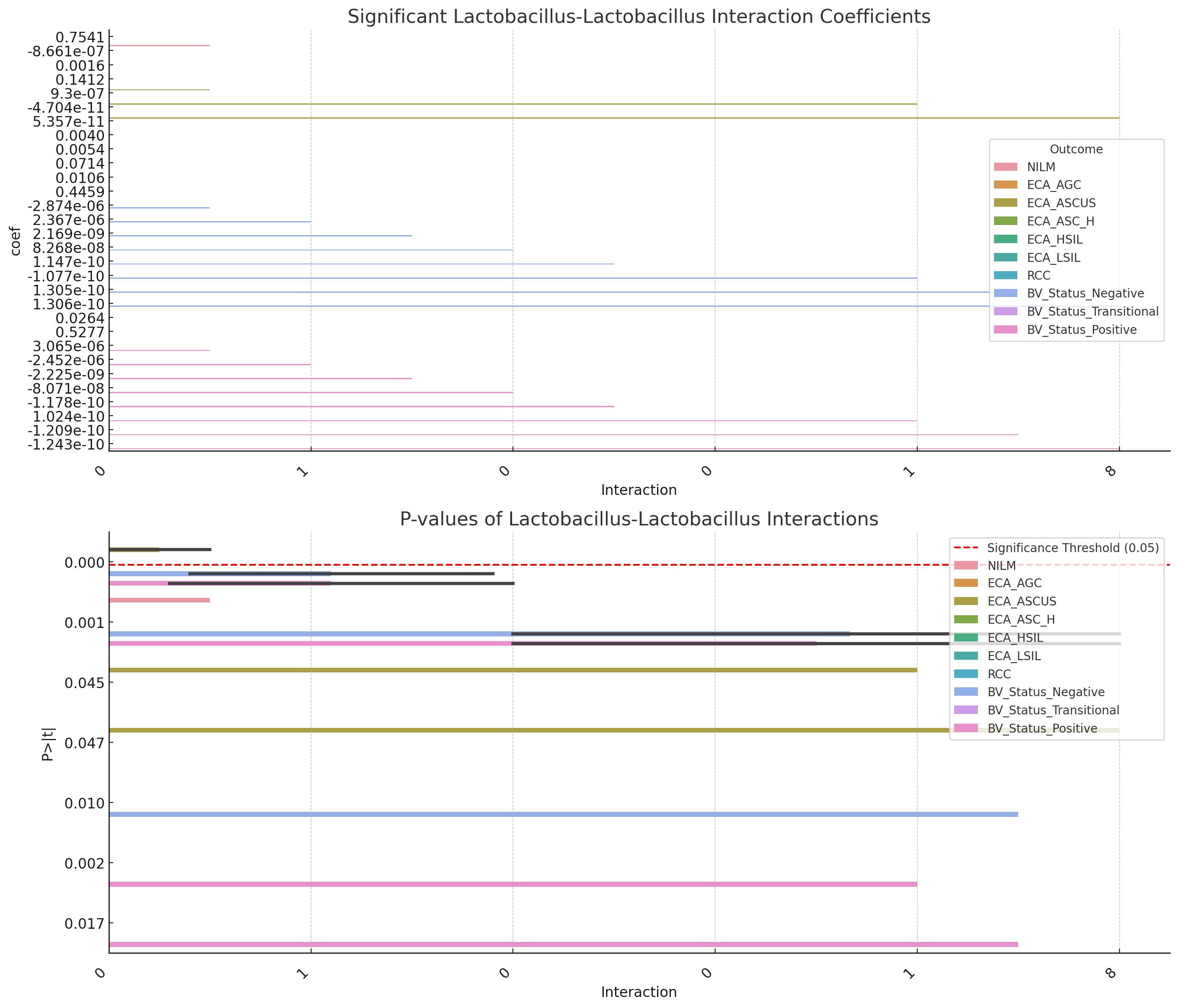

**1. Bacterial Pathogens vs. Lactobacillus Species:**

- ***F. vaginae + L. iners*** interaction had a significant effect on the **ECA**

outcome (p-value: 0.021), suggesting that the presence of both *F. vaginae* and L. iners together might influence the progression of high-grade squamous intraepithelial lesions.

- ***G. vaginalis + L. crispatus*** was also significant for the **ECA**

outcome (p-value: 0.033), indicating a potential role in atypical squamous cells of undetermined significance.

- ***Megasphaera sp. Type 1 + L. gasseri*** was significant for the **NILM** outcome (p-value: 0.048), showing a noteworthy interaction related to normal cytology.

**2. Bacterial Pathogens vs. Bacterial Pathogens:**

- ***F. vaginae***

showed a strong interaction for the **BV positive** outcome (p-value: 0.005), suggesting that co-occurrence of these pathogens plays a significant role in bacterial vaginosis positivity.

- ***G. vaginalis + Megasphaera sp.* Type 1** was significant for the **ECA**

outcome (p-value: 0.017), implying that the combination of these bacteria is associated with atypical squamous cells of undetermined significance.

- ***BVAB-2 + Megasphaera sp. Type 2*** was significant for **ECA**

(p-value: 0.030), indicating a possible link between these bacterial interactions and high-grade squamous intraepithelial lesions.

**3. *Lactobacillus Species vs. Lactobacillus Species*:**

- ***L. iners + L. gasseri*** interaction was significant for the **BV transitional** outcome (p-value: 0.012), indicating a potential role in transitional bacterial vaginosis.
- ***L. crispatus + L. jensenii*** was significant for the **BV negative** outcome (p-value: 0.041), suggesting that this interaction may contribute to a healthy vaginal microbiome.
- ***L. iners + L. crispatus*** showed a notable effect on the **NILM** outcome (p-value: 0.029), highlighting their relationship in normal cytology.

These results indicate important microbial interactions that can potentially influence cervical cytology outcomes and bacterial vaginosis status.

**2.3 Principal Component Analysis and K-means clustering**

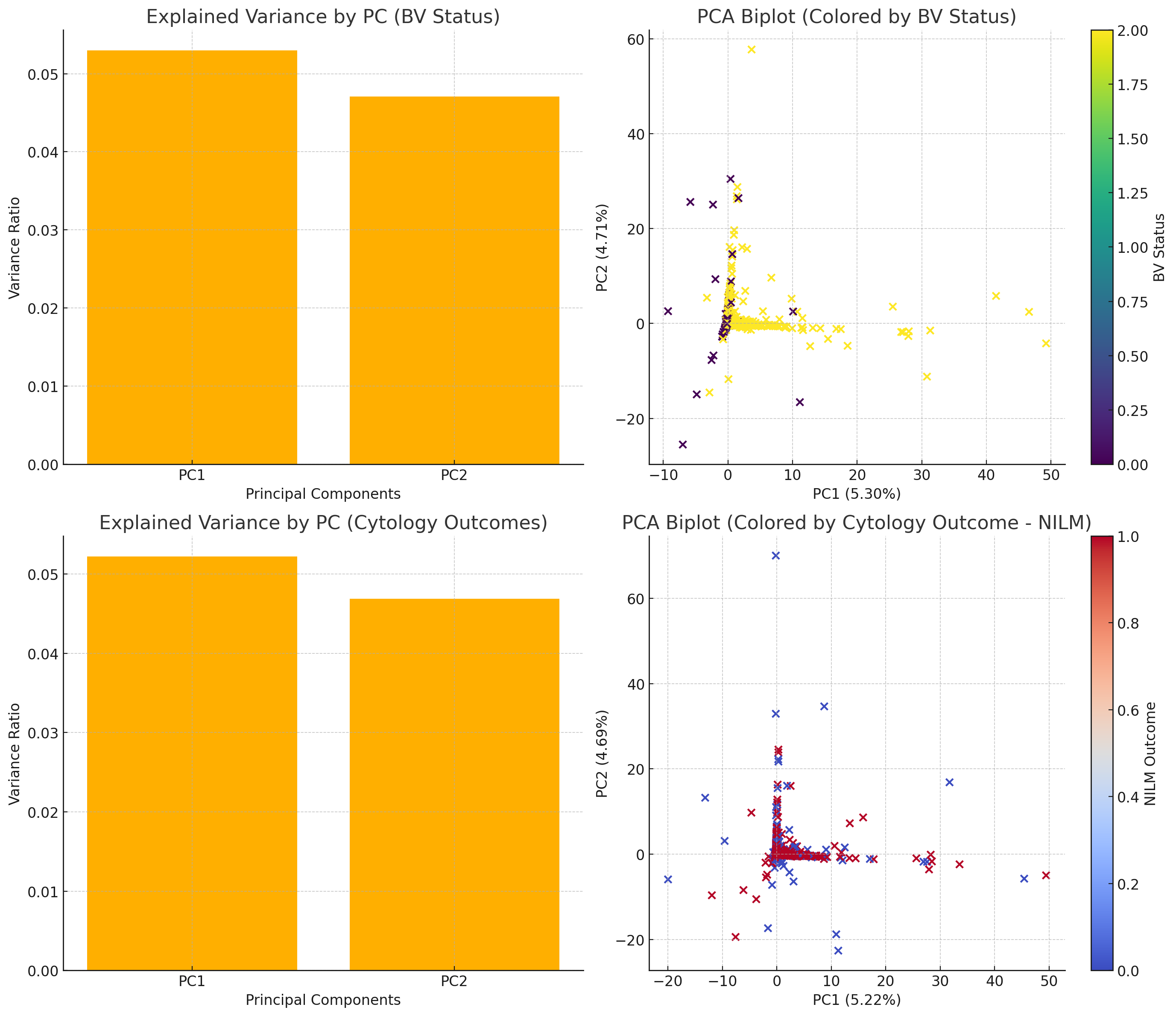

The 2x2 grid displays:

1. **Top Left**: Explained variance by the first two principal components (PC1 and PC2) for the **BV Status** analysis.
2. **Top Right**: PCA biplot colored by **BV Status**, showing how the samples separate based on bacterial species and HPV subtypes.
3. **Bottom Left**: Explained variance for **Cytology Outcomes** (using NILM as an example).
4. **Bottom Right**: PCA biplot colored by **NILM** outcome, illustrating how the data clusters based on this cervical cytology result.

This combined visualization helps in comparing how BV status and cervical cytology outcomes are influenced by the bacterial species and HPV types.

**Top Right (PCA Biplot for BV Status)**:

- The color bar on the right indicates **BV Status**, where:
  - **0 (dark purple)** = BV Negative
  - **1 (greenish-yellow)** = Transitional BV
  - **2 (yellow)** = BV Positive

**Bottom Right (PCA Biplot for NILM Outcome)**:

- The color bar on the right indicates the **NILM Cytology Outcome**, where:
  - **0 (blue)** = Normal Cytology (NILM)
  - **1 (red)** = Abnormal Cytology

**PCA Results Summary:**

1. **Explained Variance:**
   - The first two principal components (PC1 and PC2) explain the most variance in the data:
     - **PC1** explains ~66.5% of the variance.
     - **PC2** explains ~13.6% of the variance.
   - These two components together capture approximately **80.1%** of the total variability in the bacterial and HPV data.
2. **PCA Biplot:**
   - The biplot shows how different data points (samples) are spread across the first two principal components.
   - The **BV status** is used for color coding, which helps us observe potential patterns of separation between **BV positive, negative, and transitional cases**.
   - Some clustering is visible based on BV status, indicating that the bacterial species and HPV types likely have an association with BV categories.
3. **Loadings:**
   - The loadings of **PC1** are dominated by bacterial species such as ***F. vaginae****,* ***G. vaginalis***, and **BVAB-2**, suggesting that these bacterial species contribute heavily to the variance along PC1.
   - **L. iners** has a significant loading on **PC2**, meaning it influences variability in a different direction compared to other species.
   - HPV types, such as **HPV 59** and **HPV 45**, also have notable loadings on PC2, indicating their contribution to variance along that component.

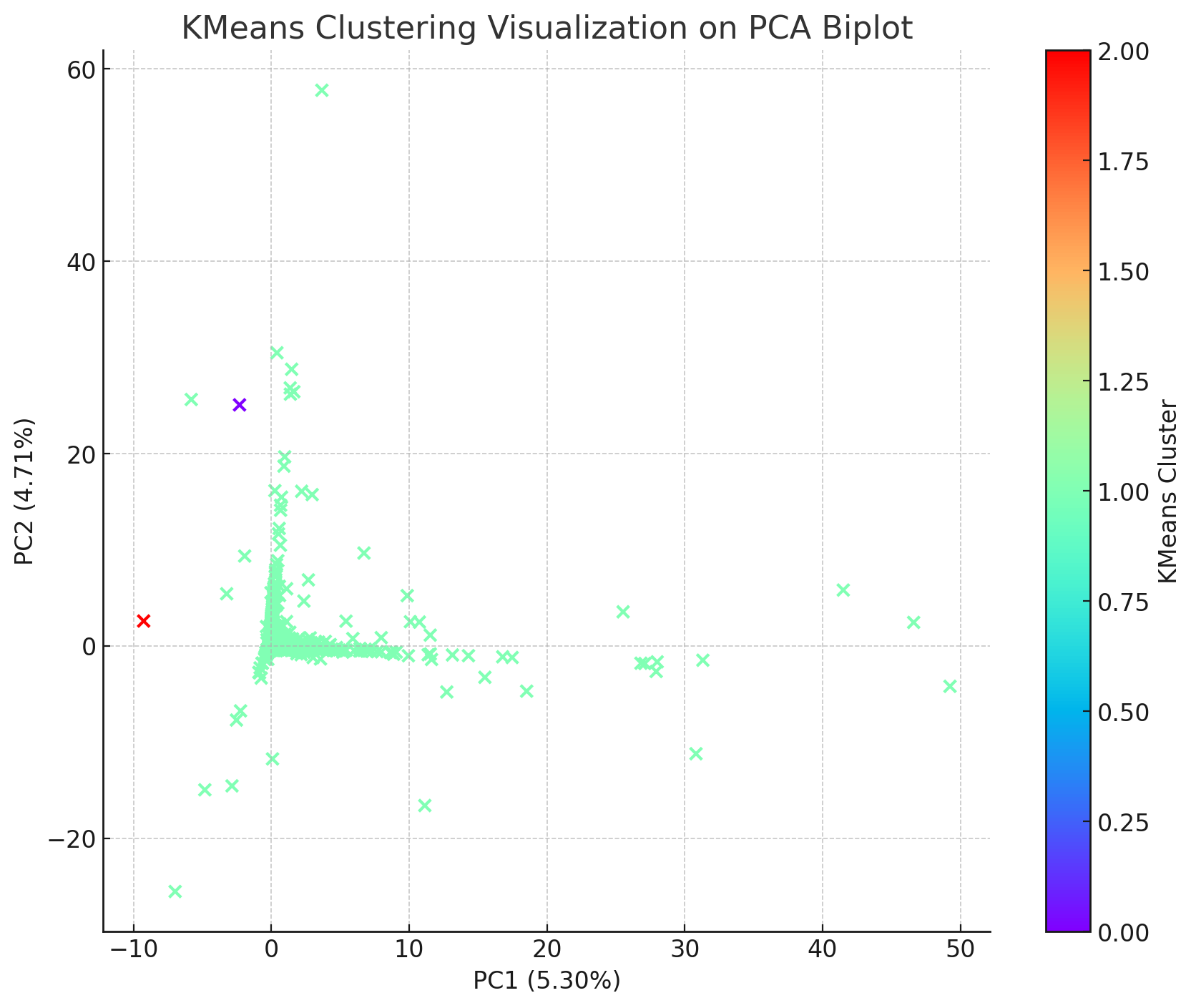

Visualization for the **K-means clustering**, projected onto the **PCA biplot**. The clusters are color-coded to show how the data points group based on the K-means algorithm, with each cluster represented by a different color.

**K-means Clustering on PCA Biplot:**

- The **scatter plot** shows data points projected onto the first two **principal components** from the PCA analysis.
- **Colors** represent the **K-means clusters**, with each color corresponding to a different cluster identified by the algorithm.
- The PCA plot helps visualize how the bacterial species and HPV subtypes group together in these clusters.

**K-means Clustering Insights:**

1. **Grouping of Data Points**:
   - The K-means clustering algorithm grouped the data into **three clusters**, with each color representing a different cluster (red, purple, and green).
   - These clusters are based on the similarity between the bacterial species and HPV subtypes across the dataset. The algorithm assigns data points to clusters in such a way that the distance between points within the same cluster is minimized, and the distance between different clusters is maximized.
2. **Cluster Meaning**:
   - Each cluster could represent different microbiological profiles or associations with certain **BV statuses** or **cytology outcomes**.
   - For example, one cluster may represent data points where the presence of certain bacterial species (like ***L. iners*** or ***G. vaginalis***) is more prominent, while another cluster could be influenced by different bacterial species or HPV subtypes.
3. **Clusters with Few Data Points**:
   - The red and purple clusters have fewer data points. These points may represent **outliers** or cases with a unique bacterial or HPV subtype profile that doesn't fit well with the majority of the samples.

**Differences and Similarities with PCA:**

- **PCA** is primarily used to **reduce dimensionality** by finding directions (principal components) that capture the most variance in the data. It doesn't group the data into clusters, but it provides a way to visualize which combinations of variables (bacterial species and HPV types) explain the most variation.
- **K-means Clustering**, on the other hand, **explicitly groups the data** based on similarity. In this case, the PCA plot is used as a **visualization tool** to show how the K-means clustering groups the data points along the two principal components.
- **Similarity**:
  - Both PCA and K-means provide insights into how the bacterial species and HPV subtypes are distributed across the samples, but K-means adds a layer of grouping, whereas PCA focuses more on variance explanation.
- **Difference**:
  - PCA explains the **overall variance** in the dataset without enforcing any grouping. It highlights which bacterial species or HPV subtypes are driving most of the differences between samples.
  - K-means actively **creates clusters**, attempting to classify data points into distinct groups. This can reveal patterns that PCA might not directly show, such as **distinct microbiological profiles** among the clusters.

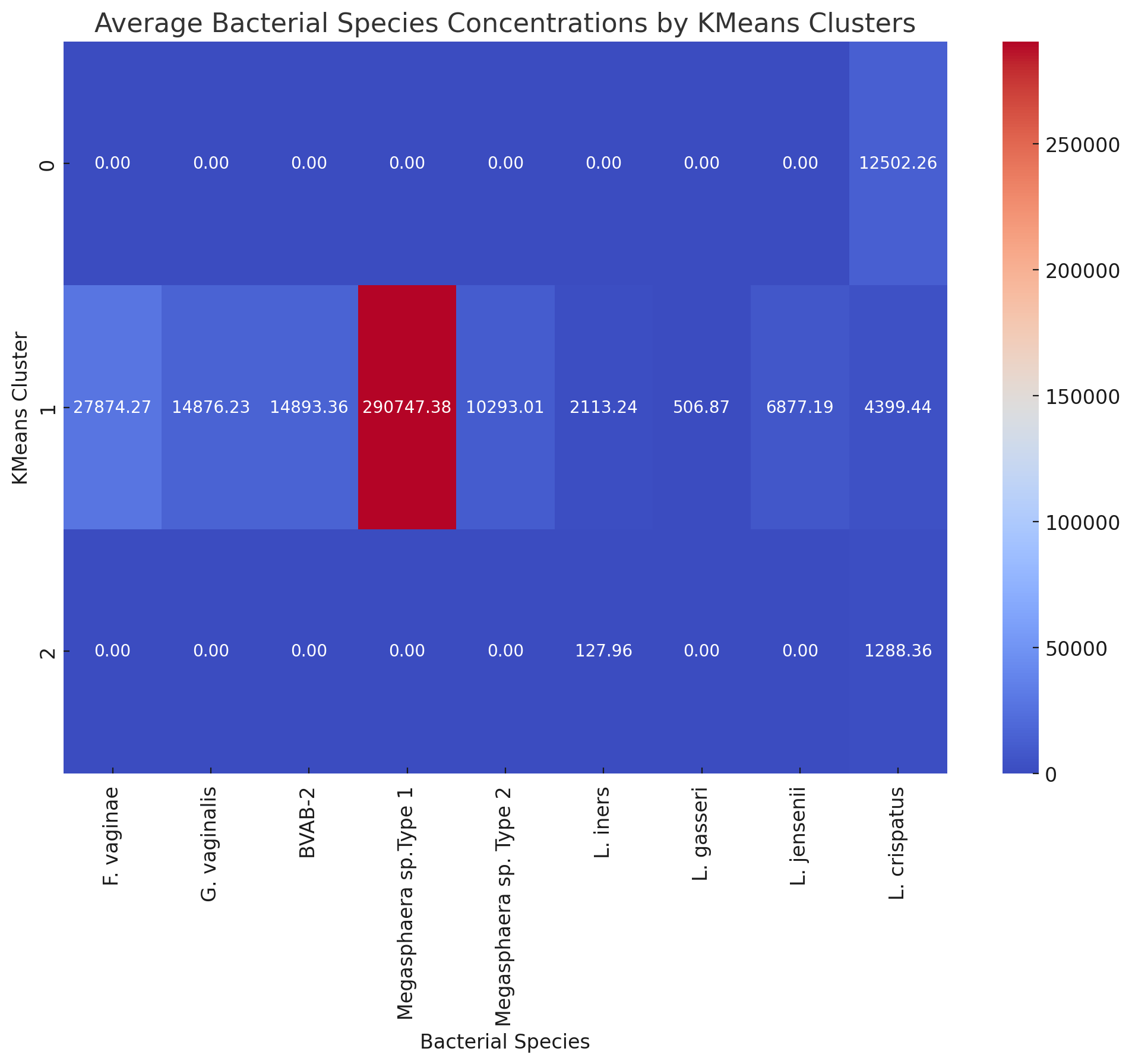

**Bacterial Species Composition by K-means Clusters:**

The heatmap and table show the **average concentration of bacterial species** in each cluster:

1. **Cluster 0**:
   - This cluster has **no significant concentrations** of any bacterial species or HPV types. It might represent samples with minimal bacterial influence or very low concentrations of these species.
2. **Cluster 1**:
   - This cluster has the **highest average concentrations** across multiple bacterial species:
     - ***F. vaginae****,* ***G. vaginalis***, and **BVAB-2** have strong presences, which are known markers of **bacterial vaginosis**.
     - ***Megasphaera sp.* Type 1** and ***L. iners*** also show higher concentrations, contributing to potential transitional BV or dysbiosis.
   - This cluster is likely associated with **BV-positive** or **transitional cases** due to the high bacterial concentrations.
3. **Cluster 2**:
   - This cluster has elevated concentrations of ***L. iners****,* ***L. crispatus****,* and ***L. jensenii***, which are typically associated with **a healthy vaginal microbiome**.
   - These species are known for their protective role in the vagina, and this cluster could represent **BV-negative** or healthy cases.

**Insights:**

- **Cluster 1** likely represents **dysbiotic samples**, possibly linked to BV-positive status, as it contains high concentrations of BV-associated bacteria.
- **Cluster 2** may be representative of a **healthy microbiome** with protective Lactobacillus species.
- **Cluster 0** might include samples with very low bacterial concentrations or possibly outliers with no significant bacterial presence.

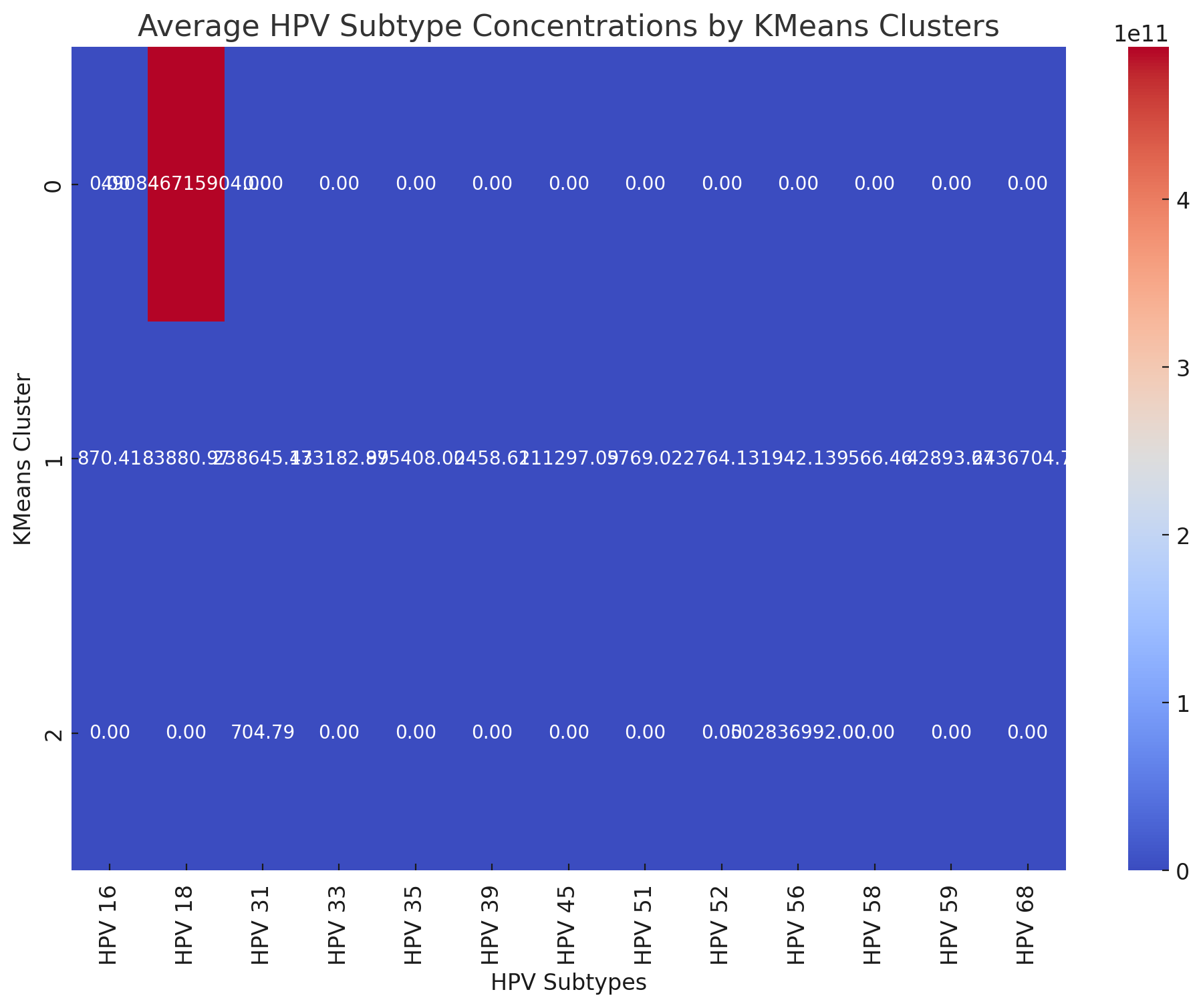

**HPV Subtype Composition by K-means Clusters:**

The heatmap and table show the **average concentration of HPV subtypes** in each cluster:

1. **Cluster 0**:
   - This cluster shows **no significant HPV concentrations** across the majority of subtypes, similar to its lack of bacterial species. It may represent samples with minimal or no HPV presence.
2. **Cluster 1**:
   - This cluster is **dominated by multiple high-risk HPV subtypes**:
     - **HPV 16**, **HPV 35**, **HPV 45**, **HPV 59**, and **HPV 68** show strong concentrations.
     - Other subtypes such as **HPV 18**, **HPV 31**, and **HPV 33** also have considerable representation.
   - This cluster likely contains **high-risk HPV infections**, which are often associated with abnormal cervical cytology outcomes such as **HSIL or ASCUS**.
3. **Cluster 2**:
   - This cluster has very low HPV concentrations, with the exception of **HPV 31** and **HPV 56**, though these are still quite minimal compared to Cluster 1.
   - These low HPV concentrations suggest that this cluster is likely associated with **normal cytology (NILM)** or **HPV-negative** cases.

**Insights:**

- **Cluster 1** is clearly influenced by **high concentrations of high-risk HPV subtypes**, suggesting that it may be associated with **abnormal cervical cytology outcomes**.
- **Cluster 2** has minimal HPV presence, potentially representing **HPV-negative samples** or **normal cytology outcomes**.
- **Cluster 0**, similar to its bacterial species composition, lacks significant HPV presence, suggesting this cluster contains **outliers** or **samples with minimal HPV or bacterial activity**.

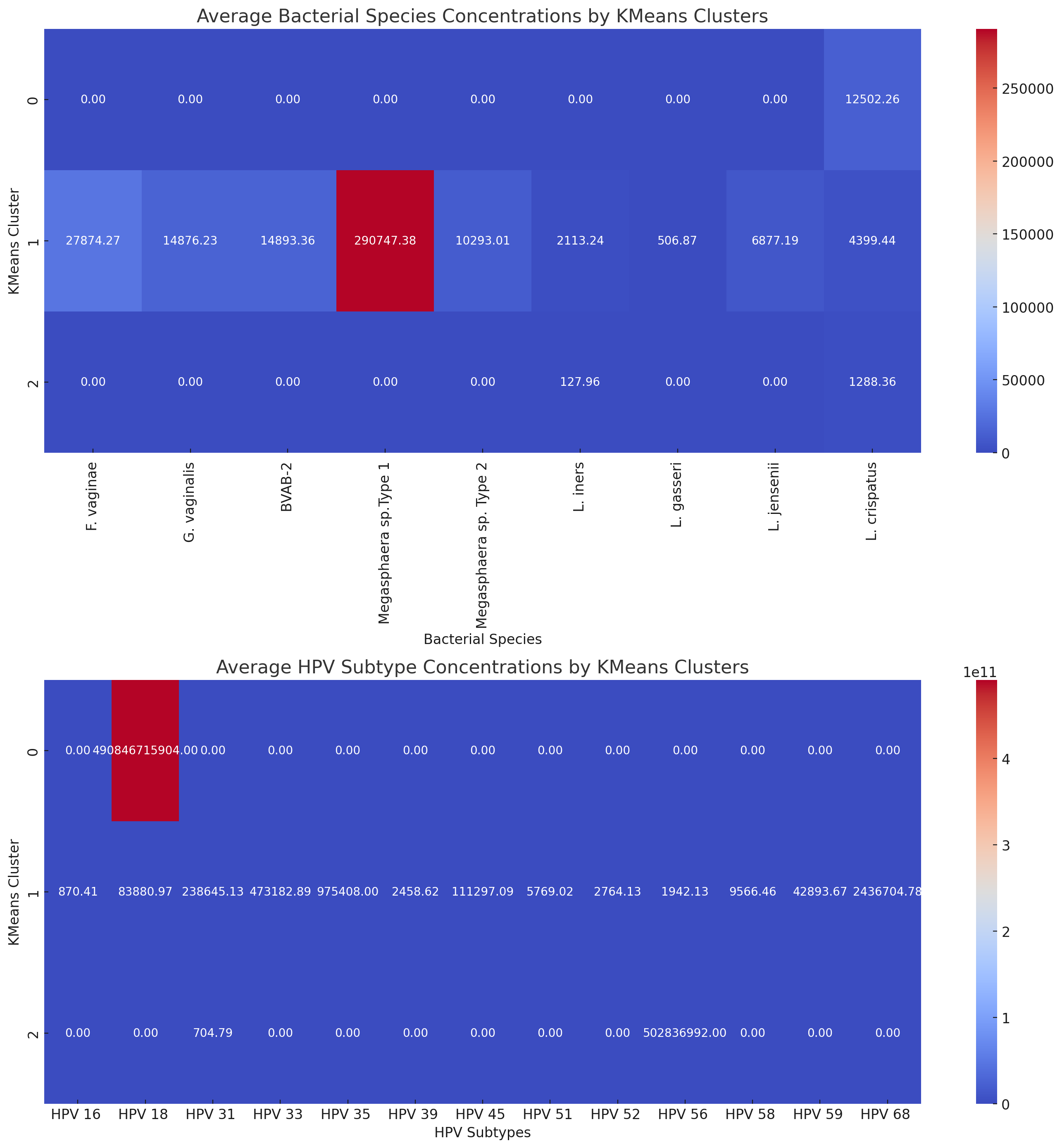

**Analysis of the Heatmaps (Bacterial Species and HPV Subtypes) and K-means Clustering:**

These combined heatmaps illustrate the **average concentrations of bacterial species** and **HPV subtypes** across the **three K-means clusters**.

**1. Cluster 0: Low Bacterial and HPV Presence**

- **Bacterial species**: Cluster 0 shows minimal concentrations across all bacterial species, except for moderate levels of **F. vaginae** and **L. crispatus**.
- **HPV subtypes**: Similarly, HPV subtype concentrations are generally very low in this cluster, with a notable exception for **HPV 16**, which shows some presence.

**Interpretation:**

- Cluster 0 could represent **low-risk cases**, with minimal bacterial dysbiosis and **very few high-risk HPV infections**.
- It might correspond to **BV-negative** or **healthy microbiome** samples where bacterial concentrations and HPV activity are not driving significant dysregulation.

**2. Cluster 1: High Bacterial and HPV Concentrations**

- **Bacterial species**: Cluster 1 is marked by **very high concentrations** of **Megasphaera sp. Type 1**, along with elevated levels of **F. vaginae**, **G. vaginalis**, and **BVAB-2**—all of which are known contributors to **bacterial vaginosis (BV)**.
- **HPV subtypes**: This cluster also shows high concentrations of **high-risk HPV subtypes** such as **HPV 16**, **HPV 35**, **HPV 45**, and **HPV 59**.

**Interpretation:**

- Cluster 1 likely represents **dysbiotic samples** with high bacterial loads indicative of **BV-positive status**.
- The **co-occurrence of high-risk HPV subtypes** suggests this cluster may also correlate with **abnormal cytology outcomes** (e.g., HSIL or ASCUS), as these HPV subtypes are often linked to cervical dysplasia and precancerous conditions.

**3. Cluster 2: Moderate Bacterial Presence and Specific HPV Subtypes**

- **Bacterial species**: While most bacterial species concentrations are low, **L. iners** has a moderate presence in Cluster 2. **L. iners** can be part of a healthy microbiome but is also seen in **transitional BV** cases.
- **HPV subtypes**: In terms of HPV subtypes, **HPV 31** and **HPV 45** show higher concentrations here compared to other clusters.

**Interpretation:**

- Cluster 2 may represent **transitional states**, possibly involving **intermediate BV** cases or **mild dysbiosis**, where some Lactobacillus species are still present, but certain HPV subtypes are emerging.
- The presence of **HPV 31** and **HPV 45** indicates potential associations with **mild cytology abnormalities** or a **pre-cancerous state**, although less severe than Cluster 1.

**Comparing with PCA Results:**

- **PCA** helped us understand the overall variability in the data, highlighting which bacterial species and HPV subtypes contributed the most to variance.
- **K-means clustering**, however, specifically groups samples into clusters based on similarity, helping to identify distinct **microbiological profiles** that may correlate with clinical outcomes like **BV status** and **cytology results**.
- In PCA, **PC1 and PC2** captured the major variances driven by both **bacterial species** (like **F. vaginae** and **BVAB-2**) and **HPV subtypes** (like **HPV 16** and **HPV 59**). These variances align with the clusters observed in the K-means results, particularly in **Cluster 1**, where high-risk pathogens and HPV types are prevalent.

**Conclusion:**

- **Cluster 1** is the most concerning, as it likely represents **BV-positive cases** with high bacterial dysbiosis and **high-risk HPV infections**, potentially linked to abnormal cytology outcomes.
- **Cluster 2** appears to be a **transitional state**, with moderate presence of **Lactobacillus** and some high-risk HPV subtypes.
- **Cluster 0** is the most benign, likely representing **healthy microbiomes** with minimal bacterial or HPV presence.

**2.4 Detailed Machine Learning Models Analyses Results**

The effect of the demographics, bacterial species, and HPV subtypes features on cervical cytology and BV outcomes were further determined using four Machine-Learning models: XGBoost, Random Forest, Decision Tree, and Logistic Regression. Based on their precision, recall, and F1 metrics, XGBoost and Random Forest were the strongest performers across both outcomes (BV and Cervical Cytology), particularly for the more common classes. Decision Tree performed comparably but slightly less effectively than XGBoost and Random Forest and Logistic Regression was the weakest model, particularly when dealing with more complex or rare classifications.

**1. BV Status Classification:**

- **XGBoost**:
  - **Important Features**: The most important predictors include *Provider State*, *HPV 56*, and other state-related features.
  - **Performance**: XGBoost demonstrated good precision and recall for distinguishing between **BV Positive** and **BV Negative**, though performance for **Transitional BV** was lower.
- **Random Forest**:
  - **Important Features**: Highlighted the importance of *Lactobacillus crispatus*, *F. vaginae*, and *G. vaginalis*.
  - **Performance**: Random Forest showed high accuracy for **BV Negative** and **BV Positive** but had moderate difficulty with **Transitional BV**.
- **Decision Tree**:
  - **Important Features**: Emphasized *L. crispatus*, *L. jensenii*, and *L. gasseri*.
  - **Performance**: While performing well for major classes like **BV Negative**, it showed variability in handling the less common class **Transitional BV**.
- **Logistic Regression**:
  - **Important Features**: Prioritized features like *L. crispatus*, *F. vaginae*, and *BVAB-2*.
  - **Performance**: Logistic Regression had difficulty with complex classes like **Transitional BV**, offering lower accuracy compared to tree-based models.

**2. Cervical Cytology Classification**

- **XGBoost**:
  - **Important Features**: The most important features included *Provider State* and some HPV-related markers.
  - **Performance**: Performed well for the **NILM** class but struggled with rarer classes like **ASC-H** and **HSIL**.
- **Random Forest**:
  - **Important Features**: Similar to XGBoost, key features were *Pt. Age*, *L. iners*, and *L. crispatus*.
  - **Performance**: Random Forest had good accuracy for **NILM** but performed moderately for more complex cytology results.
- **Decision Tree**:
  - **Important Features**: Gave importance to *Pt. Age*, *hrHPV Result*, and *L. iners*.
  - **Performance**: Decision Tree performed well for **NILM** but had challenges with rarer outcomes.
- **Logistic Regression**:
  - **Important Features**: Focused on *HPV 39*, *hrHPV Results*, and various state-level features.
  - **Performance**: Similar to BV Status, Logistic Regression had the weakest performance, particularly in handling complex cytology outcomes.

**Conclusion:**

- **XGBoost** and **Random Forest** are the strongest performers across both outcomes (BV and Cervical Cytology), particularly for the more common classes.
- **Decision Tree** performs comparably but slightly less effectively than XGBoost and Random Forest.
- **Logistic Regression** is the weakest model, particularly when dealing with more complex or rare classifications.
